# Deciphering the 3D genome organization across species from Hi-C data

**DOI:** 10.1101/2024.11.14.623548

**Authors:** Aleksei Shkolikov, Aleksandra Galitsyna, Mikhail Gelfand

## Abstract

Three-dimensional genome organization is essential for gene regulation, yet in various species it is driven by different biological mechanisms. Species-specific factors and DNA sequences influence chromatin folding, complicating cross-species comparisons. Leveraging Hi-C data and machine learning, we introduce Chimaera — a convolutional neural network that predicts Hi-C maps from DNA sequences, enabling exploration of genome folding in evolution. Chimaera’s latent representations revealed an unsupervised atlas of key chromatin features (such as insulation, loops, fountains/jets) and supported the detection and quantification of structural signatures in processes such as the cell cycle and embryogenesis. Targeted search in the latent space linked DNA sequence elements to specific chromatin structures. Applying Chimaera across multiple species confirmed the insulator roles of CTCF in vertebrates and BEAF-32 in *D. melanogaster* and identified a previously unreported insulator motif in *D. melanogaster*. In amoeba *D. discoideum*, gene orientation on the DNA strand was shown to influence loop formation. Models for other organisms also showed chromatin folding patterns associated with gene location. Finally, using cross-species predictions we tested the transferability of chromatin folding patterns and revealed evolutionary relationships, culminating in a chromatin structure-based cluster tree spanning plants to mammals.

**GRAPHICAL ABSTRACT:** 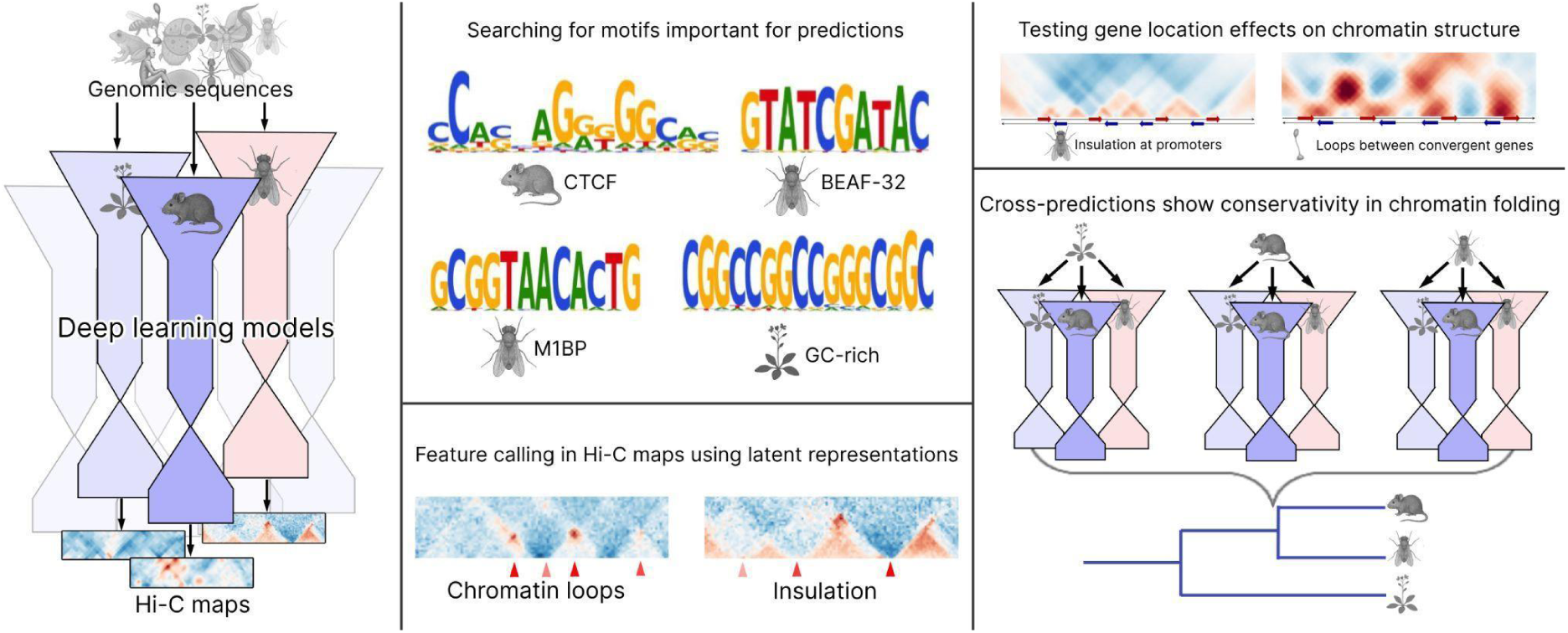

## INTRODUCTION

Three-dimensional genome (3D) organization plays an indispensable role in the regulation of gene expression and function. With the growing volume of Hi-C/Micro-C data for model and non-model species, we are gaining insights into diverse mechanisms that govern formation of 3D genome patterns.

One of the best known mechanisms of genome organization is loop extrusion (1), which folds DNA and facilitates promoter-enhancer communication in higher eukaryotes (2–5). The key player of the extrusion process is the SMC motor that loads onto DNA and reels in the DNA loop (6) until it either unloads from DNA or stalls at a barrier element (such as CTCF in vertebrates (7)). Numerous SMC motors operate in the nucleus at the same time: cohesins (8), condensins (6) and others like Smc5/6 ((9), see (10) for the review). Loop extrusion generates a variety of patterns in 3D genome interaction maps (see Suppl. Fig. 1A): (i) on-diagonal, block-enriched chromatin interactions on Hi-C maps, known as *topologically associating domains (TADs)* (1, 11), (ii) *depletion* of interactions between genomic regions separated by a boundary, known as *insulation*, (iii) dot-like enriched interactions of locus pairs, or *loops* (1, 12), enriched interactions of a single genomic locus with adjacent regions, or *stripes* (1, 13), and (iv) enriched interactions emanating from a single on-diagonal locus that spread across a characteristic distance, or *fountains or jets* (14, 15). Although the disruption of TAD boundaries has been linked to severe disorders (16), some of these patterns may simply result from mechanisms that form them without having a specific function (17).

In evolutionary terms, Hi-C visual patterns can appear similar across species: TADs were first discovered in mammals (11), then observed in *D. melanogaster* (18, 19) and *Arabidopsis* (20), and later found in other species (21). Loops have been consistently detected across multiple species, including vertebrates (12), insects, nematodes (22), and recently more broadly across the tree of life in Cnidaria, Placozoa, and Ctenophora (23). Fountains/jets have been first identified in certain mouse cell types and conditions (14, 15, 24), during zygotic genome activation in vertebrates (zebrafish, frog *X. tropicalis*, and medaka) (14), nematode *C. elegans* (25, 26), and in more distant species such as fungi (27) and plant *Arabidopsis thaliana* (28).

Visual similarity of Hi-C/Micro-C patterns between species may not reflect the conservation of biological mechanisms underlying the formation of these patterns. For example, TAD formation in mammals and in insect *D. melanogaster* has been attributed to two very different mechanisms. In mammals, individual nuclei are folded by a stochastic process of loop extrusion that may stall at CTCF barriers, forming averaged bulk Hi-C patterns of TAD boundaries, bright dot-like enriched interactions (loops), and stripes, usually invisible in individual cells (1, 12). In contrast, *D. melanogaster* TADs are very prominent in individual cells (29), usually lack corner peaks (30), and may be formed mainly due to histone modifications (29, 31). The role of loop extrusion in *D. melanogaster* remains debated (*29, 32, 33*), suggesting it may not function in the same regulatory capacity as it does in mammals.

A complementary feature of TADs is the avoidance of interactions across specific genomic regions, also known as TAD boundaries or areas of insulation (Suppl. Fig. 1A). In mammals, CTCF is a key extrusion-dependent insulator, a protein causing insulation at its binding site (12). In *D. melanogaster*, multiple proteins can be found at insulating boundaries, including BEAF-32, Su(Hw), CTCF and others (19, 34, 35). Recent studies suggest that genes themselves can serve as insulating elements (36, 37), even without loop extrusion in yeast (38).

Thus, 3D genome organization is species-specific in a broad sense, meaning that the rules governing chromatin folding can differ significantly over large evolutionary distances. More narrowly, 3D genome organization is species-specific due to variations in DNA sequence and the positioning and properties of architectural factor binding sites. For example, CTCF sites are often conserved throughout mammalian evolution (39) and deuterostomes more generally (40), leading to conserved TAD organization in syntenic regions. The evolutionary stability of TADs has been linked to conserved gene regulation, with well-studied examples including *Hox* loci (41), *Six* homeobox genes (40), and globin genes (42).

Evolutionary studies of the 3D genome organisation usually focus on a single source of variability between species: (i) evolution of chromatin factors or (ii) evolution of DNA sequence. The studies of the first type investigate species-specific chromatin factors across the nucleus and analyse general 3D genome patterns from Hi-C maps (21, 43). The studies of the second type assume conserved folding mechanisms but focus on conserved, syntenic regions of genomes (39, 44–46), or even specific conserved genes (40, 42, 47).

A major breakthrough in the field came with the development of neural networks capable of predicting chromatin structure from DNA sequences (48–50). These models learn DNA folding rules and can be tested on unseen sequences, such as rearranged cancer DNA (49). This approach enabled a unique experiment never performed *in vivo*: visualizing the chromatin folding of one species’ DNA within another species’ cellular environment, as simulated by the neural network Akita (48). Interpreting these cross-species predictions revealed significant differences in chromatin folding mechanisms between mouse and human: CTCF binding at B2 SINE elements in mice is hindered, resulting in a loss of insulation at these CTCF sites, but not in human (48).

More broadly, cross-species prediction is a powerful tool that enhances the performance of genomic models (51), allows for the creation of unified gene regulation models for species like mouse and human (52–54) and enables tracking the evolution of regulatory elements along hundreds of millions years of evolution (55). Furthermore, this approach is crucial for imputing missing data, such as mammalian methylation patterns (56) and single-cell expression profiles (57). It also facilitates functional annotation of genomes by identifying promoters (58), predicting cell types from single-cell Hi-C data (59), and transferring knowledge of features like translation and splice sites to less studied plant genomes (60). Ultimately, these combined capabilities are foundational to ambitious projects like building a "tree of life" for cell types from single-cell RNA-Seq data (61).

In this study, we leverage the wealth of existing Hi-C and Micro-C datasets from diverse species, ranging from plants to mammals, and build upon the idea of interpretable machine learning to predict DNA folding patterns from DNA sequence. We introduce *Chimaera*, a neural network-based tool for predicting species-specific 3D genome organization. Chimaera’s autoencoder-based architecture (62) combined with a two-step training process that first learns Hi-C patterns and then links them to DNA sequence, enables its application to various tasks in chromatin biology. These tasks include predicting 3D structures from DNA sequences, searching for and quantifying Hi-C patterns, and interpreting associations between DNA sequences and 3D genome patterns. This work presents the first case study of species-specific 3D genome folding mechanisms analysed through neural networks across a wide variety of organisms, shedding light on the conservation and variability of chromatin folding in evolution, and paving the way for future advancements in understanding 3D genome evolution and distal gene regulation.

## MATERIAL AND METHODS

### Data collection

In total, we collected data for 22 organisms (Suppl. Table 1): Micro-C for *Homo sapiens* HFFc6 cell line from (63), Micro-C for *Mus musculus* embryonic stem cells from (64), Micro-C data with depletion of structural proteins from the same study (64), for cell cycle of erythroid cells G1E-ER4 from (65), Hi-C for *Mus musculus* in conventional dendritic cells from (5), Hi-C for *Xenopus tropicalis* embryos from (66), Hi-C for *Danio rerio*: for embryos at 5.3 hours past fertilization from (14) and for muscle cells from (67), Hi-C for gastropods *Pomacea canaliculata* from (68) and *Arion vulgaris* from (69), Hi-C for bee *Apis cerana* from the drone pupae from (70), Hi-C for ant *Cataglyphis hispanica* from (71), Hi-C for silk moth *Bombyx mori* from (72), Micro-C for fruitfly *Drosophila melanogaster* for nc14 stage embryos from (73), Hi-C for mosquitoes *Anopheles merus* from (45) and *Culex quinquefasciatus* from (74), Hi-C for mites *Sarcoptes scabiei* from (75) and *Archegozetes longisetosus* from (76), Hi-C for nematode *Caenorhabditis elegans* from (77), Micro-C for comb jelly *Mnemiopsis leidyi* and placozoan *Trichoplax adhaerens* from (23), Hi-C for social amoeba *Dictyostelium discoideum* from (37), Micro-C XL for yeast *Saccharomyces cerevisiae* at 90 minutes after release from G1 from (78) and cells on G2/M stage with exogenous bacterial DNA from (79), Micro-C for fission yeast *Schizosaccharomyces pombe* from (80), Hi-C for dinoflagellate *Symbiodinium microadriaticum* from (81), and Micro-C for plant *Arabidopsis thaliana* seven-day-old seedlings from (82).

Although we considered more candidate species for our analysis, we had to exclude them for various reasons. For example, archaeon *Haloferax volcanii*, known for its unique chromatin organization (83), had a very small genome relative to a meaningful resolution of Hi-C map, resulting in less than 100 windows for training, not sufficient for our model.

When the original studies did not provide iteratively corrected data in .cool format with the desired resolution, we re-mapped the data using distiller-nf pipeline (84) that relies on pairtools (85) and cooler (86) libraries for Hi-C data processing.

### Hi-C data preparation

The following pipeline is illustrated in Suppl. Fig. 1B.

#### Hi-C snipping

We extracted specific regions, or *snippets*, from Hi-C interaction matrices for the model input. This approach provided focused sections that emphasized local chromatin interactions, optimizing computational efficiency in downstream processing. Snippet positions were determined by the model training scheme, as described below: the map was segmented into constant-sized fragments (W, see Suppl. Fig. 1B, also Suppl. Table 1), shifted along the genome with a step equal to half the fragment size to guarantee that individual samples are non-overlapping. We excluded contacts from the first two diagonals (below two dataset resolutions) of Hi-C data as potential sources of self-circles, dangling ends, mirror reads (87), and other short-range artifacts (88, 89).

#### Snippet size definition

The Hi-C snippet size is a key hyperparameter because it sets the model’s receptive field — the linear genomic span covered by an input Hi-C snippet (window "W" in Suppl. Fig. 1B/1). Larger windows reduce the size of a training sample; therefore, we choose the smallest window that still captures target Hi-C/Micro-C features — ideally 2 – 3 times their characteristic scale and not smaller than that scale. The snippet height (on the contact-distance axis) is set to W/2, and is referred to as "the receptive distance" throughout.

#### Iterative correction

When snipping, we obtained Hi-C/Micro-C contact frequencies normalized by iterative correction (89), which (i) removed bin-specific mappability, amplification and other biases, (ii) normalized interactions and allowed interpretation as contact frequencies. If the source studies did not provide iteratively corrected Hi-C/Micro-C maps, we performed iterative correction using the default parameters.

#### Log-transformation

To further stabilize the variance of contact frequencies, we applied a log transformation. This adjustment reduces the impact of high-frequency outliers and makes the distribution of contact frequencies more amenable to neural network training. Note that we aimed to avoid obtaining non-interpretable values for Hi-C elements with zero contacts, and added a pseudocount of 10^−3^ to all elements of the matrix prior to the log-transformation.

#### Observed-over-expected normalization

To account for the tendency of contact frequency to decrease sharply with increased genomic distance, we normalized observed log-transformed Hi-C contact interactions by subtracting the expected interactions for each genomic separation distance. Expected interactions were calculated as mean log-transformed average values for each genomic separation for each chromosome. This observed-over-expected transformation helps reveal local interaction patterns that may otherwise be overshadowed by distance-dependent effects.

#### Normalization by standard deviation

Next, we divided all values by the standard deviation of the signal for each chromosome.

#### Treating unmapped regions

We used two approaches to treat missing data in the input Hi-C maps. First, we detected unmapped bins as those that failed to be iteratively corrected by *cooler* (86) with default parameters. Then, we interpolated those bins by *cooltools*-based bilinear interpolation (90). This approach was used for training the Hi-C autoencoder. Alternatively, for the training that requires more rigor (such as DNA encoder), we have excluded the unmapped bins from training and evaluation to avoid learning Hi-C artifacts instead of the true interaction signal. For that, we have dynamically modified the loss function (see below) and zeroed out unmappable regions of the Hi-C maps. Hi-C/Micro-C snippets with more than 25% of unmapped bins were permanently excluded from both training and validation samples. This approach allowed us to minimize the effects of the repetitive DNA which is frequently unmappable.

#### 45-degree rotation

To minimize the influence of noisy long-range interactions, we rotated each normalized Hi-C map by 45 degrees by *scipy* (91) *ndimage.rotate* using spline interpolation of 0 order (no smoothing), effectively rearranging interaction patterns to emphasize local contacts. Then the target map fragment is cut off and resized to 128 pixels width (Suppl. Fig. 1B). This transformation (i) excludes long-range interactions beyond half the map (snippet) window size and (ii) resizes the Hi-C map to dimensions of 128×32, standardizing the inputs and outputs across all models. This approach improves the model’s focus on essential chromatin interaction structures while minimizing noise from distant genomic contacts. For datasets with worse data quality, we applied the 1-st order spline interpolation.

### DNA data preparation

To match the DNA sequence data with Hi-C interaction matrices and optimize it for model training, we performed several preparatory steps to standardize the input format and address resolution mismatches.

#### One-Hot encoding

DNA sequences were encoded in the one-hot format, where each nucleotide (A, T, C, G) is represented by a unique binary vector. Nucleotides with ambiguous base calls were replaced with "N" (zero one-hot encoded vector), ensuring that unresolved regions do not contribute to the sequence representation.

#### Choice of DNA fragment matching Hi-C map

DNA sequence was taken from the region centered at the corresponding Hi-C snippet location. The size of the DNA window was larger than those covered by Hi-C to allow the model to learn from the context of the input Hi-C interactions. The choice of the offset size was species-specific, and guided by balancing computational efficiency and predictive quality. Larger offset windows improve alignment accuracy but increase the required memory and the training time. Therefore, the final offset size was optimized to provide high predictive quality while managing memory and processing constraints effectively. The offset size was capped at no more than half the Hi-C window size, allowing for adequate coverage while limiting excessive data expansion.

### Chimaera implementation

We implemented Chimaera with Python framework pytorch (92). Chimaera consists of two parts: the Hi-C autoencoder and the DNA encoder that are trained separately (Fig. 1, Suppl. Fig. 1C).

**Figure 1.**
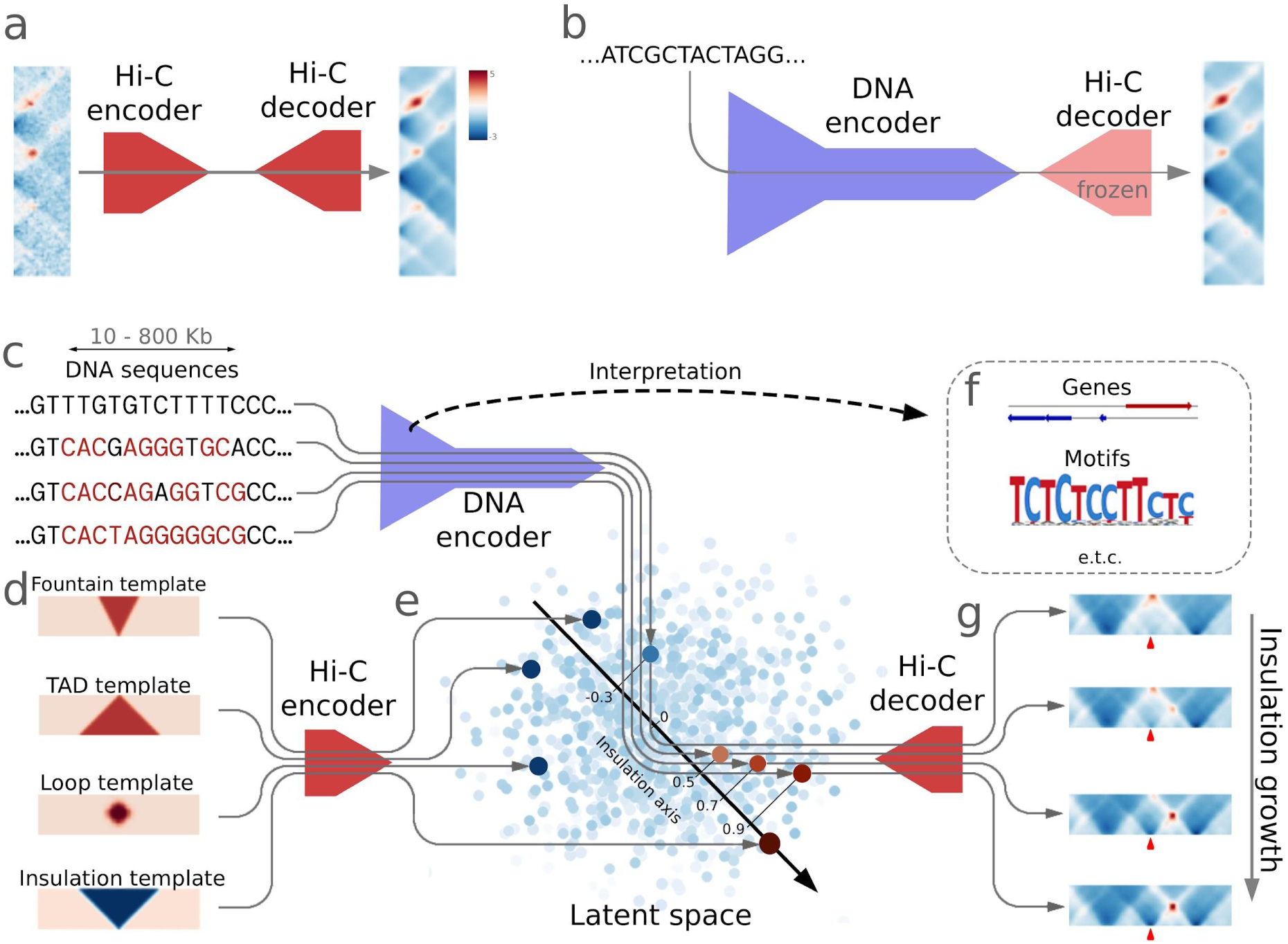
Summary of Chimaera, Convolutional neural network for Hi-C maps prediction using autoencoder for maps representation. Each grey line corresponds to the forward pass of the single input (DNA or Hi-C map) through the network. The schematics summarizes the training scheme (A, B) and three principal ways of using Chimaera in this work: Hi-C prediction from DNA sequence (C, G), pattern search and quantification (D, E), and interpretation of DNA sequence / 3D genome pattern associations (F). For the full architecture, see Suppl. Fig. 1C. Real data is shown for human. (**A**) Training strategy for the Hi-C autoencoder. The network consists of two blocks: Hi-C encoder and Hi-C decoder that are trained simultaneously to denoise Hi-C maps. (**B**) Training strategy for the DNA encoder. DNA serves as input to predict latent representation of Hi-C map for corresponding genomic region. Weights of Hi-C decoder are frozen (do not change during training). See Suppl. Table 2 for comparison of different training strategies. (**C**) Example DNA sequences serving as input to the DNA encoder. Red letters represent parts of CTCF motif instances in the genome. Top: sequence does not have any CTCF sites, bottom: sequence contains a recognizable CTCF site. (**D**) Example of patterns serving as input to the Hi-C encoder for feature calling and quantification. We use 45-degree rotated Hi-C maps, where the horizontal axis corresponds to genomic coordinates and the vertical axis reflects genomic separation. (**E**) Latent space of autoencoder and results of projection into it. Light-blue dots: embeddings of Hi-C maps for each genomic region. **From (D) to (E)**: each pattern is projected onto a vector in the latent space. Note that the TAD pattern is far from the insulation pattern, while the fountain (jet) and the loop (dot) are located somewhat in between. Note the location of the insulation vector used below for the quantification of the insulation pattern. **From (C) to (E)**: each sequence is projected onto the vector in the latent space. In this example, we project each vector onto the vector of insulation to quantify the prominence of insulation at each genomic location. The length of the projection serves as a similarity measure of genomic location to an input pattern (here, insulation). (**F**) Maps predicted by Chimaera’s Hi-C decoder based on latent representations (E) made by DNA encoder (C). Note how insulation grows in these examples, reflecting the prominence of the CTCF motif in the input sequence (C). (**G**) Model interpretation. Extracting the information about DNA determinants of the 3D genome organization, such as motifs of binding factors or gene locations.

The first component is a *Hi-C autoencoder* designed to produce both a denoised Hi-C map and its latent representation. This 2D convolutional network combines an encoder that transforms the input image into a multi-dimensional latent vector, and a decoder that reconstructs a denoised image from this latent representation. We implemented max pooling operations between the sequential layers of the network and added two fully connected layers at the transition to and from the latent space (Suppl. Fig. 1C, the horizontal component). We used 128 dimensions for most species, and 96 dimensions for species with relatively small train samples.

The second part is a *DNA encoder*, a model that converts the nucleotide sequence into a multi-dimensional vector. It is a 1D convolutional network with residual blocks. Again, we used max pooling between some convolutional layers and added a fully connected layer at the end (Suppl Fig. 1C, the vertical component). The number of dimensions of the latent space matched the number of dimensions for the Hi-C autoencoder.

Given existing tools and frameworks for deep learning on DNA sequence (93–96), our aim was to design a specialized solution optimized for the chromatin organization. We implement an API that abstracts the Chimaera model creation and training. The model requires cool-formatted Hi-C/Micro-C files as input, and DNA sequence in fasta format. The model then allows for selection of the Hi-C map resolution and the fragment size. The problem of parameter selection is simplified by the Chimaera module performing data quality control and visualization.

Next, the user can set the hyperparameters for the Hi-C and DNA encoders, including: the number and the size of convolutional filters, the number of residual blocks, the percentage of dropout, and many more. We have three ready-to-use presets that we recommend for different sizes of the genomes:

(i) “Small”, designed to be trained on small sample sizes that are prone to overfitting. It has higher dropouts, no residual blocks, and 96-dimensional latent space.

(ii) “Middle” and (iii) “Big” are both designed for larger sample sizes that are expected to result in less overfitting. The dropout rates are smaller, the latent space dimensionality is 128, and residual blocks are used (4 in “Middle” and 8 in “Big”).

Chimaera’s API implements model training, validation, visualization of results, and interpretation of the model as separate modules. The API and the examples of usage are available at GitHub link: https://github.com/ashkolikov/chimaera/

### Hi-C autoencoder training

The Chimaera workflow began with the training of a Hi-C autoencoder (Fig. 1A). Weights are initialized using the Kaiming initialization (97).

#### Model setup

As input, the model takes Hi-C/Micro-C pre-processed snippets. The loss function is the Mean Squared Error (MSE) between the original and reconstructed maps, with the Kullback–Leibler (KL) divergence regularization. The regularization penalizes the difference of the latent space representation of the Hi-C autoencoder and the standard normal distribution, which makes the encoder variational and is a prerequisite for the continuity of the latent space (98). When calculating the loss function between the true and predicted maps, we used maps with interpolated unmapped bins (see “Hi-C data preparation”).

#### Model training

The Chimaera network weights were optimized by the Adam gradient descent (99). We stopped training of the model when the plateau of the validation quality metrics was achieved.

#### Whole-genome model application

Once the Hi-C encoder was trained, we ran it on all snippets of Hi-C maps and obtained (i) latent representations and (ii) reconstructed maps across the genome of each species. We later used this information for training the encoder, model quality control, and interpretation.

#### Hi-C autoencoder transfer

Since Hi-C maps for all species are treated in the same way and have the same final size (128×32). Since many organisms (even distant ones) usually have a common set of chromatin structures, it is possible to train a Hi-C autoencoder on one species and transfer it for Hi-C maps for other species (especially if the expected sizes and the types of Hi-C features are similar). We used such autoencoder transfer when the data was not good enough to train a new autoencoder (Suppl. Table 1).

### DNA encoder training

After pre-training the Hi-C autoencoder, we fixed the weights of the Hi-C decoder and stacked it to the DNA encoder (Fig. 1B). Weights of the DNA encoder were also initialized using the Kaiming initialization (97). As input, the DNA encoder takes a one-hot encoded sequence of the reference genome. During training, we used the Adam gradient descent (99).

#### Loss function

Importantly, we set the loss function as MSE between the DNA-based predicted Hi-C map and the autoencoder-reconstructed Hi-C map for the same snippet. The rationale behind this choice is that decoded Hi-C maps after the Hi-C decoder (i) contain only the information that is possible to pass through a lower-dimensional vector, and (ii) are denoised from irrelevant random fluctuations of pixel values. We did not use an alternative with similar functionality, the Gaussian blur, because we aimed to allow our denoising method to keep the shapes of actual chromatin structures sharp, as can be achieved by an autoencoder, see Fig. 2A, B).

**Figure 2.**
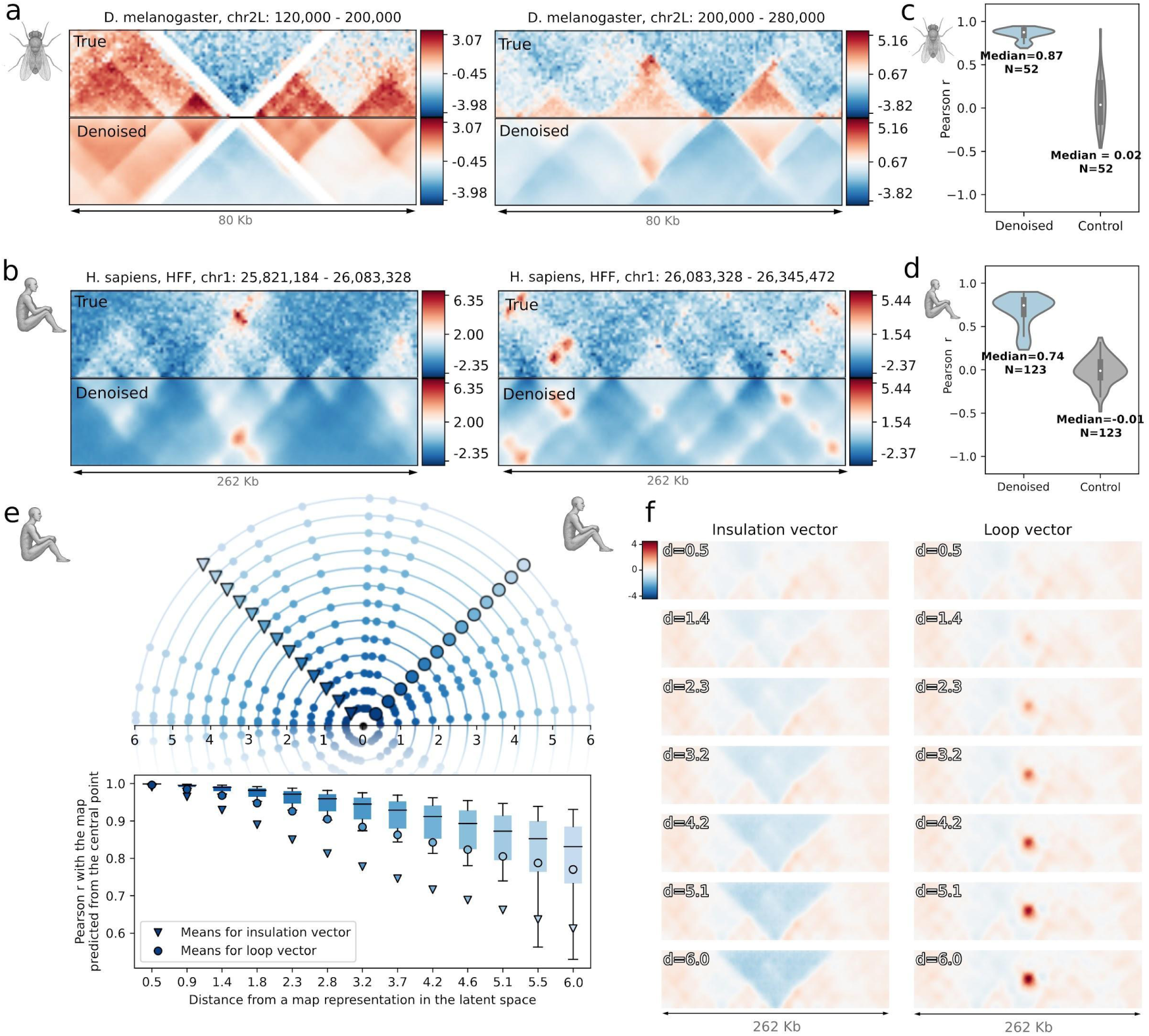
Key features of the Hi-C encoder are denoising of Hi-C maps (A-D) and generating a continuous, interpretable latent space (E-F). (**A, B**) Examples of noise reduction with the Hi-C autoencoder for maps of *D. melanogaster* (A) and *H. sapiens* (B). The top half of each plot is the actual map, and the bottom half is the mirrored predicted one. The represented values are the Z-score normalized observed-over-expected signal (a true Hi-C/Micro-C map *vs.* the map produced by Chimaera). (**C, D**) The Pearson correlation on the test set between the original maps and the autoencoder-denoised maps for *D. melanogaster* (C) and *H. sapiens* (D). Controls: randomly paired true and predicted maps. (**E**) Demonstration of latent space continuity. Top: We start with the vector representations of the real Hi-C maps in the latent space of the Hi-C encoder (zero point). Next, we step away from each point with a fixed step size in a given direction (triangle – towards insulation, circle – towards loop). At each step, we decode the maps for the resulting shifted vectors and calculate the similarity to the original map (shown by the boxplots). (**F**) Average outputs of shifts in the latent space. Left: shifts across the latent representation of the insulation pattern. Right: shifts across the latent representation of the loop pattern. *d* is the distance in the latent space to the original points, as defined in (E). Note that the shift towards positive values in the direction of the insulation pattern vector results in more pronounced insulation in the generated Hi-C maps. This observation is consistent with the fact that intact genomic regions with larger size of the projection onto the insulation vector have higher insulation scores (Suppl. Fig. 2A).

In Chimaera API, we retain an option for a user to use raw Hi-C maps as the ground truth for training. However, this option yielded worse performance (Suppl. Table 1). Note that in both cases the evaluation metrics were calculated with raw maps. For clarity and uniform interpretation of the model quality, we run validation on the original Hi-C maps, without applying the Hi-C decoding.

In contrast to Hi-C autoencoder training, when calculating the loss function between the true (decoded) and predicted maps, we masked out interpolated unmapped bins (see “Hi-C data preparation”) — for these pixels the loss function was set to 0, and for the remaining ones it was reweighted to preserve the average value. In this way, the model learned to predict only pixels obtained from experimental data, ignoring possible interpolation artifacts.

#### Data augmentation

Hi-C data, by design, is not strand-specific; the Hi-C map for the forward strand mirrors that of the reverse complement strand when rotated 180 degrees. To prevent the strand bias, we implemented a random reversal of data points (reverse Hi-C along with the corresponding reverse complement DNA). This approach also effectively doubles the data pool for augmentation purposes.

#### DNA encoder transfer

DNA fragments in our study had different sizes for different species and chromatin folding mechanisms differ between taxa. Thus, unlike the Hi-C autoencoder transfer, the DNA encoder transfer is applicable only between taxonomically close species with comparable genome sizes and needs a consequent fine-tuning. We used it to compare *H. sapiens* and *D. rerio* models (Suppl. Table 1).

### Data segmentation and splitting

#### Segmentation of chromosomes into fragments

Since the input fragments were parts of contiguous chromosomes, they could be sliced with overlaps. At the same time, feeding the same sequences into the model, but in different contexts, would help to better resist overfitting. To do this, each training epoch included several sub-epochs, after each of which the genome marking of the training sample was shifted by a small step. We chose a step of 1/12 of the fragment length, which improved the metrics on the validation sample by up to 20%. For the final prediction of the models, predictions were made for both direct and reverse sequences, and the results were averaged. Empirically, this averaging enhanced prediction accuracy across all organisms. Low-quality fragments with mappability less than 50-95% (depending on the input data quality for each species) were removed.

#### Train-test-validation splitting

Fragments for test and validation samples were taken from full-size chromosomes or contiguous parts of long chromosomes separated for this purpose. We ensured that the train and test/validation sets never contained overlapping Hi-C or DNA segments. For species with abundant high-resolution data, we reserved whole chromosomes as test/validation. For species with more limited data, we selected continuous parts of long chromosomes (to ensure at least 30 fragments) for the test/validation sample. To ensure absence of leakage between the train and test sets, we implemented an internal check function in Chimaera that ensures that no overlapping fragments happen to be both in the train and test/validation sets. The split between the test and validation was 2:1.

### Model performance evaluation

Throughout this work, we used two generic and one structure-based evaluation of Chimaera performance.

The generic metrics are not specific for Hi-C maps or local structures in them: (i) the Pearson correlation between true and predicted Hi-C snippets (most of the quality controls in the paper if not mentioned otherwise), (ii) the Pearson correlation between each row (corresponding to Hi-C contacts at a specified distance) of true and predicted windows with a subsequent selection of a row with the highest median metric (for Suppl. Table 1, 2 and Suppl. Fig. 6). This allowed us to study the model performance for different genomic separations, select characteristic distances where the model performs the best, and avoid the issue of Hi-C having worse quality and signal-to-noise ratio at higher genomic separations. To ensure there is no correlation between random pairs of map fragments, we calculated correlations between predicted maps and randomly selected maps from the dataset. Significances for each distance were calculated using the two-sided Mann-Whitney test and p-values were adjusted using the Benjamini-Hochberg procedure.

The procedure for the calculation of structure-based metrics includes obtaining latent representation-based profiles of the main structural patterns (insulation, loop, TAD, and fountain) for both true and predicted maps (see “Pattern calling in Hi-C maps”). The profiles are then treated as full maps in previous two metrics but the Spearman correlation coefficient is used instead of the Pearson correlation as it is more appropriate for possible nonlinear dependences.

#### Robust predictions for validation

For interpretation, visualization, and scoring, we have implemented robust predictions by Chimaera, when we run multiple predictions of adjacent regions and average overlaps between them for the final output. Robust predictions usually have higher quality (by ∼10%), but were not used during the model training.

### Pattern calling in Hi-C maps

#### Selection of genomic regions for pattern calling

For pattern calling, we used whole genome Hi-C maps. Here, we did not separate them into the train and test sets, as we were interested in the presence of certain patterns in the entire genome.

#### Genome scanning to evaluate the latent representation

We applied Chimaera to snippets of Hi-C maps, with the fragment size as required by the model (Suppl. Table 1) and the step equal to 8 bins in the transformed Hi-C matrix dimensions (after 45-degree rotation, see “Hi-C data preparation”). Each snippet served as an input for the Hi-C encoder, and the output was recorded.

#### Obtaining known pattern templates

First, we constructed Hi-C pattern templates for insulation, loop, and fountain. We started with an *in silico* Hi-C matrix filled with zeros. The size of the matrix was set to the size of the Chimaera input (128×32).

For the insulation pattern, we positioned a right triangle filled with negative values (based on the map values distribution), with its apex at the bottom (in genomic separation coordinates), centered in the snippet (in genomic position coordinates). The insulation in the pattern template is thus centered in the middle of the snippet (Fig. 1D).

For the loop pattern, we placed a rhombus with the diagonal size of 16 pixels at the position equal to half of the window size (in genomic separation coordinates). Note that compartmental interactions might appear as loops in Hi-C maps and can be in theory learned by the model (see the legend of Suppl. Fig. 1A).

For the fountain pattern, we positioned a triangle filled with positive values, with the apex at the bottom (in genomic separation coordinates) centered in the snippet (in genomic position coordinates). The angle at the bottom-facing apex was 45 degrees, reflecting the growth of the fountain spread with genomic separations (14).

For the TAD pattern, we positioned a right triangle filled with positive values, with its apex at the top (in genomic separation coordinates) centered in the snippet (in genomic position coordinates).

#### Obtaining the latent representation of the known pattern templates

The known pattern templates were input into the Chimaera Hi-C encoder, and their latent representations were obtained.

#### Similarity of latent representations

To measure the similarity between the latent representation of Hi-C snippets from real Hi-C maps and the templates, we projected the latent vector of the real Hi-C representation onto the latent vector of the template. This results in a number (measure of similarity) of Hi-C signal at a genomic position to the template. Combined with the whole-genome scanning, this results in a genomic track of similarity to the given pattern.

#### Thresholding of the similarity track

To obtain the list of called instances of patterns present in Hi-C maps, we applied thresholding of the similarity track. We used a two step approach to choosing the threshold:

i. *Constructing an average map adjusted for the expected pattern.* The aim of this step is to obtain the average of maps that represent the target pattern and fall above a specific threshold. However, naive averaging of all Hi-C/Micro-C maps above the threshold can have an appearance of average pattern even if maps of individual loci do not feature this pattern (see Suppl. Information of (14)). Thus, we adjusted each individual Hi-C/Micro-C map by subtracting the template, resulting in an observed over expected pattern map. This procedure eliminated the described problem.
ii. *Threshold selection*. Adjusted maps obtained at step (i) are similar to the template pattern only if the target pattern is present in maps of individual loci (Fig. 3A, B). Thus, we used the Pearson correlation between the templates and the maps from step (i) as a threshold validity metric. We calculated this metric for a range of thresholds and it appeared to have one major maximum for each data sample (Suppl. Fig. 3A, B, E). The selected thresholds were the ones corresponding to these maxima with two exceptions: (1) if the maximal correlation was lower than 0.2, the structure was considered to be absent in the studied map; (2) if the correlation exceeded 0.9 around the maximum, the threshold was selected to be the minimum value at which this correlation is achieved, The reason is that in such cases the correlation reached a broad plateau with uncertain, not robust position of the maximum; hence, this procedure retained more meaningful instances of the pattern.

**Figure 3.**
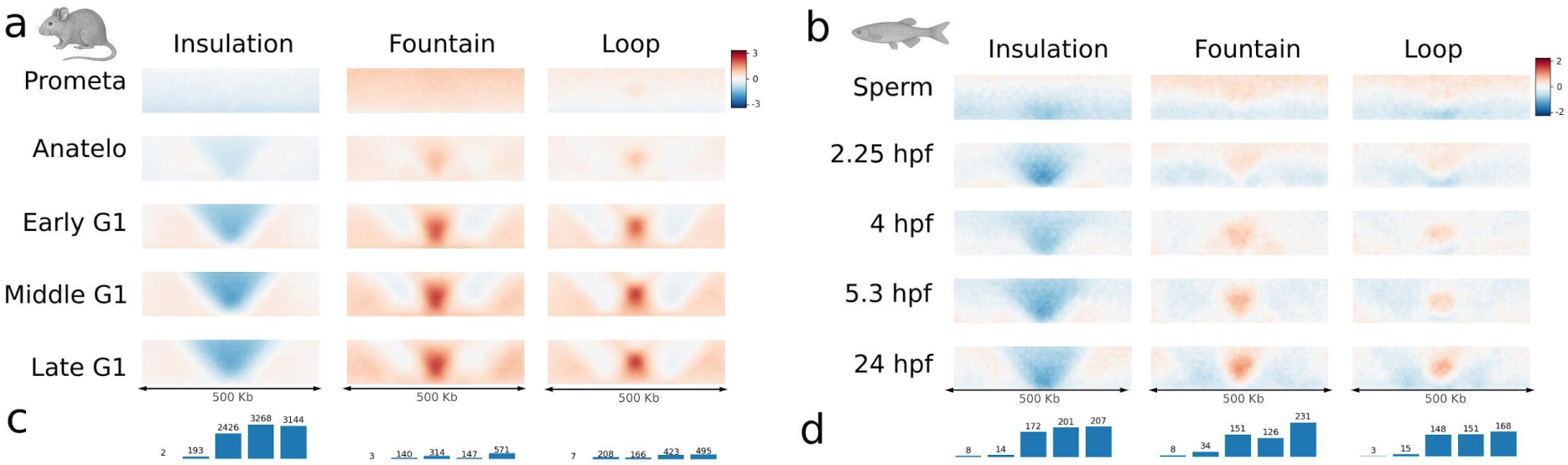
Chimaera quantification of the input patterns in input Hi-C maps across two processes in two species: (A, C) *M. musculus* cell cycle (G1E-ER4 cells, data from (65)), and (B, D) *D. rerio* embryogenesis (whole-embryo, data from (14, 113)). hpf — hours past fertilization. (**A, B**) Average pileup of the top 0.5% findings of insulation, fountain, and loop patterns (from left to right) using the latent space projections. (**C, D**) Numbers of 4 Kb bins with projection larger than the selected thresholds for each stage. See Methods “Pattern calling in Hi-C maps” for the description of the threshold selection procedure. The bars correspond to the stages in (A, B) and the numbers are shown above each bar.

### *De novo* DNA motif search with Integrated Gradients (IG)

*Integrated Gradients* (100) is a method that allows for obtaining information about positions in the input data that most significantly affect the model prediction. The IG is calculated based on the inputs and weights of the model, with higher values corresponding to higher influence.

We calculated the IG for the DNA encoder model as follows:

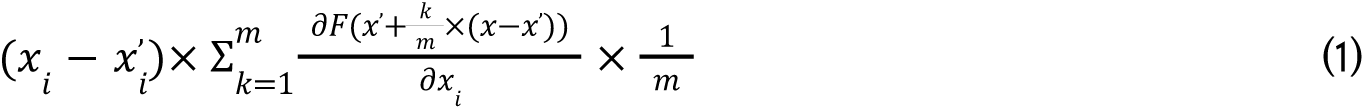

where: *i* is a position in the input data (DNA nucleotide position), *x_i_* is the initial value in the given position (DNA base), *x’* is the base level (no DNA input, empty input), *m* is the number of interpolation levels, *k* is a given interpolation level, and *F* is the neural network model (fully differentiable). We set the number of interpolation levels to 20.

The resulting values of integrated gradients were used to detect the importance of the genomic position on the prediction (Fig. 6A, 7M).

To select the most impactful regions for the prediction by IG, we selected the maximal value for each genomic fragment. We then extracted the 50 bp sequence around the peak and set profile values for it to zero. Then this procedure was repeated 15 times for each DNA fragment. Then the resulting 50 bp sequences were fed into the re-implemented MEME algorithm to find motifs.

### *De novo* DNA motif search with Gradient Descent (GD)

*Gradient Descent* is a method for searching DNA motifs associated with a specific pattern of Hi-C maps. This method is based on the latent representations of Hi-C maps obtained by the Hi-C encoder (see Suppl. Fig. 1D for schematics of the GD motif search).

(i) As an input for this method, we provide a map with a pattern template at its center (for example, insulation). This map is converted to its latent representation by the Hi-C encoder.

(ii.a) We then select a random DNA fragment from the genome (test sample), and replace the central 20 bp (the *target window*) with random nucleotides (sampled with frequencies from uniform distribution). This serves as a random initialization for the GD.

(ii.b) Next, we represent the input sequences as one-hot encoded, where each position is a 4-digit vector (one digit for each nucleotide). We then pass the matrix representation of the whole sequence through the softmax activation layer. This transformation does not affect non-target DNA positions since the softmax transformation of one-hot encoded vector reproduces the same vector. For target DNA positions, the softmax transformation works as a normalization of each column (nucleotide position) to 1, preventing the algorithm from converging to uniform sequences.

(iii) We then take the DNA encoder (note that the weights of the neural network are frozen at this stage), and provide it with the one-hot encoded DNA from (ii). This results in the latent representation of DNA sequence in the same space as the pattern template representation from (i).

(iv) Next, we calculate the loss function for GD, which is the negative projection of the latent vector predicted from the input sequence onto a latent vector predicted from a Hi-C pattern template. We backpropagate the resulting loss function through the network (without modifying the weights) but update the values in the target window of the one-hot encoded input DNA. The values of the remaining input sequence are not changed.

(v) We repeat steps (ii)-(iv) in total 50 times (after confirming qualitative convergence of the resulting target window). As a result, the one-hot encoded target window passed through the softmax layer can be used as a positional frequency matrix.

### *De novo* DNA motif search with Evolutionary Search (ES)

*Evolutionary Search* is another targeted motif search method, implemented as a genetic algorithm based on *in silico* mutagenesis (see Suppl. Fig. 1E for the schematic of ES for motifs).

(i) A map with an ideal pattern template centered within it is provided as input. This map is then converted into its latent representation using the Hi-C encoder, equivalent to step (i) in GD (see above).

(ii.a) We generate 200 random short sequences (10-20 bp) with a uniform nucleotide distribution. The sequence length is chosen based on the organism, and both shorter and longer sequences were tested for each species.

(ii.b) For each short sequence, a DNA fragment is selected from the test sample, and multiple copies of the short sequence are inserted at the center of this fragment (10 copies with a 20-nucleotide interval). Each such insert is thus embedded within a random DNA context from the real genome.

(iii) Similar to step (iii) of GD, we then take the DNA encoder, and provide it with one-hot encoded DNA sequences from step (ii). This results in the latent representation for each input DNA sequence in the same space as a representation of the pattern template from step (i).

(iv) *Estimate fitness.* We calculate projections of DNA-based latent vectors onto the ideal Hi-C-based latent vector and raise this metric to the 10th power to accentuate differences. The resulting values serve as fitness scores for each short sequence. Standard normalization is applied to the fitness score distribution to maintain consistent selection pressure.

(v) *Reproduction.* Next-generation sequences are derived based on fitness scores. We normalize the fitness scores from step (iv) by their sum to use them as sampling probabilities. Then, 180 sequences are sampled with replacement, allowing sequences with higher fitness scores to have more offspring and lower-fitness sequences to have fewer or none.

(vi) *Mutation.* Each offspring sequence undergoes random mutations, with a 5% probability of substitution at each position.

(viii) The top 20 sequences from the previous generation, ranked by fitness, are carried over to the next generation. This ensures that highly fit sequences are retained in the population, preventing loss of the best candidates.

(ix) The new sample of 200 short sequences then serves as input for the next iteration, cycling through steps (iii) to (vii). This process repeats for a set number of epochs or until the mean fitness reaches a plateau.

(x) Finally, a position frequency matrix (PFM) is constructed from the resulting sample, summarizing the evolutionary search outcome.

### Tools for DNA motifs and *in silico* mutagenesis

In this work, we implemented and used a wide variety of the tools for working and DNA sequence and DNA motifs:

i. obtain known DNA motifs from the JASPAR database (101) as position frequency matrices (PFMs) (Fig. 6, bottom),
ii. convert PFMs to positional weight matrices (PWMs) by applying the log2 transformation to the background-normalized frequencies with a pseudocount, assuming equal nucleotide contribution at each position,
iii. retrieve consensus sequence from PFM and sample sequences based on a given PFM – for *in silico* mutagenesis,
iv. insert sequence into genomic DNA sequence in different orientations and combinations – for *in silico* mutagenesis,
v. search for *de novo* motifs in a set of DNA sequences, based on MEME (102) – for Integrated Gradients (Fig. 6),
vi. compare motifs with those from JASPAR database (101), based on Tomtom (103),
vii. manipulate PFMs: reverse complement, randomize frequencies, shuffle positions. These tools were applied consistently throughout the manuscript.

### Checking motifs by in silico mutagenesis

#### Significance of motif impact

For each known or *de novo* motif, we tested its significance by *in silico* inserting the sequences sampled from the motif into the genome and predicting the effect of insertion. Each insertion into the test region was in 10 copies with the 20 bp step. For a given motif, we predicted maps for wild type and mutated sequences and calculated the difference. We then calculated the maximal absolute value of the difference, and used it as a metric of the impact. This procedure was repeated 50 times with randomly picked genomic regions. As a control, we used two types of randomizations: shuffled motifs (used in Fig. 5E, *ps* values in Fig. 6) and random sequences of the same length (used in Fig. 5F, *pr* values in Fig. 6). For each randomisation, the procedure was the same and yielded 50 values of the control metric. To calculate significance, the two-sided Mann-Whitney test on the observed and control samples was used.

#### PWM-based deletion of CTCF sites from the genome

To remove CTCF sites from the genome for predicting maps of cells with depleted CTCF (Fig. 5D), we used the vertebrate CTCF PFM from the JASPAR database (101). To account for cryptic sites, we selected a threshold yielding the number of sites slightly larger than the reported number of CTCF protein molecules bound to the mouse genome (104).

### Analysis of gene positions

In an approach similar to motif-based predictions via *in silico* mutagenesis, we developed a method for assessing gene-based effects. Using genome annotations from Ensembl (105) and RefSeq (106), we defined the following regions based on transcript annotations:

1. Promoters: regions around transcription start sites (TSS), spanning from -200 bp upstream to +100 bp downstream.
2. Gene bodies: regions from TSS to transcription end sites (TES), including both exonic and intronic sequences.
3. Intergenic regions: sequences between gene end and start annotations.

Promoter regions were used for collecting statistics for Integrated Gradients in Fig. 7N-R.

#### Gene-based in silico mutagenesis

To design and test the sequences for assessing the effect of gene positioning in the DNA (Fig. 7A-L, Suppl. Fig. 11), we implemented and run the following steps:

i. We created a background DNA sequence by concatenating random intergenic regions until reaching the desired fragment length, producing a gene-free sequence.
ii. This background sequence was used to predict 3D genome organization with Chimaera, serving as a baseline for later gene-insertion comparisons.
iii. To insert genes, we set the target gene length, usually the one that allows for eight insertions within the fragment (see the setup in Fig. 7A-L) and closely matches the species’ median gene length. We randomly selected eight genes within 20% of the specified length.
iv. These selected gene sequences were then inserted into the background sequence, replacing non-gene DNA.
v. Using the modified sequence, we predicted its 3D genome organization with Chimaera, generating the target structure to be compared with the baseline from (ii).
vi. We subtracted the baseline map from the target map to quantify the effect of gene insertion on the 3D organization (*effect maps*).
vii. Steps (i)-(iv) were repeated 256 times, and the resulting effect maps were averaged.

This gene-based *in silico* mutagenesis approach enabled us to estimate the influence of various types of gene insertions on the genome architecture.

### Cross-predictions

#### Selection of genomic regions for cross-predictions

For each species in the analysis, we selected 5-20 Mbp-long continuous genomic regions from the test sample, prioritizing the regions with the minimum number of unmapped bins. Note that Chimaera predicts Hi-C maps for fragments of size ranging from 65 to 500 Kb (Suppl. Table 1). Thus, we segmented these selected genomic regions into fragments of size specific to each model.

#### Cross-predictions run

We then input the selected DNA regions from each species to models trained on other species. Since the region of each species was continuous, and we ran the prediction with the step equal to half the fragment size, we were able to reconstruct the Hi-C signal for the whole DNA region (for the genomic distances below the model receptive distance, see Methods).

#### Model output adjustments to enable cross-species comparisons

Because the fragment lengths accepted by the models varied across organisms, the predicted maps were trimmed and scaled to a uniform size.

#### Cross-species comparisons

We then calculated the Spearman correlation between map windows predicted by each model against maps predicted by the model trained on the correct organism. To estimate significance, we also shuffled the windows to get correlations for random pairs. Then we applied the Mann-Whitney test to correct and control correlations for each pair of organisms. The resulting *p*-values were corrected by the Bonferroni procedure.

#### Building the tree

The resulting correlation matrix was symmetrized by averaging with the transposed one. Then it was subtracted from 1 and used as the distance matrix for the Neighbor joining algorithm to build a tree. The tree was visualized using the iTOL service (107).

## RESULTS

We start by designing a machine learning model (Fig. 1) that has DNA sequence as an input, predicts a Hi-C map for a given region, and encodes the information about the genome folding in a lower-dimensional vector, akin to word2vec for text (108) or DNABert for DNA sequences (109). We adopt a two-step training process that first learns Hi-C patterns and then links them to DNA sequence.

### Chimaera learns representations of Hi-C maps

The first part of the Chimaera model – the Hi-C encoder – is trained to encode the Hi-C map via low-dimensional latent representations (Fig. 1A). We first normalize the input matrices of Hi-C interactions by iterative correction to avoid sequencing and mappability biases (89). The contact frequency is known to vary over several orders of magnitude and drop rapidly with larger genomic distances (110, 111), which might complicate the training and ability to automatically detect delicate patterns in Hi-C maps. We thus normalize the contact frequencies by expected for each genomic separation (90). These observed-over-expected Hi-C maps are then 45-degree rotated, truncated to remove noisy long-range interactions, cut into fragments and subjected to a convolutional autoencoder. When the Pearson correlation between true and reconstructed maps stops growing, we stop the training, check the quality of reconstruction and denoising of Hi-C matrices (Fig. 2A-D), and freeze the weights of the Hi-C decoder.

The following properties of the latent space are important: (i) the zero vector corresponds to a map with uniform interactions and no other specific structures (Suppl. Fig. 2B-C), and (ii) the latent space is continuous, so that a small shift in any direction and from any point yields small changes in the decoded map (Fig. 2E). As an illustration of these properties, we implemented *in silico* generation of a given pattern at the center of Hi-C map by a stepwise movement in the latent space. For example, by moving along the vector encoding the insulation pattern, we can generate insulation in the central bin of a real Hi-C map (Fig. 1D, E, Fig. 2F).

Notably, the Hi-C encoder can embed not only real Hi-C datasets but also succeeds at embedding artificial user-supplemented maps of interactions, such as stereotypical insulation, dot, or fountain (Fig. 1D, E). Of note, the insulation and fountain patterns are distant from each other in the embedded space, suggesting that they are different and rarely overlap in real Hi-C maps.

### Chimaera enables quantitative calling of loops, fountains, and insulation in Hi-C maps

We next asked whether the latent space representation would allow us to detect typical patterns of the genome organization, such as insulation, loops, and fountains (Suppl. Fig. 1A and Suppl. Fig. 4). Motivated by our observation of the continuity of the latent space and significant spread of representations of typical structures there, we defined a score of similarity to a given, already known, pattern of the Hi-C map (e.g., the insulation triangle of depleted interactions). For that, we projected the target genomic bin (the one queried for the presence of the pattern) onto the vector corresponding to a map with the known pattern in the latent space (Suppl. Fig. 4). We then used these projections to study the development of three features (insulation, fountain, and loop) over the course of the mouse cell cycle (65) and fish embryogenesis (14) with 500 Kb windows (Fig. 3), with the choice of templates dictating the characteristic size of the patterns (e.g., the loops had the characteristic size of ¼ of the window size (see Methods for details).

At the early stages of both processes, we observed very small similarities to any of the studied patterns, suggesting little prominence of these structures in the Hi-C maps at these stages (Fig. 3A-B). However, the similarities grew with the progression into either the cell cycle or the embryo development. We calculated the number of the genomic regions scoring above a given threshold (see Methods “Pattern calling in Hi-C maps” and Suppl. Fig. 3A-B for the description of the threshold selection procedure) and treated them as hits, for example, insulating boundaries. This allowed us to quantitatively trace the changes in Hi-C maps in these biological processes (Fig. 3C-D).

In the cell cycle, all features are almost absent in prometaphase and gradually progress from ana-telophase to late G1 (Fig. 3A, C). Notably, loops emerge together with TADs and are positioned as their corner peaks, suggesting that they are formed by the same mechanism, likely loop extrusion (although some of the called loops might be small or micro-compartments (112) that have similar appearance in the Hi-C map). In fish embryogenesis, all structures are not expressed at the early stages and emerge at the 4 hours past fertilization (hpf) stage (Fig. 3B, D). In line with the emergence of fountains at zebrafish zygotic genome activation (ZGA) (14), we observe fountains appearing at the 4 hpf (weak) and 5.3 hpf (stronger) stages (Fig. 3B, D).

However, large fountains (2-5 Mb, flares (113)) have also been reported for zebrafish sperm stage but could not be probed with the selected window size (500 Kb). Thus, we repeated feature calling with 3.2 Mb windows (Suppl. Fig. 3C-E). At this resolution, fountains are profound at the sperm stage and absent at the 2.5 hpf stage. Notably, at the later stages (4 and 5.3 hpf), top hits for the fountain template of this artificially large size (the typical reported size of fountains in D. rerio ZGA is 50-200 Kb size (14)) are mostly compartment-like structures (Suppl. Fig. 3C).

Together, these results indicate that the latent space of Chimaera preserves the information about typical patterns of the 3D genome organization (insulation, fountains, and loops), and is suitable for differentiating these patterns, akin to semantic embedding in the natural language processing (114).

### Chimaera predicts 3D genome architecture from DNA sequence across diverse species

The second part of the Chimaera model — the DNA encoder — is trained after the Hi-C encoder is fixed. The DNA encoder takes a DNA sequence as the input and learns to predict the representation of the Hi-C map for the respective genomic region in the latent space (Fig. 1B-C). By combining the pre-trained Hi-C decoder with the DNA encoder, we achieve the linkage between the genomic sequence and characteristic patterns of the genome organization.

To test this capability of Chimaera, we predicted the effects of a genomic rearrangement associated with congenital F-hand syndrome (Fig. 4B). While the Hi-C map for this genetic variant is not available, Chimaera predicts that, following the rearrangement, human cells form a domain around the Wnt6 gene enhancer that also includes the Wnt6 gene itself. This configuration leads to deregulation of Wnt6, which plays a crucial role in the embryonic morphogenesis.

**Figure 4.**
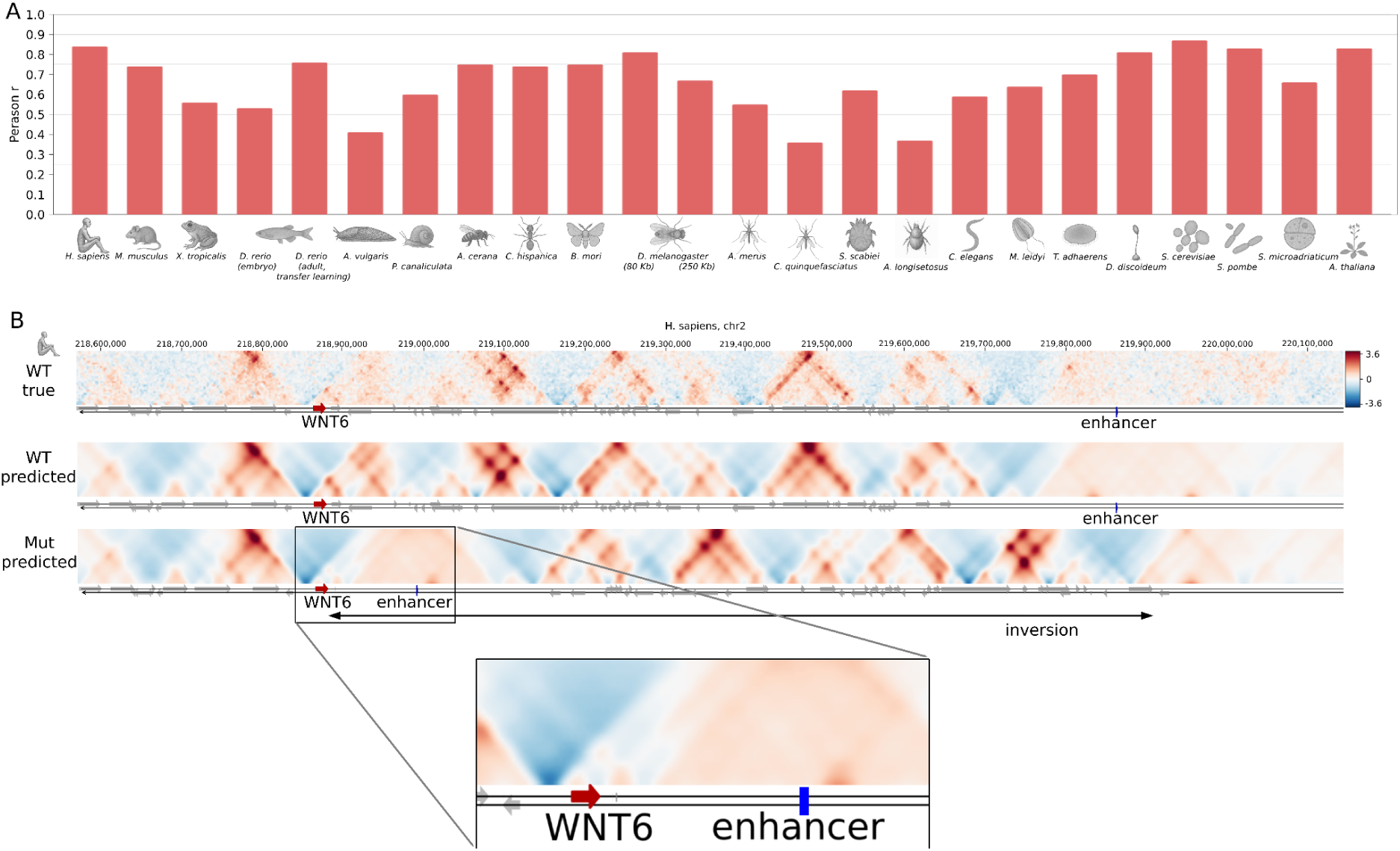
Chimaera predictions. **(A)** Median Pearson correlations between true and predicted maps at the best distances for test samples of all studied organisms (see Suppl. Table 1 and Suppl. Fig. 6 for more details) **(B)** Prediction of the effect of structural variant that causes the F-hand syndrome during development. Wild-type Micro-C data for the HFF6c cells is from (63), the rearrangement coordinates are based on (16, 115). The rearrangement that *in vivo* causes the F-hand syndrome is an inversion that relocates the enhancer closer in the linear genomic distance to the WNT6 gene promoter. However, the chromatin organization of this rearrangement is not known (while prediction exists for other variants causing F-hand and limb malformations (49)). With Chimaera, we predict the effect of this rearrangement and observe the formation of the domain around the enhancer. This domain also includes WNT6, providing the basis for de-regulation of WNT6.

Chimaera is a model with a relatively small receptive field (500 Kb, in comparison to recent models like Borzoi (52)), and has a similar number of parameters and computational resource requirements to existing alternatives (∼3-5M parameters as opposed to ∼700K parameters for Akita (48) and ∼4M parameters for ORCA (49)). Thus, we did not aim to achieve significantly better performance of prediction against Akita (48) or Orca (49), but instead aimed to improve the interpretability of the model. Yet, when we tested Chimaera against an adapted version of Akita, we observed a better or similar prediction quality (Suppl. Table 2).

Next, we studied the limits of Chimaera’s ability to learn complex DNA and chromatin folding patterns. We trained Chimaera on data from 22 different organisms, including animals, plants, fungi, amoebozoans and alveolates (Fig. 4A, Suppl. Fig. 5, Suppl. Table 1). For fruitfly *D. melanogaster*, we trained two models on different receptive fields, 80 Kb and 250 Kb.

For species with high-quality Micro-C data, known for its higher resolution and feature sharpness (63), we observed substantially better prediction performance (median Pearson correlation: 0.79 in human, 0.68 in mouse, 0.76 in fruitfly, 0.86 in fission yeast *S. pombe,* 0.79 in plant *A. thaliana*). However, some datasets showed lower prediction accuracy. We hypothesized that this is due to lower data quality (Suppl. Table 1) and applied a transfer learning approach. Although we tried this approach on multiple species (data not shown), it showed the best improvement for *D. rerio*, where training on original data resulted in correlations of 0.4, and pre-training on human Micro-C data followed by fine-tuning on *D. rerio* improved correlations to 0.6 (Suppl. Fig. 8A, B). Interestingly, without fine-tuning, the human model achieved 0.4 correlation on *D. rerio*, indicating conserved chromatin folding patterns among vertebrates.

For all species, we achieved significant but variable quality of prediction (Fig. 4A, Suppl. Table 1, Suppl. Fig. 5, also Suppl. Fig. 6-7). We hypothesised that this could be due to variable quality of the input datasets. To test that, we extracted several quality characteristics of Hi-C/Micro-C datasets, including resolution, number of non-zero bins, cis-to-trans ratio, number of contacts below the receptive distance of the model (Suppl. Fig. 7). Although each individual characteristic did not significantly correlate with the Chimaera performance, linear combination of pairs of characteristics explained the performance well for most of the species (R2 of 0.547 for resolution and cis-to-total ratio, 0.545 for non-zero bins and size of genome in the training set). For *Arion vulgaris, Culex quinquefasciatus*, and *Archegozetes longisetosus* (Suppl. Table 1), the Chimaera performance was the lowest (the maximum Pearson correlation at the best diagonal was lower than 0.5), which could be attributed to poor Hi-C data quality (Suppl. Fig. 7). We thus removed these species from further analysis.

Finally, we measured the Chimaera performance with alternative, structure-based metrics that are specific to the Hi-C/Micro-C data (see Methods, Suppl. Table 1). These metrics compare the recovery of key chromatin structures (loops, TADs, insulation, fountains) in Chimaera predictions. For example, the highest accuracy in predicting loops is achieved in *D. discoideum*, whose chromatin consists predominantly of loops, whereas insulation prediction is inferior to models for species with abundant insulation (Suppl. Table 1). In contrast, some models do not perform well on the common structures for their species, such as the *P. canaliculata* model at TAD reconstruction (Suppl. Fig. 5F). This effect may be explained by very low Hi-C data quality (Suppl. Table 1 and Suppl. Fig. 7), although we cannot exclude that TADs in these species arise as a consequence of a mechanism without local DNA determinants.

### Chimaera confirms the importance of known motifs for chromatin structure

Next, we aimed to confirm that Chimaera learns meaningful patterns from DNA sequences, performing *in silico* mutagenesis by insertion of known motifs of architectural proteins in different species.

As expected, insertions of several CTCF sites significantly affects predictions of models in Tetrapods (mouse, human, and frog), causing the values in the maps to change by up to 2 standard deviations (Fig. 5A, E). Insertion of this motif causes insulation at the position of insertion, while two loci of convergent sites form a loop between them (Fig. 5A). The YY1 motif produces a similar but smaller effect (Fig. 5B, E). Our model correctly accounts for the CTCF site orientation and can reproduce the results of *in vivo* CTCF site inversion. The effect of the CTCF site removal and inversion near the Sox2 gene, shown experimentally by (116), is well reproduced by Chimaera (Suppl. Fig. 9).

**Figure 5.**
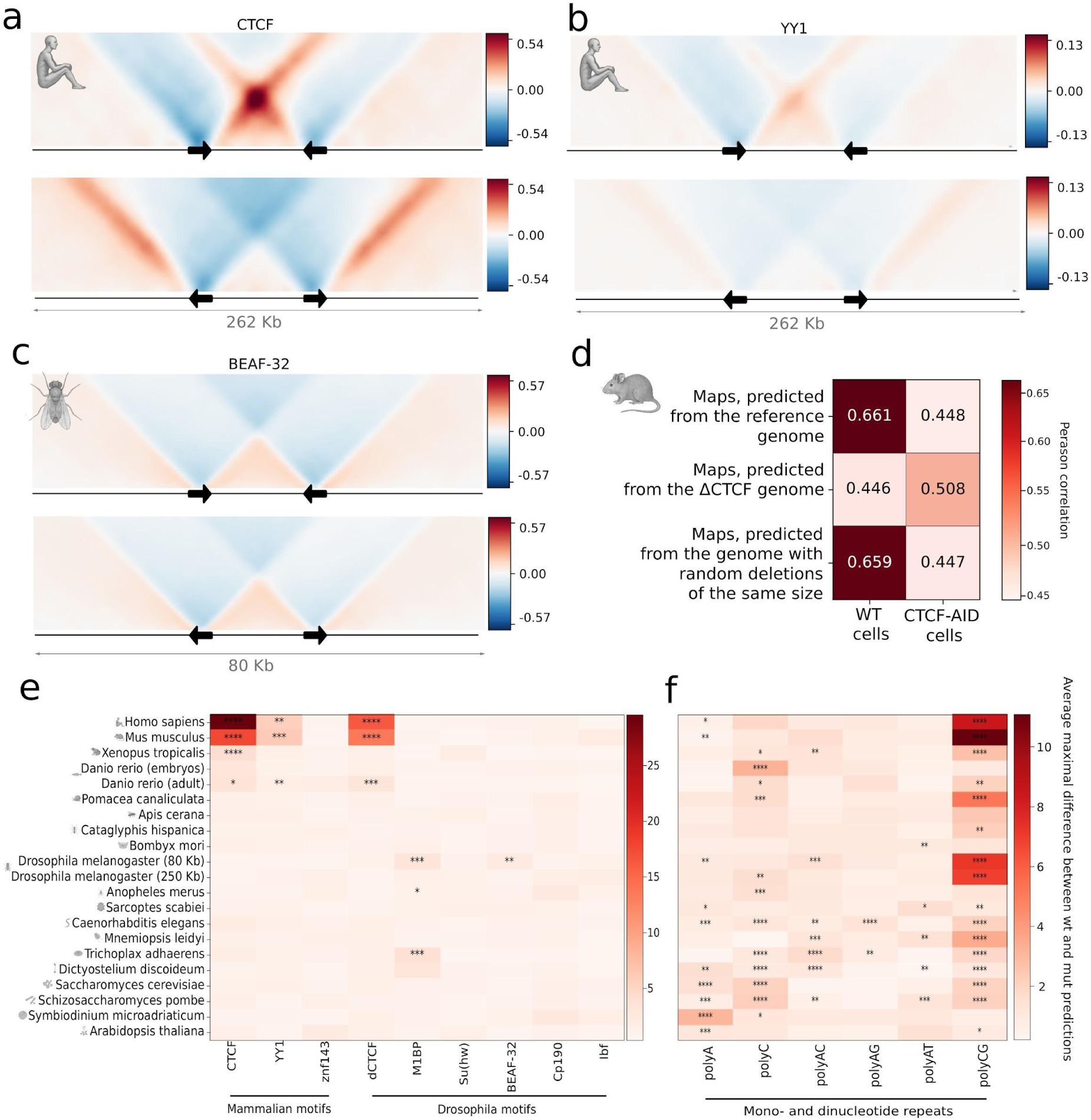
*In silico* mutagenesis shows importance of known DNA motifs for Chimaera predictions in diverse species. ΔCTCF – genome with *in silico* deletion of CTCFs. AID – auxin-induced degradation. wt – wild-type genome, mut – genome with *in silico* mutagenesis by insertion. * – *D. rerio* fine-tuned model on adult muscle cells (67) with transfer learning from human, ** – *D. rerio* model on embryo cells (14) without transfer learning. (**A-C**). The average effect of insertion of two tandems with sites into a large number of sequences. Top: convergent orientations, bottom: divergent orientations. Averaging over n=50 randomly selected genomic regions (from the Chimaera validation/test set). (**A**) Insertions of CTCF sites into the human genome. (**B**) Insertions of YY1 sites into the human genome. (**C**) Insertions of BEAF-32 sites into the *D. melanogaster* genome. (**D**) Average correlations of predictions with the true Micro-C maps of WT and CTCF-AID cells. Top row: predictions based on the reference genome, middle row: reference genome with *in silico* removed CTCF sites (via replacement by random sequences, see Methods, ΔCTCF), bottom row: reference genome with deletions of randomly selected regions (same length as the CTCF motif). The ΔCTCF-genome predictions are more similar to the CTCF-AID Micro-C map. Results for chromosome 19, which was not used for Chimaera training. (**E, F**) Average maximum changes in predictions caused by multiple insertions of sites with motifs of architectural factors (E) and simple repeats (F) into the reference genomes of different species (rows). Averaging over n=40 randomly selected genomic regions (from the Chimaera validation/test sample). Each row is normalised by its median value, p-values are adjusted using the Benjamini-Hochberg procedure. * *p*<0.05, ** *p*<0.01, *** *p*<0.001, **** *p*<0.0001.

Next, we retrieved six motifs of insulator factors from fruitfly *D. melanogaster* (CTCF, M1BP, Su(hw), BEAF-32, Cp190, Ibf (34)) and tested their importance for Chimaera predictions in different species (by comparing insertions of motifs sites to the shuffled motif controls, see Methods). Two motifs (BEAF-32 and M1BP) produced significant changes in *D. melanogaster* (Fig. 5E, insulatory effect for BEAF-32 demonstrated in Fig. 5C). Notably, this change was observed at 80 Kb, which was possible to study in *D. melanogaster* due to high quality of its Micro-C data (73), but not at 250 Kb, which was a typical resolution in other insects. Thus, the fact that we do not observe importance of these motifs for other insects (such as bee *A. cerana* and moth *B. mori*) can be attributed to lower quality of their Hi-C data, and might be improved with better quality of datasets.

Among other species, six insulators of *D. melanogaster* did not show any significance, with a notable exception of M1BP in mosquito *A. merus* and placozoan *T. adhaerens* (Fig. 5E). The strong effect of M1BP in *T. adhaerens* can be explained by the fact that the M1BP motif contains a 4-nucleotide pattern generally associated with insulation in this organism, as we show below. The *D. melanogaster* insulator dCTCF did not show significance in any of the insects, but had a significant impact in tetrapods, due to the similarity of the dCTCF and CTCF (Fig. 5E).

Next, we reasoned that repeated low-complexity sequences might be interlinked with genome organization and tested how *in silico* insertion of repeated nucleotides and dinucleotides would affect the predicted maps. The low-complexity sequence with the highest impact across all species was the CG repeat (Fig. 5F). In accordance with this observation, GC-repeats are enriched at insulation sites in jelly comb *M. leidyi* (23). While the CG importance might be expected for species with a CpG-methylation mechanism (e.g., vertebrates (117)), the importance of CG repeats in some organisms may result from another chromatin folding mechanism. For example, CGs are frequently present in yeast gene bodies (118), which might be captured by the model.

Finally, we tested Chimaera’s effectiveness for *in silico* mutagenesis by motifs removal. We took a mouse-based Chimaera model and replaced all instances of CTCF motifs identified in the genome by a position weight matrix (see Methods) with random sequences (ΔCTCF). Next, we checked whether this *in silico* perturbation resulted in serious changes in the prediction.

Predicted maps for the ΔCTCF genome showed lower similarity to untreated mouse Micro-C data (64) (correlation of 0.446) compared to maps predicted for the intact reference genome (correlation of 0.661, Fig. 5D). Conversely, maps predicted from the ΔCTCF genome were more similar (correlation of 0.508) to the Micro-C data for cells with CTCF depleted using the auxin-inducible degron system (CTCF-AID) than compared to the maps predicted for the reference genome (correlation of 0.448). To test whether other motifs can explain the genome organization of CTCF-AID data, we trained a separate model on the CTCF-AID Micro-C data (obtaining relatively high Pearson’s correlation of 0.55) and applied all our *de novo* motif search methods. Neither CTCF nor any other motifs were important for Chimaera prediction of CTCF-AID Micro-C maps. Together, these observations demonstrate that Chimaera effectively captures the critical role of CTCF and predicts the impact of removal of its sites from the genome.

### *De novo* search for motifs associated with chromatin structure

Search for DNA determinants of the genome architecture is challenging because it requires either *a priori* knowledge of potential DNA motifs or an exhaustive search by *in silico* mutagenesis. The search space is very large: for example, exhaustive search for a significant 10-bp motif would require testing insertions of ∼4^10^ sequences. While optimizations of computational time for exhaustive *in silico* mutagenesis exist (119, 120), they do not help with shrinking the search space. Thus, we designed a solution that (i) would be more targeted than the existing solutions but still (ii) would not require prior knowledge about the DNA motifs.

To that end, we first adapted the Integrated Gradient (IG) approach, which allowed us to calculate the importance of each input DNA position by estimating the change in the gradient in response to the gradual change of the given input (see Methods for a detailed explanation). This approach results in a genomic track of the importance of each genomic position (IG track) without requiring multiple model launches. Next, we detect peaks in the IG track and search for motifs with re-implementation of MEME (121) in the DNA sequences around them. The IG approach allowed us to detect motifs important for genome organization (Fig. 6A, B-left). Its findings were limited to the CTCF motif causing the insulation pattern in the species where the CTCF role was already demonstrated (mouse, human, frog, fish), and to the ARGCCAW motif similar to Site II motif in *A. thaliana* (Fig. 6B-left).

**Figure 6.**
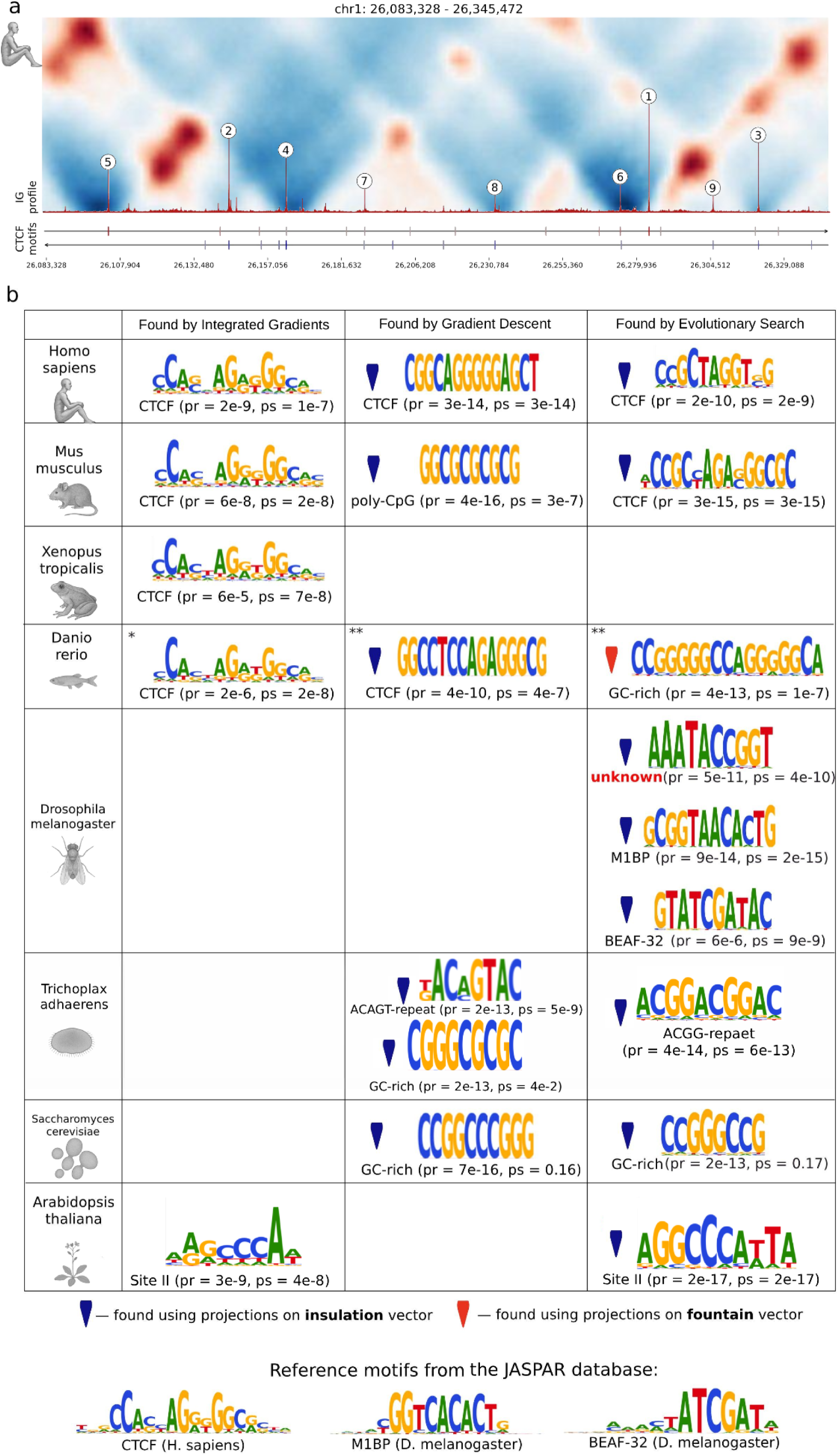
***De novo* motif search, three methods: Integrated Gradients (IG), Gradient Descent (GD), and Evolutionary Search (ES).** Motif impact significance: *pr* – *p*-value for the control with randomized motifs of the same length, *ps* – *p*-value for the control with shuffled motifs (see Methods, “Significance of motifs impact by in silico mutagenesis”). * – *D. rerio* fine-tuned model on adult muscle cells (67) with transfer learning from human, ** – *D. rerio* model on embryo cells (14) without transfer learning. (**A**) Example of an IG profile for a region of the reference genome (denoised map shown above). Peaks indeed correspond to CTCF sites detected in the same region (bottom tracks). Brightness of the color at the bottom panel refers to the CTCF site score based on its motif. (**B**) Table of motifs detected by three methods. For GD and ES we searched specifically for motifs associated with insulation (blue signs) or fountains (red signs). Empty cells indicate no motif found by a given method. The height of each letter in the motif logo represents the information content of the position in the position-specific weight matrix based on the method output. The ES and IG position weights can be interpreted as the importance for the prediction, but should be taken with caution as ES usually overestimates the importance of each position relative to the true motif (see the CTCF example in the table).

The IG approach does not distinguish between different patterns formed in Hi-C maps. Thus, the importance of a genomic position in DNA is a cumulative measure of its contribution to insulation, TADs, loops, fountains, and other Hi-C patterns around it. To overcome this limitation, we aimed to design an approach that (in addition to the requirements listed above) will be targeted at different patterns of 3D genome organization.

To that end, we utilized the model’s latent space and the encoded semantics of different chromatin patterns. As in the Hi-C pattern quantification approach above, we input a desired pattern into the Hi-C encoder (e.g., the insulation triangle of depleted interactions), and obtain its representation in the latent space. This representation serves as the target direction in which we can gradually push the input sequence for DNA encoder by modifying the nucleotides.

We implemented two ways of converging towards the target direction. In the approach that we called Gradient Descent (GD) in the DNA input space (Suppl. Fig. 1D), we define the loss between the desired target Hi-C embedding and the current DNA embedding, then backpropagate the gradients to the input DNA sequence, and iteratively repeat this procedure (see Methods).

Finally, inspired by the model of random mutagenesis and selection happening in natural biological systems, we designed Evolutionary Search (ES) similar to genetic algorithms in optimization (Suppl. Fig. 1E). At each generation of ES, random DNA sequences are generated and then selected for goodness-of-fit of the predicted maps to the desired latent representation. At the onset of each generation, DNA sequences are amplified proportionally to their fitness and then slightly randomly modified (see Methods).

Both GD and ES allowed us to explore all possible DNA variants, while focusing on target chromatin patterns. This effectively helps to reduce the number of model launches (relative to the exhaustive search of motifs) by shrinking the search space for the motif search, which is now focused around DNA motifs that can potentially generate specific Hi-C patterns.

We applied all three methods to the motif search across different species, with GD and ES targeting insulation and fountain patterns. Methods often produced different results and demonstrated varying efficiency for different organisms (Fig. 6B, see the Discussion for details), suggesting that the methods may complement each other and can be combined to produce the best results.

For all vertebrates, at least one of the methods identified motifs similar to the CTCF motif (see Methods). The CTCF motif, when found by GD or ES, was associated with insulation and never associated with fountains. For *D. melanogaster*, two ES motifs correspond to known insulators (BEAF-32, M1BP). Motifs rich in alternating C and G have been found for many organisms (associated with either insulation or fountains). For *Trichoplax,* we found repeating motifs ACGG and ACAGT (Fig. 6B). The former one belongs to a transposable element and was already shown to be associated with insulation (23). For *A. thaliana*, we found an AGGCCCATTA motif that is similar to the ARGCCAW motif detected by IG too, and might be related to the Site II motif (122). Also, a motif that could not be matched with any previously known motif has been identified for *D. melanogaster* (Fig. 6B).

As a control, we tested all the found motifs by *in silico* mutagenesis and confirmed that they indeed significantly affect the predictions (see Methods). By randomly inserting the resulting motifs into the genome and assessing the maximum change between true and predicted Hi-C maps, we estimate the significance of the motif impact by comparing to the expected control (Fig. 6B). We designed two expected controls, inserting random sequences of the same length (*pr* in Fig. 6B) and inserting shuffled motifs (*ps* in Fig. 6B, preserving nucleotide content). The first control shows that almost all found motifs are significant. The second control shows that the order of the nucleotides in GC-rich motifs usually does not matter (except for mouse and *D. rerio* embryos), confirming that these motifs represent the oligonucleotide sequence biases but not factor binding sites.

Finally, we questioned the biological relevance of the newly discovered motifs AAATACCGGT for *D. melanogaster* and AGGCCCATTA for *A. thaliana*. Although the latter motif might be Site II (122), it has not been previously associated with chromatin organization of this plant, and we decided to characterize its properties in depth. Strikingly, we found that both novel motifs are associated with active histone marks, producing a prominent peak at the average pileups, significantly larger than expected for randomized controls (Suppl. Fig. 10).

The *A. thaliana* AGGCCCATTA motif causes insulation with stripes emanating from its site (Suppl. Fig. 10A) which cannot be explained simply by its nucleotide content (shuffling control, Suppl. Fig. 10A). Its sites are typically located in open chromatin and are associated with a very high and narrow peak in the ATAC-Seq profile (Suppl. Fig. 10B), with H3K27ac and H3K4me3 histone marks enriched at ∼500 bp around it, suggesting strong positioning of nucleosomes (123, 124). The genes closest to the AGGCCCATTA motif are associated with RNA processing, suggesting that an AGGCCCATTA-binding factor in *A. thaliana* might be involved in regulation of this process.

The *D. melanogaster* AAATACCGGT motif causes mild insulation in Micro-C maps (Suppl. Fig. 10D), does not have a strong ATAC-Seq peak, but is enriched in PolII, BEAF-32, Cp190, and H3K9ac modification (Suppl. Fig. 10E). The absence of an open chromatin peak suggests that the binding at this site might happen without nucleosome displacement, akin to Zelda pioneering factor of *D. melanogaster* (125), whose binding indeed is enriched around the AAATACCGGT motif in our analysis (Suppl. Fig. 10E). Although our newly discovered motif is not similar to the reported Zelda motifs (CAGGTAG (126)), it might bind a Zelda co-factor or position nucleosome for Zelda binding.

### Chimaera demonstrates the importance of gene positions for chromatin structure in most studied organisms

Given the indispensable role of chromatin 3D organization in the regulation of gene expression, we next questioned whether the opposite effect holds true, that is, whether positioning of genes would influence the chromatin organization. Indeed, *in silico* gene insertions have a drastic effect on the Hi-C patterns predicted by Chimaera (Fig. 7). Additionally, gene starts in many organisms are important when examined by IG (Fig. 7M-R).

**Figure 7.**
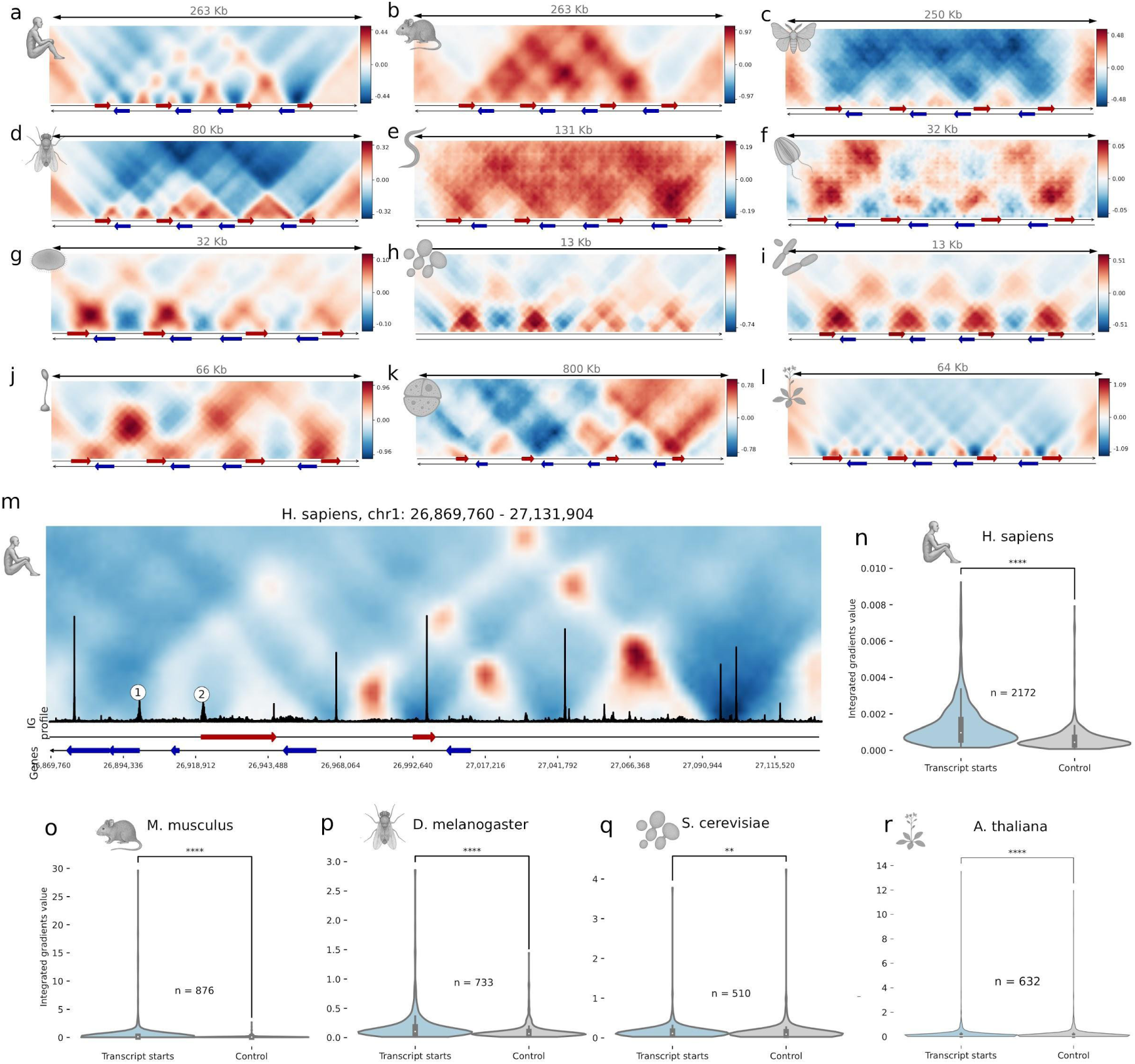
Effect of gene location on model predictions. Gene annotations are based on Ensemble database (105) for all species except dictyBase for *D. discoideum* (127). (**A-L**) Averaged predictions of Chimaera models based on *in silico* constructed sequences containing genes in the specified orientation. Genes are shown by arrows at the bottom of each panel. The color scale for each heatmap represents standard deviations of train samples, calculated for each organism independently. (A) - *H. sapiens*, (B) - *M. musculus*, (C) – moth *B. mori*, (D) – fruitfly *D. melanogaster* (80 Kb model), (E) – nematode *C. elegans*, (F) – jelly comb *M. leidyi*, (G) – placozoan *T. adhaerens*, (H) – budding yeast *S. cerevisiae*, (I) – fission yeast *S. pombe*, (J) – amoeba *D. discoideum*, (K) – dinoflagellate *S. microadriaticum*, (L) – plant *A. thaliana*. (**M**) Example of an IG profile for a region of the human reference genome (the denoised map shown above), based on a model trained on human. Broad IG peaks are present at promoters of some genes (genes are shown with arrows at the bottom). (**N-Q**) Comparison of average IG values around promoters against randomly selected genomic regions of the same size (control). The *p*-values are for the two-sided Mann-Whitney.

In most studied organisms, Chimaera reveals insulation at gene starts (Fig. 7A-D, G-I, L). In insects, it lacks any other regularities (Fig 7C-D). In mammals (Fig 7A-B), this rule is supplemented by loops between genes, in placozoan *T. adhaerens* and both yeast species (Fig. 7G-I), by more dense chromatin in genes and more open in intergenic regions. *A. thaliana* has also insulation at gene ends (Fig. 7L). Genes in *C. elegans* slightly interact with all their surroundings (Fig. 7E), while genes of jelly comb *M. leidyi* interact with each other (Fig. 7F).

Interestingly, two species with asymmetric patterns for convergent and divergent genes (amoeba *D. discoideum* and dinoflagellate *S. microadriaticum*) have qualitatively different structures in these contexts. In *D. discoideum*, Chimaera predicted loops between two convergently oriented gene pairs but not two divergently oriented gene pairs, suggesting that the orientation of genes plays a key role in formation of this Hi-C pattern (Fig. 7J). At that, highly expressed genes in the *D. discoideum* development (37) have a larger impact on the 3D genome (Suppl. Fig. 11A), which may be correlated with oligonucleotide frequencies in genes with various expression levels (Suppl. Fig. 11B). In *S. microadriaticum,* convergent gene pairs cause insulation, while divergent gene pairs have enriched interactions across them (Fig. 7K), consistent with observations in (81). This difference between *D. discoideum* and *S. microadriaticum* suggests that *S. microadriaticum* structures might be caused not by slow loop extruders pushed by the moving polymerase, as proposed for *D. discoideum* (*37*), but by some other mechanism.

### Cross-Species Chimaera modeling reveals evolutionary conservation and divergence in chromatin folding mechanisms

Applying Chimaera for chromatin feature quantification, *in silico* mutagenesis, and *de novo* motif discovery, we demonstrated the existence of species-specific chromatin folding rules. A well-trained Chimaera model tailored to a specific organism can act as a black box, effectively representing the nuclear environment and *in silico* folding of distinct 3D genome structures.

We thus decided to use multiple Chimaera models as a proxy to assess differences in the chromatin folding rules between species. To that end, we folded DNA sequences of one species by all possible models trained for other species, and assessed the quality of the predictions. When repeated for all species in our analysis, this allowed us to build the complete matrix of cross-predictions (Suppl. Fig. 12).

In most cases, species that are taxonomically close exhibited high cross-prediction accuracy; for example, models trained on human data accurately predicted Hi-C structures from mouse DNA. However, close species usually have similar GC-content, which in theory can dictate the model preferences. To test this, we utilized the setup of the unique experiment by Meneu et al. (79) that artificially introduces GC-neutral DNA of bacteria *Mycoplasma pneumoniae* (40% GC) and GC-poor DNA of *Mycoplasma mycoides* (24% GC) into the budding yeast nucleus (38% GC). By launching the Chimaera trained on yeast *S. cerevisiae* for these two genomes, we show that Chimaera successfully recapitulates DNA folding patterns of exogenous DNA with both similar and different DNA composition (median correlation 0.5 for *M. pneumoniae* and 0.46 for *M. mycoides*, Suppl. Fig. 13A and individual examples in Suppl. Fig. 13B). Thus, Chimaera effectively predicts folding of DNA with different compositions, albeit with slightly lower accuracy than for original DNA. This example demonstrates that Chimaera captures the rules of chromatin folding of the species it was trained on.

This observation prompted us to generate a cluster tree based on cross-prediction matrices (Fig. 8). This chromatin-based tree agrees with the phylogenetic tree for vertebrates, yeasts, *Diptera* and *Hymenoptera*, which form distinct clades. Thus, chromatin folding mechanisms are largely conserved over short evolutionary distances. However, our chromatin-based tree is not well resolved at large evolutionary distances, suggesting that cross-predictions between distant organisms give little information due to largely different mechanisms of chromatin organization. For example, dinoflagellate *S. microadriaticum* with distinctive crystalline chromosomes has no significant positive correlation with any other studied organism (Suppl. Fig. 12), with the basal location in the chromatin-based tree (Fig. 8). Thus, our observations suggest that chromatin folding mechanisms are conserved and can result in successful cross-species predictions within classes or phyla, but do not reflect evolutionary divergence at larger distances.

**Figure 8.**
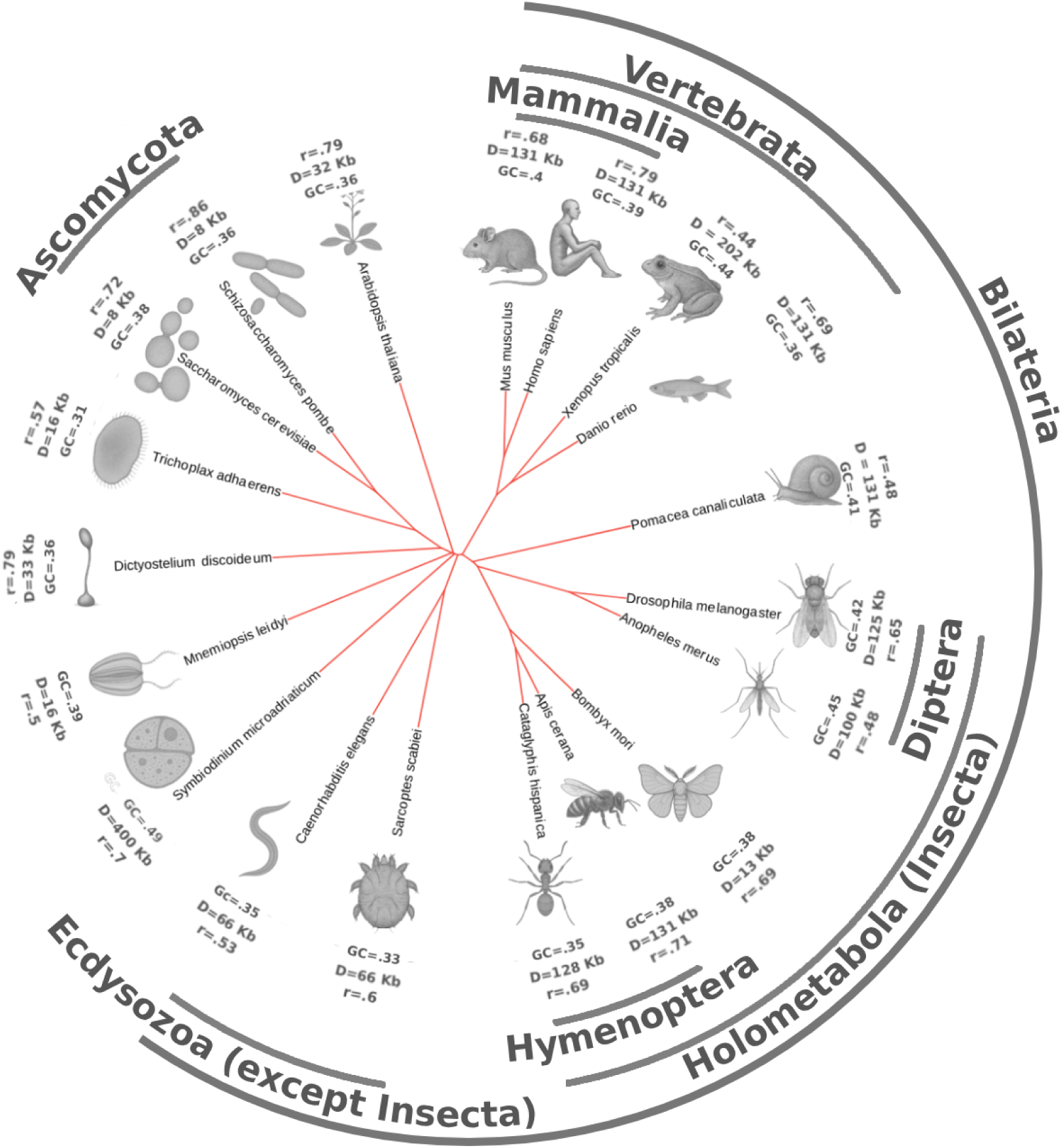
Cluster tree of the studied species based on the correlation matrix of cross-predictions of models trained for different species. The matrix of cross-species predictions is presented in Suppl. Fig. 12. The tree is built using the Neighbor-Joining algorithm. Arcs near the tree show taxonomic groups corresponding to the underlying branches. For *D. melanogaster*, we used the model with a 250 Kb receptive field for better comparison with other insect models, for *D. rerio* the model trained on adult fish was used. Next to each species, we display three factors that can potentially affect the tree: the GC-content (GC); the receptive distance (the maximal distance of contacts visible to the model, set to half of reception field size, or 64×resolution, see Methods); *r,* the model’s median Pearson correlation coefficient (full map) on the test sample.

## DISCUSSION

Recent studies on chromatin organization have predominantly focused on mammals, from which we have uncovered key mechanisms such as loop extrusion resulting in the formation of loops between CTCF sites (1, 12, 110). However, chromatin folding rules, even the rules of loop extrusion, vary widely across species, resulting in some organisms exhibiting distinct patterns, like fountains in the fish chromatin (14), and loops between convergent gene pairs in amoeba *D. discoideum* (*37*). Comparative analyses of chromatin folding between species are typically restricted to closely related genomes (45, 46, 128–130), conserved syntenic regions (11, 39, 41, 44, 131), and homologous genes (40, 42, 132), or rely on highly abstract qualitative comparisons of Hi-C maps (21, 22, 43, 133, 134). A significant advancement in the field came with the development of Akita (48), a deep learning-based neural network capable of predicting Hi-C data from DNA sequences. Akita was one of the first tools to successfully fold the mouse genome using a model trained on human data, revealing differences in the chromatin folding rules based on the observed discrepancies in predictions between species. Another breakthrough in sequence-based chromatin features prediction was the Enformer model (135), which takes advantage of the transformer architecture to predict RNA-Seq and epigenetics tracks. Despite the transformers’ powerful performance, they are known to require large training samples (136) so they are not immediately applicable organisms with small genomes that are also of considerable interest from from the functional and evolutionary point of view

Continuing the trend set by Akita, we have designed Chimaera, a neural network that allows for easy interpretation of the learned mechanisms and diverse cross-species predictions. Chimaera demonstrates good performance for Hi-C map reconstruction, especially for the organisms with high data resolution, and outperforms other existing models on organisms with small genomes (Suppl. Table 2). Notably, these results are achieved despite the fact that Chimaera: (i) uses only information about DNA and not epigenetics as input for the Hi-C map prediction, (ii) might be biased against rare, underrepresented structures of the genome, (iii) uses reference genome as input and omit genomic variants that can affect genome folding (such as cell line-specific mutations in CTCFs), does not account for global properties of chromatin organization such as *P*(*s*), the contact probability at a distance, and (v) has a limited receptive window both on DNA and Hi-C/Micro-C map enforcing Chimaera to learn only local patterns.

Chimaera’s main strength is not the quality of the prediction of the Hi-C map (although it is competitive in this respect as well), but the interpretability of the learned folding rules. We achieve interpretability by a characteristic chimeric design of our model based on an autoencoder. Firstly, the Hi-C encoder is trained to compress the Hi-C map snippet into a multi-dimensional vector. During training, the Hi-C encoder is exposed to various 3D genome architectures across the genome and learns various stereotypical patterns, such as insulation, TADs, loops, and fountains. As in semantic embeddings in natural language processing (137), the model learns the characteristic features of chromatin that can be then extracted from the embedded space and used, for example, for feature generation (Fig. 2E-F) and quantification (Fig. 3, Suppl. Fig. 3-4). Since feature calling relies on an auto-encoder typically trained on a single species, it can also be used for feature quantification in different cell types (Suppl. Fig. 4) and stages of biological processes (mitosis and embryogenesis, Fig. 3).

Next, we (i) split the Hi-C autoencoder and retain its decoder part, (ii) freeze the weights of the Hi-C decoder, and (iii) connect it to the DNA encoder that is trained to predict the compressed Hi-C maps through the same multi-dimensional latent representation. This strategy allows us to avoid fitting the noise of the data, and separate the task of learning the biological rules from the task of drawing the Hi-C map (Suppl. Table 2). As a result, Chimaera matches Akita in sequence-based prediction accuracy (Suppl. Table 2) and extends to diverse contexts, genomic rearrangements (Fig. 4B), targeted CTCF-site edits (Fig. 9), and exogenous inserts (Suppl. Fig. 13), highlighting its generalization capacity.

Moreover, the chimeric design allows us to use the semantics of the embedded space to connect stereotypical features of chromatin to underlying DNA determinants. As a proof of concept, we confirm the most prominent association “insulation - CTCF motif” in a wide range of species with both Chimaera’s *in silico* mutagenesis, and *de novo* motif search. While the original Chimaera training on *D. rerio* did not produce any association with CTCF, the adaptation of transfer learning from human data demonstrates the importance of CTCF motifs for the insulation pattern in zebrafish, in line with recent studies (138) (Fig. 5E). Additionally, we confirm the correctness of the model by *in silico* mutating each CTCF instance in the mouse genome (Fig. 5D) and demonstrating that the Chimaera-predicted maps look much more similar to the map generated in the *in vivo* rapid depletion of CTCF experiment than to the map of untreated cells (64). The ability of Chimaera to connect chromatin features to the underlying DNA suggests that it can be used for *de novo* design of synthetic DNA with the desired Hi-C/Micro-C patterns, akin to *in silico* design of enhancers and cis-regulatory elements (139, 140), promoters (141, 142), and nucleosomal organization (143).

Chimaera is capable of revealing species-specific associations, exemplified by insulation association with BEAF-32, M1BP (detected both by *in silico* mutagenesis and *de novo* search), and Cp190, dCTCF (detected by *in silico* mutagenesis) motifs in fruitly, which are known to act as insulators in this species. We note that these results were achieved by the *D. melanogaster* model on a more narrow and fine-grained window size of 80 Kb, while the model trained on 250 Kb windows did not show any significance of known insulatory motifs. It can be a sign that short DNA-binding motifs of insulators are not the only factor influencing the genome folding of *D. melanogaster*, which might also be genes, nucleotide content, or yet unknown DNA patterns instead.

Chimaera reveals the association of insulation with GC-enrichment in most studied organisms, especially in *M. musculus* and *S. cerevisiae*. The GC-content has been previously associated with promoter regions in vertebrates (117) and gene bodies of yeast (118), which might indicate the importance of active transcription for the formation of three-dimensional structure. While there is no final, universal explanation for the connection between insulation and active transcription, some hypotheses on the potential mechanisms have been suggested (144, 145). In line with that, we observe that *in silico* insertions of genes play a critical role in the 3D organization in most of the studied species (Fig. 7).

Moreover, Chimaera shows not only importance of gene locations but also of their mutual orientation. This effect was observed in two protists, amoeba *D. discoideum* and dinoflagellate *S. microadriaticum*. Despite their maps having very different visual properties (loops in *D. discoideum* versus domains and insulation in *S. microadriaticum*), chromatin organization of both these species, as already reported (37, 81), depends on convergent and divergent gene loci (Fig. 7J, K). Loops in the chromatin of *D. discoideum* are formed at sites of convergent transcription due to RNA polymerase moving along DNA, pushing extruder proteins, and unloading at direct collisions (37).

To explore the associations with other 3D genome patterns, we consider formation of fountains, a recently reported architectural pattern, which looks as enriched interactions emanating from a single genomic locus (14, 15). With Chimaera’s Evolutionary Search, we establish the association between fountains and GC-motifs in *D. rerio* and *A. thaliana*, which have not been previously associated with fountains but can provide important insight for future mechanistic and functional interpretation of fountains.

Notably, the *de novo* motif search methods we implement — Integrated Gradients, Gradient Descent, and Evolutionary Search — do not always yield a single dominant association within the data. These methods can be run independently and complement each other in identifying significant associations. Both Gradient Descent and Evolutionary Search can be launched multiple times (see Methods), potentially producing varying results due to the stochastic nature of each approach, which may reflect the presence of multiple local optima in search for DNA motifs influencing chromatin folding. At that, the Evolutionary Search produces both the DNA motif and the position-specific weight matrix that can be used to estimate the importance of each position.

However, for some organisms, no significant motifs were identified. This could be partially due to the low prediction quality of the models for these species (Suppl. Table 1 and Suppl. Fig. 7). However, despite the high accuracy of the models for amoeba *D. discoideum* or *Hymenoptera* insects (*A. cerana* and *C. hispanica*), no motifs were detected. It suggests that, in these species, the chromatin structure formation may not rely on sequence-specific DNA binding factors such as transcription (*e.g.* in *D. discoideum*, see above).

Since we train the DNA encoder for each species on a whole organism or a single cell type data, we can potentially miss DNA-encoded organization patterns that are present in only a fraction of cells or are specific to unprobed cell types (such as fountains/jets of fungus *Fusarium graminearum* induced upon toxin production (27)). Additionally, some chromatin features arise from the epigenetic state rather than the DNA sequence. In fruitfly, TAD organization reflects and can be modeled from histone modification and transcriptional state, with inactive chromatin aggregating and active chromatin delimiting domains (29, 31). In mouse, oocyte-derived H3K27me3 acts as a maternally inherited, DNA-methylation-independent imprint in early embryos (146). In zebrafish, gametes retain nucleosomal packaging with defined histone PTMs from the parental chromatin (147).

The aim of performed *in silico* mutagenesis is not to predict the consequences of real mutations, which clearly requires information beyond sequence alone, but to interpret the obtained models. For example, the model may change its predictions after inserting only a part of some significant sequence if it has never seen this part separately in the train sample. In this case, the model may not predict the real effect of the mutation but will indicate a possible mechanism. At that point, we observe that mutating CTCF sites *in silico* has similar consequences for the structure prediction as experimental degradation of CTCF.

Finally, we constructed a chromatin tree of life based on Chimaera models trained for different species (Fig. 8). In this approach, each species-trained model serves as a proxy for the mechanism of chromatin organization in the nucleus of this species. The tree obtained from the correlation matrix of cross-predictions generally reflects taxonomic relationships between the studied organisms. We speculate that this reflects the conservation of chromatin-folding mechanisms in evolution and their gradual change with the increasing evolutionary distance between species. At that, the differences in chromatin-based and conventional trees may be due to several reasons, starting from insufficient or low-resolution data and, consequently, variability of the models’ performance. The quality of input Hi-C/MicroC in our analysis guided the selection of the map resolution and the respective field size (see marks next to each species in Fig. 8), which can potentially contribute to the tree clustering. Indeed, we expect that with more consistently obtained Hi-C, Micro-C, or any other 3D genomic assay datasets in multiple species this problem will be mitigated. Moreover, future Chimaera-like models might consider not only 3D genome interactions close to the diagonal, but long-range and inter-chromosomal interactions taking off-diagonal Hi-C signal into account).

Further exploration in the direction of transfer learning between species will allow improvements and, potentially, data imputation for species with lower quality of datasets.

Finally, the misfits in the chromatin tree of life might represent real evolutionary events when the mechanisms governing three-dimensional genome architecture have drastically changed, potentially with no opportunity to establish similarities with other species. In accordance with this hypothesis, the mechanisms of epigenetic regulation are known to vary substantially between species (148).

To sum up, Chimaera is a powerful tool for interpreting chromatin data, providing a new approach to understanding the 3D genome organization, with insights into how evolutionary pressure may have shaped the chromatin architecture. Chimaera utilizes a vast amount of Hi-C and Micro-C datasets published for multiple species, including non-model organisms, and yields a biologically relevant interpretation of associations between 1D and 3D genome patterns. Hence, the present study may inform future studies on the evolution of chromatin folding and the history of structural patterns and structure-based regulation of gene expression.

## ACKNOWLEDGEMENTS

We are grateful to the research groups who shared the preliminary data with us: Prof. Dr. S.V. Razin, his lab and Dr. Sergey Ulianov from IGB RAS (*Dictyostelium discoideum* Hi-C and RNA-Seq and *Danio rerio* Hi-C with fountains), Dr. Daria Onichtchouk from the Freiburg University (*D. rerio* Hi-C with fountains), Prof. Dr. Boris Reizis and Dr. Nicholas Adams from NYU (mouse Hi-C on dendritic cells).

Aleksei Shkolikov and Mikhail Gelfand thank Prof. Andrey A. Mironov for extensive discussions and the idea of building the chromatin-based tree of life. Aleksandra Galitsyna thanks Prof. Leonid Mirny, Prof. Geoff Fudenberg, and Dr. Simon Grosse-Holz for critical discussions at the early stages of the project development. Mikhail Gelfand and Aleksei Shkolikov thank Anastasia Kashtanova for pre-processing some of the Hi-C data and sharing it with the team. Aleksandra Galitsyna and Aleksei Shkolikov thank students Dmitry Skripka, Alexandr Yakushev, and Alexandra Madorskaya for testing Chimaera API.

## AUTHOR CONTRIBUTIONS

Aleksei Shkolikov: conceived the original idea, designed, implemented and tested the model, developed the Chimaera toolkit. Aleksandra Galitsyna: conceived the original idea, wrote the first draft, analysed the data. Mikhail Gelfand: supervised bioinformatics and analysis. All the authors commented and edited the manuscript.

## SUPPLEMENTARY DATA

Supplementary Data are available at NAR online.

## DATA AVAILABILITY

Raw and processed Hi-C and Micro-C data were obtained from BioProject accessions PRJNA606649 (*Xenopus tropicalis*), [PRJNA630123](https://www.ncbi.nlm.nih.gov/bioproject/630123) (*Anopheles merus*), [PRJNA665323](https://www.ncbi.nlm.nih.gov/bioproject/665323) (*Culex quinquefasciatus*), [PRJNA749654](https://www.ncbi.nlm.nih.gov/bioproject/749654) (*Sarcoptes scabiei*), [PRJNA683935](https://www.ncbi.nlm.nih.gov/bioproject/749654) (*Archegozetes longisetosus*), PRJNA680311 (*Arion vulgaris*), [PRJNA427478](https://www.ncbi.nlm.nih.gov/bioproject/?term=PRJNA680311) (*Pomacea canaliculata*), PRJNA792953 (*Cataglyphis hispanica*) PRJCA014302 (*Arabidopsis thaliana*); BioSample accession [SAMN13118423](https://www.ncbi.nlm.nih.gov/biosample/13118423) (*Apis cerana*); GEO datasets GSE178982 (*Mus musculus* ESC), GSE129997 (*Mus musculus* cell cycle), GSE178982 (*Mus musculus* with depleted structural proteins) GSE171396 (*Drosophila melanogaster*), GSE128568 (*Caenorhabditis elegans*), GSE151553 (*Saccharomyces cerevisiae* wild type), GSE217017 (S*accharomyces cerevisiae* with exogenous DNA), GSE85220 (*Schizosaccharomyces pombe*) GSE260572 (*Trichoplax adhaerens, Mnemiopsis leidyi*) GSE152150 (*Symbiodinium microadriaticum*), GSE195609 (*Danio rerio* embryos), GSE134055 (Danio rerio muscle cells) GSE247397 (*Dictyostelium discoideum*), GSM7120275 (Bombyx mori); 4DN dataset [4DNBSZOFFFM6](https://data.4dnucleome.org/experiments-hi-c/4DNEX5F27GL2/) (*Homo sapiens*).

Preprocessed data for all studied organisms and trained models are posted online at OSF (doi: 10.17605/OSF.IO/YF7CR). Chimaera code and illustrative examples are available at Zenodo (doi: 10.5281/zenodo.17418710).

