## Supplementary materials for "Deciphering the 3D genome organization across species from Hi-C data"

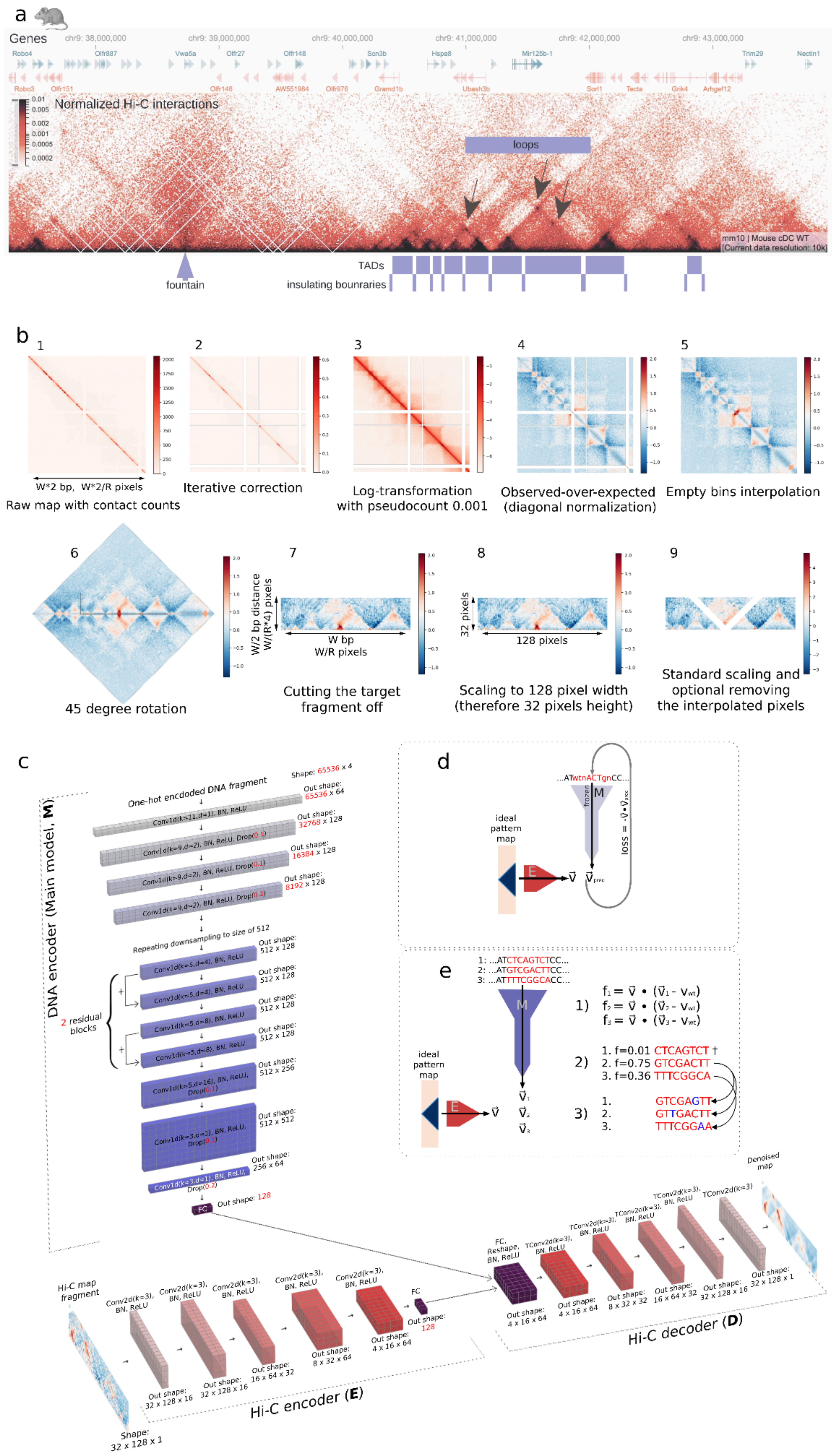

a
Genes
chr9: 38,000,000
chr9: 39,000,000
chr9: 40,000,000
chr9: 41,000,000
chr9: 42,000,000
chr9: 43,000,000
Normalized Hi-C interactions
loops
mm10 | Mouse cDC WT
[Current data resolution: 10k]
TADs
fountain
insulating bounraries
b
1
W*2 bp, W*2/R pixels
Raw map with contact counts
2
Iterative correction
3
Log-transformation with pseudocount 0.001
4
Observed-over-expected (diagonal normalization)
5
Empty bins interpolation
6
45 degree rotation
7
W/2 bp distance W/(R*4) pixels
W bp
W/R pixels
Cutting the target fragment off
8
32 pixels
128 pixels
Scaling to 128 pixel width (therefore 32 pixels height)
9
Standard scaling and optional removing the interpolated pixels
c
DNA encoder (Main model, M)
One-hot encoded DNA fragment
Shape: 65536 x 4
Conv1d(k=21,d=1), BN, ReLU
Out shape: 65536 x 64
Conv1d(k=9,d=2), BN, ReLU, Drop(0.1)
Out shape: 32768 x 128
Out shape: 16384 x 128
Out shape: 8192 x 128
Repeating downsampling to size of 512
2 residual blocks
Conv1d(k=5,d=4), BN, ReLU
Conv1d(k=5,d=8), BN, ReLU
Out shape: 512 x 128
Conv1d(k=5,d=16), BN, ReLU, Drop(0.1)
Out shape: 512 x 256
Conv1d(k=3,d=1), BN, ReLU, Drop(0.2)
Out shape: 512 x 512
Out shape: 256 x 64
FC
Out shape: 128
Hi-C map fragment
Shape: 32 x 128 x 1
Conv2d(k=3), BN, ReLU
Out shape: 32 x 128 x 16
Out shape: 16 x 64 x 32
Out shape: 8 x 32 x 64
Out shape: 4 x 16 x 64
Hi-C encoder (E)
FC, Reshape, BN, ReLU
TConv2d(k=3), BN, ReLU
Out shape: 4 x 16 x 64
Out shape: 8 x 32 x 32
Out shape: 16 x 64 x 32
Out shape: 32 x 128 x 16
Out shape: 32 x 128 x 1
Denoised map
Hi-C decoder (D)
d
...ATwtnACTgnCC...
frozen
M
ideal pattern map
E
loss = -v⃗•v⃗pred
e
1: ...ATCTCAGTCTCC...
2: ...ATGTCGACTTCC...
3: ...ATTTTCGGCACC...
M
ideal pattern map
E
1) f1 = v⃗ • (v⃗1 - vwt)
f2 = v⃗ • (v⃗2 - vwt)
f3 = v⃗ • (v⃗3 - vwt)
2) 1. f=0.01 CTCAGTCT †
2. f=0.75 GTCGACTT
3. f=0.36 TTTCGGCA
3) 1. GTCGAGTT
2. GTTGACTT
3. TTTCGGAA

**Supplementary Figure 1.**

**(A)** Representative Hi-C map and typical patterns: fountain/jet, TADs, insulating boundaries, loops. Here, we show a ~6 Mb region of chr9 of mouse conventional dendritic cells (cDCs) from (1) that contains all these features. The resolution of the Hi-C map is 10 Kb (mm10 genome assembly). Representative patterns were selected manually for representation purposes. Gene annotation is shown above the Hi-C map.

**Fountain (**often called *jet* (2), *flare* (3), or *hairpin* (4)) is an off-diagonal ridge in a Hi-C/Micro-C map that emanates from a single locus on the main diagonal and extends for hundreds of kb up to 1–2 Mb. Fountains can differ in width (5), length (2, 5), and angle of the ridge (2).
**TAD (topologically associating domain)** is a contiguous genomic interval within which chromatin contacts are enriched relative to regions outside the interval (6, 7). On a tilted Hi-C/Micro-C map (as we use throughout this work), it appears as a triangle along the first line. In a square representation of Hi-C map, it looks like a square along the diagonal. There is an active debate on what the TADs are and how to define them (8).

**Insulation** is the depletion of interactions between two genomic areas that are separated by the *insulation boundary* (9). Typically, insulation boundaries are located at the borders of TADs.

**Loop (dot)** is an enriched interaction between pair of genomic loci, visible as an off-diagonal bright dot (10).

In this work, we also consider **stripes** (enriched interactions of a single genomic locus with neighboring regions (11, 12), not shown on the representative Hi-C map). Typically for mammals, stripes are located at the TAD edges, and have the same epigenetic/genomic signature as loops and TAD boundaries. Thus, we do not investigate their formation further.

**Compartments** (not shown on the representative Hi-C map) are alternating domains of enriched interactions spanning whole chromosomes and whole genomes (13). Compartments are visible in the Hi-C/Micro-C maps as blocks of off-diagonal enriched interactions, which can be broad (0.5-1 Mb (13) and ~12.5 kb in more recent studies (14) in mammals, ~15 kb in fruitfly (15), ~80-120 kb in silkmoth (16)) or very narrow (several Kb-microcompartments (17)). In this work, we focus on patterns close to the diagonal (300-500 Kb away), the length scale comparable to large classical compartment size, and do not study the compartments. However, we acknowledge that some loops might be in fact classical- or micro-compartments, and some TADs might be in fact compartmental domains (18).

**(B)** Hi-C windows preprocessing pipeline. W – target window size (in base pairs), R – original data resolution.

**(C-E)** Chimaera architecture (C) and interpretation methods (D, E)**.** E – Hi-C Encoder, D – Hi-C Decoder, M – DNA Encoder, or Main model. frozen – the weights of the model block are fixed and not updated at this step. **(C)** Model architecture. Parameters in black: the same for all species. Parameters in red: parameters that we varied and selected depending on the size of the input DNA/Hi-C fragment (see

Table 3), and/or sample size. Conv1d/Conv2d – convolutional layers, TConv – transposed convolutional layer, BN – batch normalization, FC – fully connected, ReLU – rectified linear unit. Between convolutional layers we have used max pooling (see the change in dimensionality of the convolutional blocks). **(D, E)** Chimaera-based *de novo* calling of DNA motifs corresponding to a given feature: two approaches that we provide together with Chimaera, the given example is the insulation pattern search. **(D)** Gradient Descent in the space of DNA. We maximize the projection of the embedded vector ($v_{pred}$) onto a given interaction pattern (v) in the successive iterations of the model run on DNA sequence. The weights of both Hi-C and DNA encoders are frozen, and only the input DNA sequence is changed. **(E)** Evolutionary Search in the space of DNA. As an input, we generate a set of random DNA sequences, propagate them through the DNA encoder, and then iteratively run steps (1-3): 1) — calculate the fitness function, 2) — select the best DNA sequences and let them reproduce, 3) — randomly mutate the reproduced sequences.

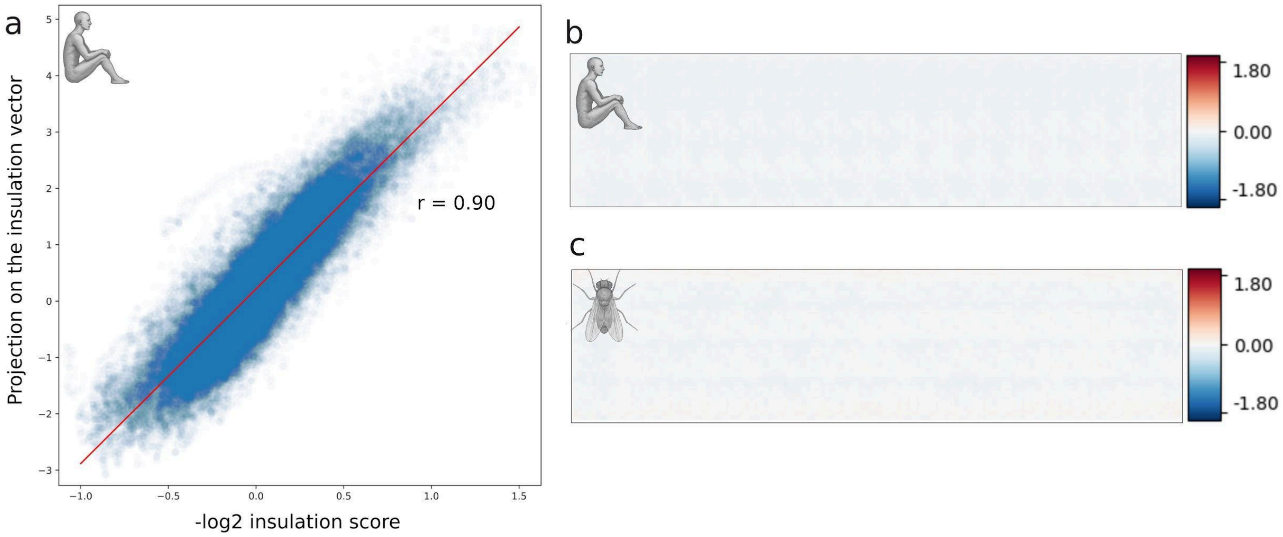


**Supplementary Figure 2.**

**Properties of the latent space.** Supporting information for main Fig. 2.

**(A)** Correlation plot of the projection onto the insulation vector against the insulation score. We plot sign-flipped values of the traditional insulation scores (9), so that positive values of (-log (insulation scores)) correspond to more pronounced insulation patterns. Insulation scores were calculated for the whole genome by *cooltools (19)*. **(B-C)** Decoded Hi-C map corresponding to the zero vector in the latent space. Patterns are absent; note that the scale of the colorbar is much smaller than for main Fig. 2A-B. **(B)** For the human model. **(C)** For the *D. melanogaster* model.

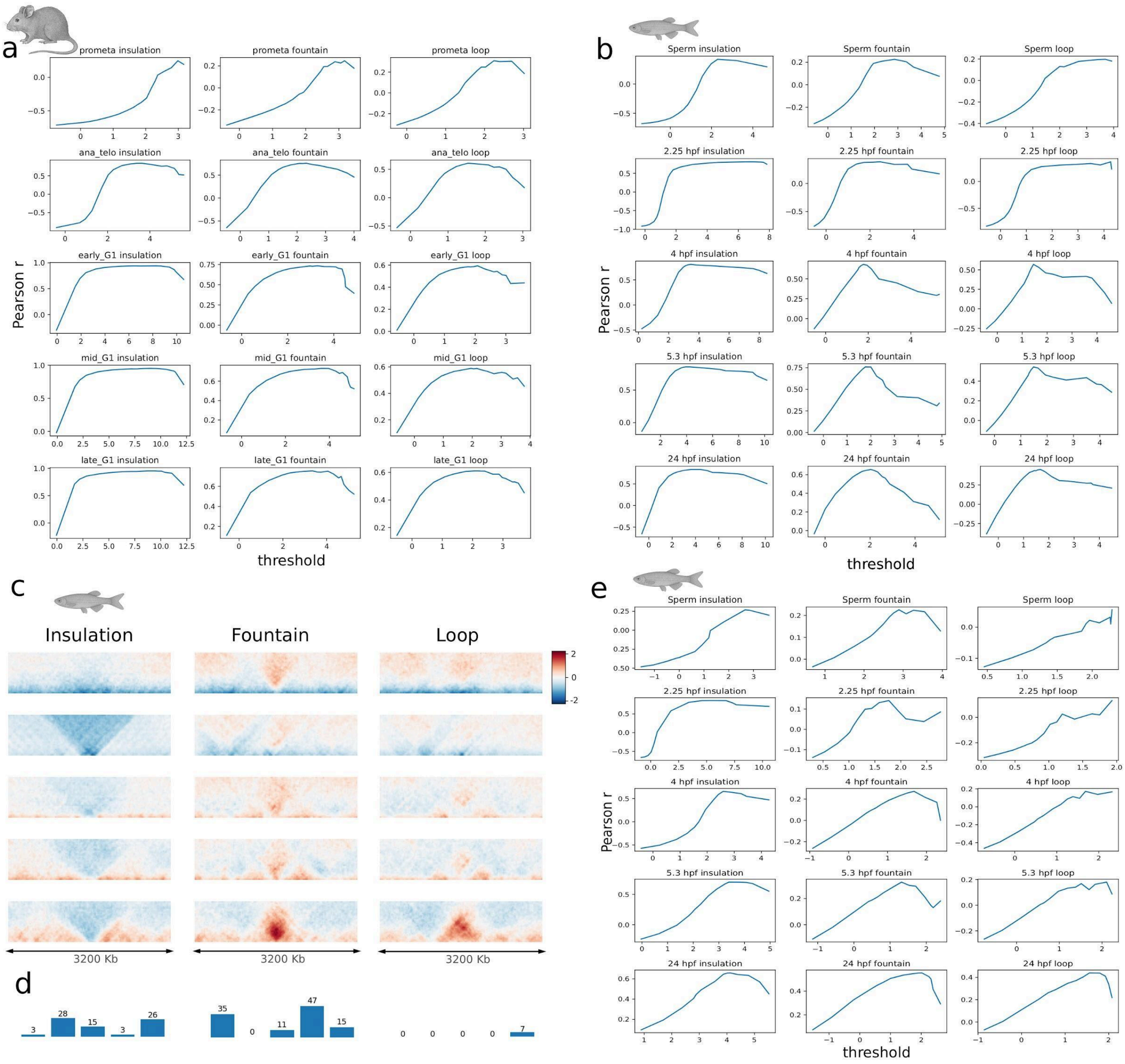


**Supplementary Figure 3.**

**Feature calling.** Supporting information for Figure 3.

**(A–B)** Threshold selection plots for Hi-C pattern calling in main Fig. 3.

**(A)** *M. musculus* cell cycle (500 Kb windows), **(B)** *D. rerio* embryogenesis (500 Kb windows). X axis represents thresholds, Y axis is the Pearson correlation coefficient between the pattern template and the preprocessed averaged map obtained from a sample selected using this threshold (see Methods, "*Pattern calling in Hi-C maps*"). Most graphs have only one maximum.

**(C-E)** Feature calling for *Danio rerio* with increased (3.2 Mb) window size. This way, we aimed to capture large-scale fountains (flares) observed previously in sperm cells (central top). The size of the features here is increased proportionally to the increase of the window size.

**(C)** Average pileup of top 0.5% best findings of insulation, fountain, and loop patterns using latent space projections for *D. rerio* embryogenesis (3.2 Mb windows).

(**D**) Numbers of 25 Kb bins with projection larger than selected thresholds for each stage.

**(E)** Threshold selection for (D), similar to (A) and (B).

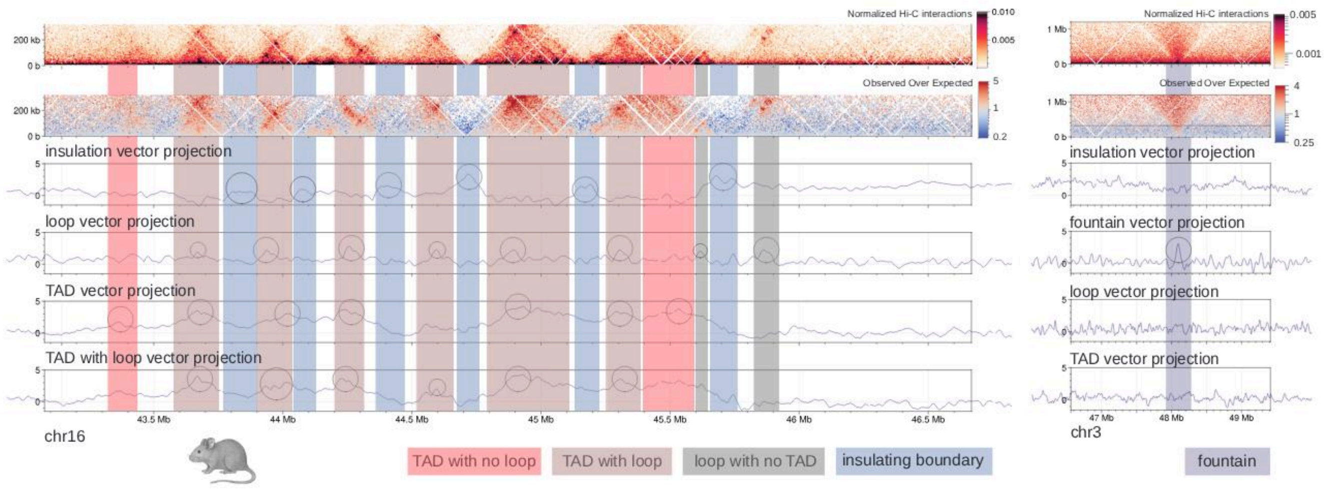


**Supplementary Figure 4.**

**Feature calling in Hi-C maps with Chimaera autoencoder.**

Illustrative feature calling (insulation, TADs and loops (left), and fountain (right)) for segments of mouse conventional dendritic cells (cDCs) Hi-C data from (1). Multiple features may appear at the same location and be called by Chimaera (e.g. TAD with loop).

**Top to bottom:** normalized Hi-C map, observed over expected Hi-C map, Chimaera-produced tracks of projections to the pattern templates. Circles represent elevations in the tracks (manual demarcation). Vertical boxes represent the called regions (manual demarcation).

**(Left)** Hi-C map at the 5 Kb resolution.

**(Right)** Hi-C map at 10 Kb resolution. Horizontal line represents maximal receptive distance of Chimaera (320 Kb in this case).

Note that the Chimaera autoencoder is agnostic to the mechanism of formation of the structures. Thus, some of the detected loops might be some loops might be in fact classical- or micro-compartments (14, 17), and some TADs might be in fact compartmental domains (Rao et al. 2017).

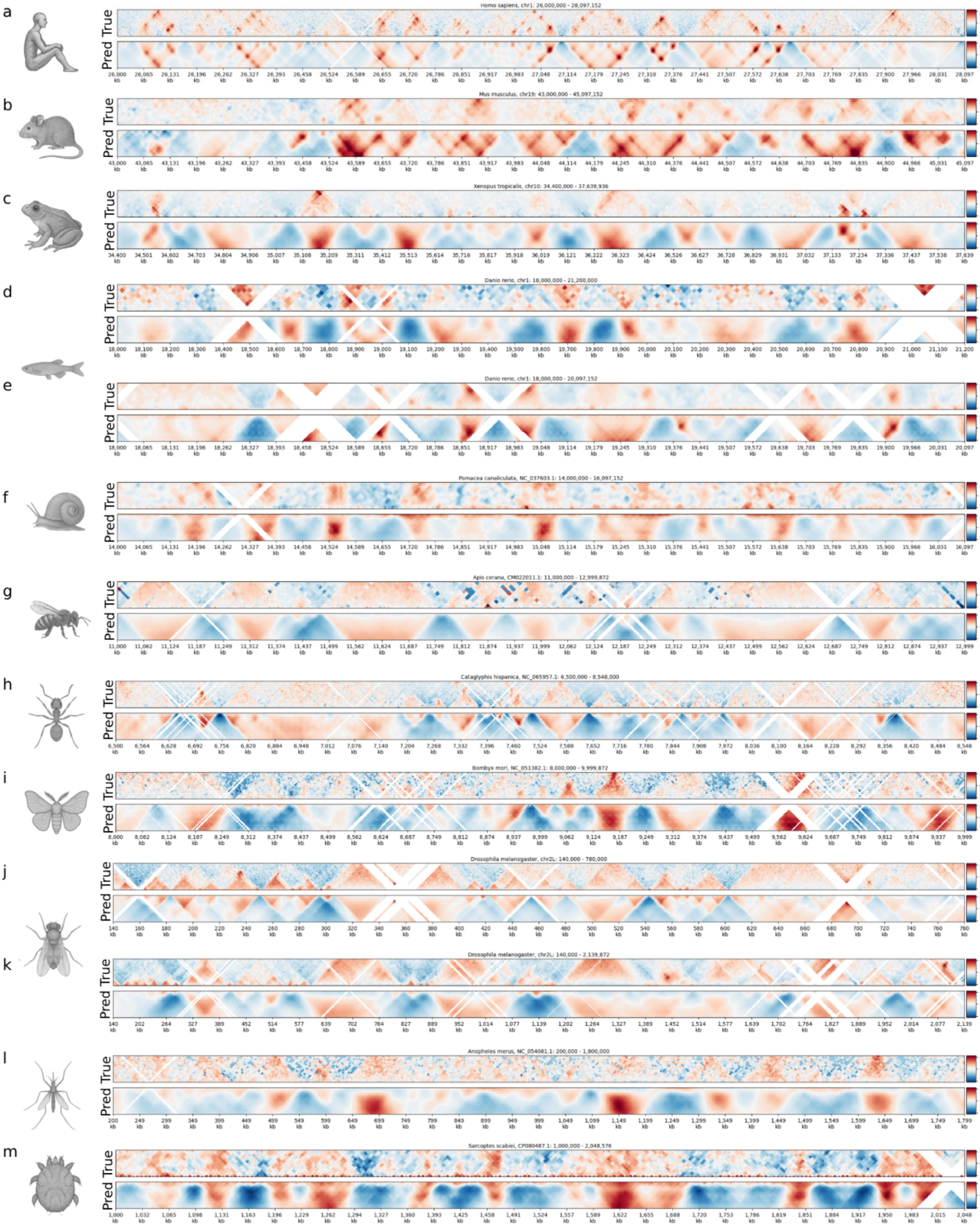
Homo sapiens, chr1: 26,000,000 - 28,097,152
Mus musculus, chr19: 43,000,000 - 45,097,152
Xenopus tropicalis, chr10: 34,400,000 - 37,639,936
Danio rerio, chr1: 18,000,000 - 21,200,000
Danio rerio, chr1: 18,000,000 - 20,097,152
Pomacea canaliculata, NC_037603.1: 14,000,000 - 16,097,152
Apis cerana, CM022011.1: 11,000,000 - 12,999,872
Cataglyphis hispanica, NC_065957.1: 6,500,000 - 8,548,000
Bombyx mori, NC_051382.1: 8,000,000 - 9,999,872
Drosophila melanogaster, chr2L: 140,000 - 780,000
Drosophila melanogaster, chr2L: 140,000 - 2,139,872
Anopheles merus, NC_054081.1: 200,000 - 1,800,000
Sarcoptes scabiei, CP080487.1: 1,000,000 - 2,048,576
True
Pred

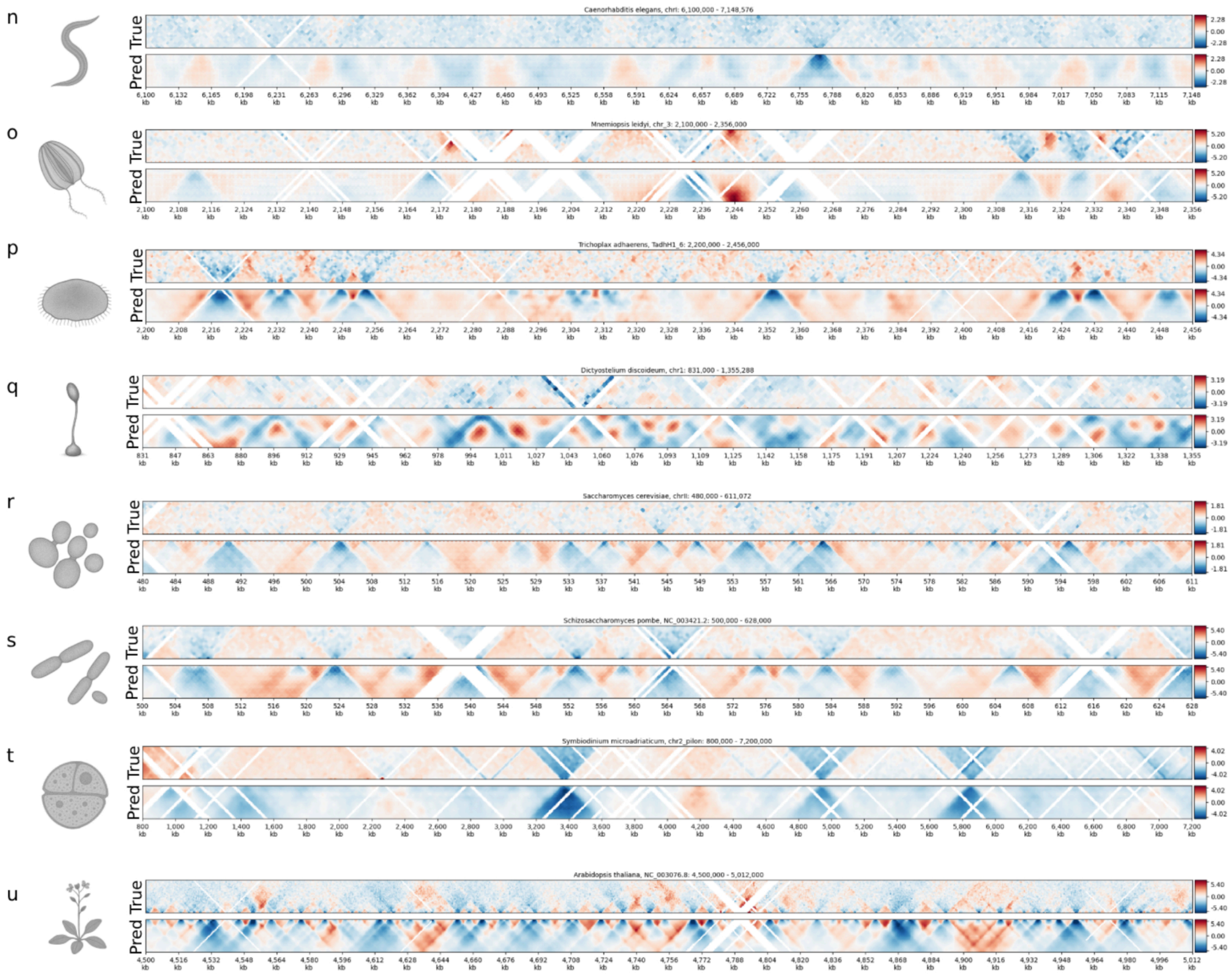


**Supplementary Figure 5.**

**Examples of Hi-C predicted maps based on DNA sequence for the studied species .** The top of each panel is a true input map, the bottom is a mirrored predicted one. From top to bottom: (A) Micro-C for *Homo sapiens* HFFc6 cell line from (20), (B) Micro-C for *Mus musculus* embryonic stem cells from (21), (C) Hi-C for *Xenopus tropicalis* embryos from (22), for *Danio rerio*: for embryos at 5.3 hours past fertilization (D) from (5) and for muscle cells (E) from (23), (F) for  snail *Pomacea canaliculata* from (24), (G) for bee *Apis cerana* from the drone pupae from (25), (H) for ant *Cataglyphis hispanica* from (26), (I) for butterfly *Bombyx mori* from (16), (J, K) Micro-C for *Drosophila melanogaster* (J is for 80 kb window predictions, K is for 250 kb window predictions) for nc14 stage embryos from (27), (L) Hi-C for mosquitoes *Anopheles merus* from (28), (M) for mite *Sarcoptes scabiei* from (29), (N) Hi-C for nematode *Caenorhabditis elegans* from (30), (O) for comb jelly *Mnemiopsis leidyi* from (31), (P) for placozoan *Trichoplax adhaerens* from (31), (Q) for social amoeba *Dictyostelium discoideum* from (32), (R) Micro-C

XL for yeast *Saccharomyces cerevisiae* at 90 minutes after release from G1 from (33), (S) for fission yeast *Schizosaccharomyces pombe* from (34), (T) Hi-C for dinoflagellate *Symbiodinium microadriaticum* from (35), and (U) Micro-C for plant *Arabidopsis thaliana* from (36).(34)

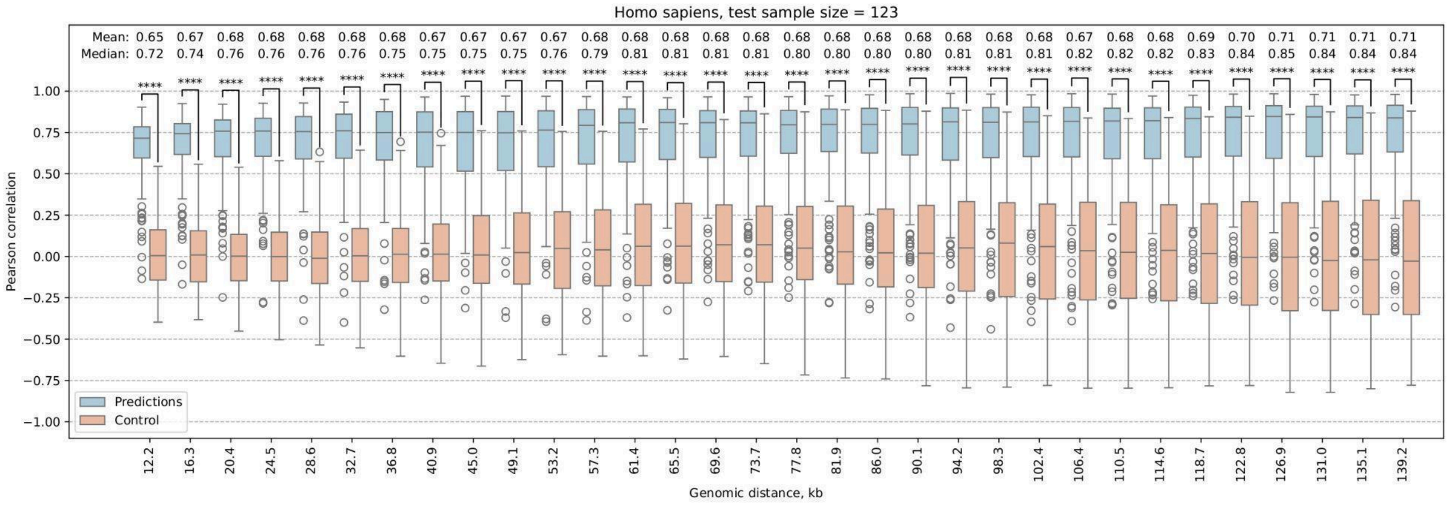
Homo sapiens, test sample size = 123
Mean: 0.65 0.67 0.68 0.68 0.68 0.68 0.68 0.67 0.67 0.67 0.67 0.68 0.68 0.68 0.68 0.68 0.68 0.68 0.68 0.68 0.68 0.68 0.68 0.67 0.68 0.68 0.69 0.70 0.71 0.71 0.71 0.71
Median: 0.72 0.74 0.76 0.76 0.76 0.76 0.75 0.75 0.75 0.75 0.76 0.79 0.81 0.81 0.81 0.81 0.80 0.80 0.80 0.80 0.81 0.81 0.81 0.82 0.82 0.82 0.83 0.84 0.85 0.84 0.84 0.84
Pearson correlation
Predictions
Control
12.2 16.3 20.4 24.5 28.6 32.7 36.8 40.9 45.0 49.1 53.2 57.3 61.4 65.5 69.6 73.7 77.8 81.9 86.0 90.1 94.2 98.3 102.4 106.4 110.5 114.6 118.7 122.8 126.9 131.0 135.1 139.2
Genomic distance, kb

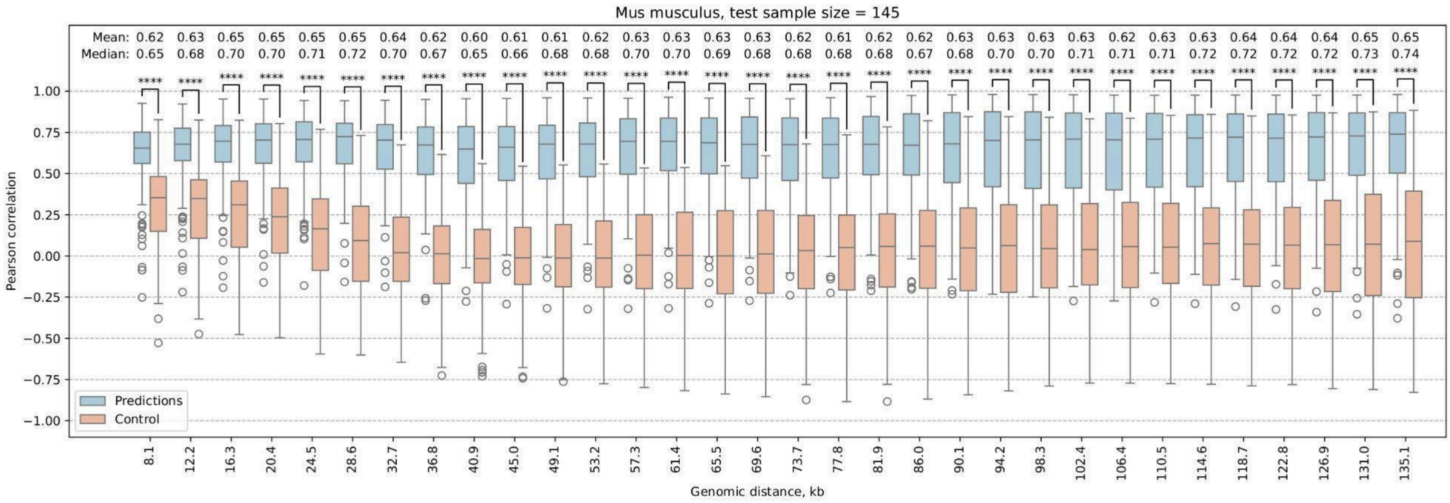
Mus musculus, test sample size = 145
Mean: 0.62 0.63 0.65 0.65 0.65 0.65 0.64 0.62 0.60 0.61 0.61 0.62 0.63 0.63 0.63 0.63 0.62 0.61 0.62 0.62 0.63 0.63 0.63 0.63 0.62 0.63 0.63 0.64 0.64 0.64 0.65 0.65
Median: 0.65 0.68 0.70 0.70 0.71 0.72 0.70 0.67 0.65 0.66 0.68 0.68 0.70 0.70 0.69 0.68 0.68 0.68 0.68 0.67 0.68 0.70 0.70 0.71 0.71 0.71 0.72 0.72 0.72 0.72 0.73 0.74
Pearson correlation
Predictions
Control
8.1 12.2 16.3 20.4 24.5 28.6 32.7 36.8 40.9 45.0 49.1 53.2 57.3 61.4 65.5 69.6 73.7 77.8 81.9 86.0 90.1 94.2 98.3 102.4 106.4 110.5 114.6 118.7 122.8 126.9 131.0 135.1
Genomic distance, kb

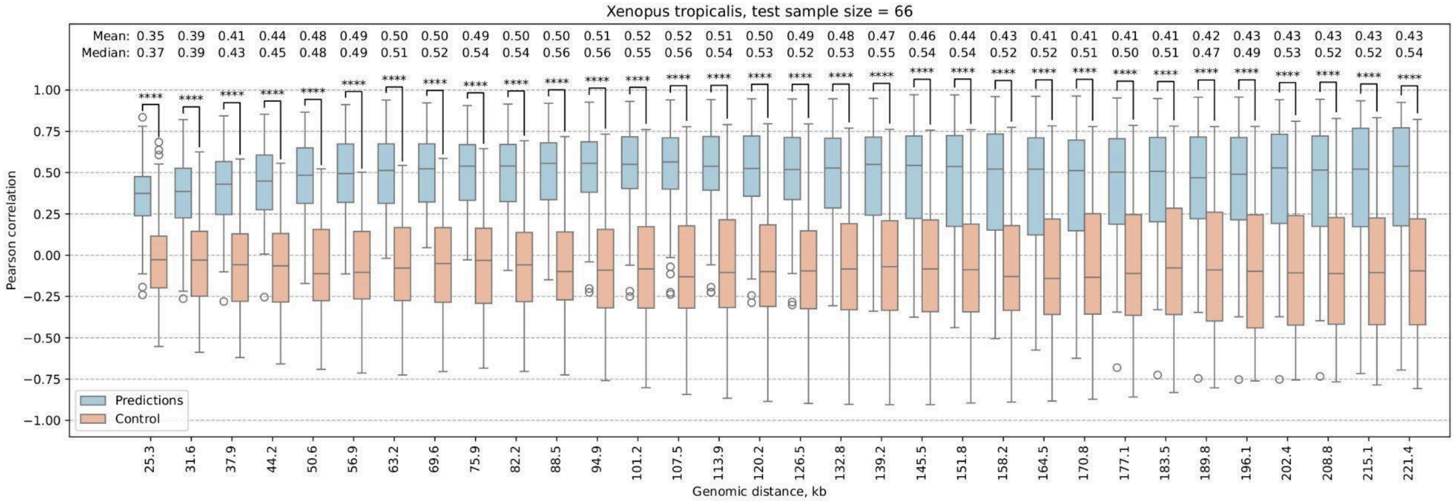
Xenopus tropicalis, test sample size = 66
Mean: 0.35 0.39 0.41 0.44 0.48 0.49 0.50 0.50 0.49 0.50 0.50 0.51 0.52 0.52 0.51 0.50 0.49 0.48 0.47 0.46 0.44 0.43 0.41 0.41 0.41 0.41 0.42 0.43 0.43 0.43 0.43 0.43
Median: 0.37 0.39 0.43 0.45 0.48 0.49 0.51 0.52 0.54 0.54 0.56 0.56 0.55 0.56 0.54 0.53 0.52 0.53 0.55 0.54 0.54 0.52 0.52 0.51 0.50 0.51 0.47 0.49 0.53 0.52 0.52 0.54
Pearson correlation
Predictions
Control
25.3 31.6 37.9 44.2 50.6 56.9 63.2 69.6 75.9 82.2 88.5 94.9 101.2 107.5 113.9 120.2 126.5 132.8 139.2 145.5 151.8 158.2 164.5 170.8 177.1 183.5 189.8 196.1 202.4 208.8 215.1 221.4
Genomic distance, kb

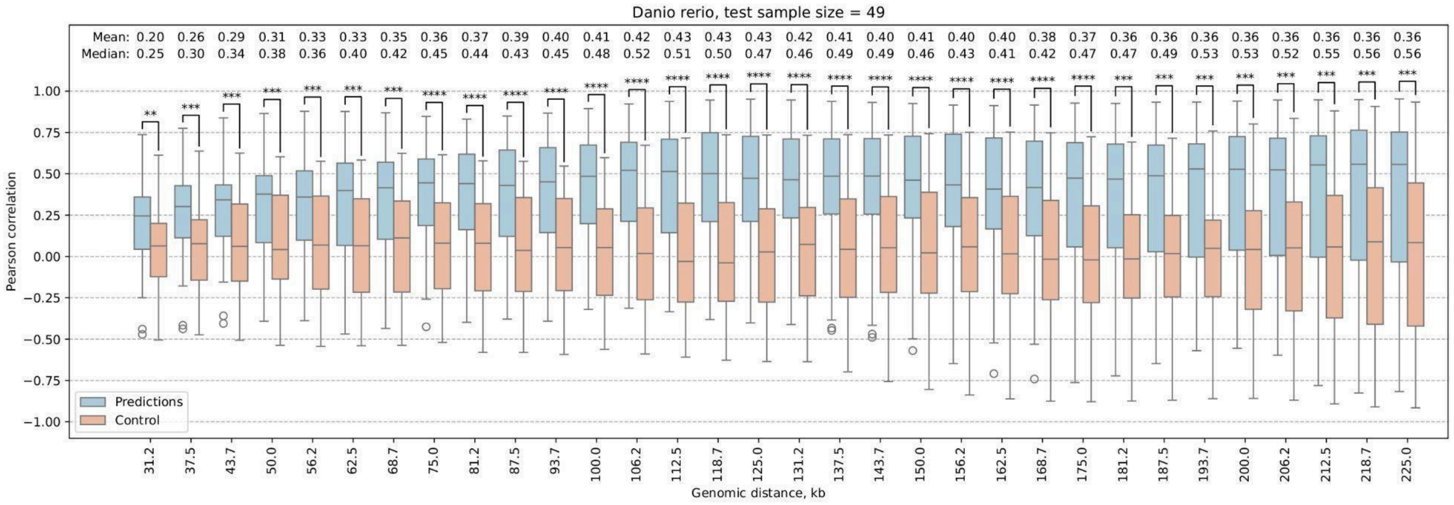
Danio rerio, test sample size = 49
Mean: 0.20 0.26 0.29 0.31 0.33 0.33 0.35 0.36 0.37 0.39 0.40 0.41 0.42 0.43 0.43 0.43 0.42 0.41 0.40 0.41 0.40 0.40 0.38 0.37 0.36 0.36 0.36 0.36 0.36 0.36 0.36 0.36
Median: 0.25 0.30 0.34 0.38 0.36 0.40 0.42 0.45 0.44 0.43 0.45 0.48 0.52 0.51 0.50 0.47 0.46 0.49 0.49 0.46 0.43 0.41 0.42 0.47 0.47 0.49 0.53 0.53 0.52 0.55 0.56 0.56
Pearson correlation
Predictions
Control
Genomic distance, kb

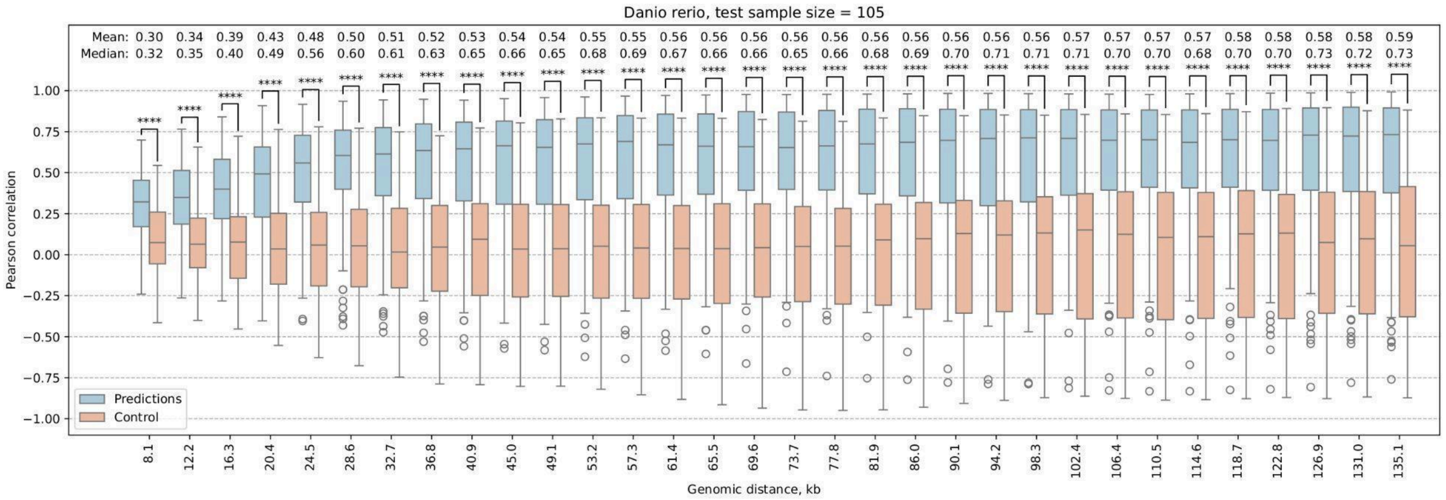
Danio rerio, test sample size = 105
Mean: 0.30 0.34 0.39 0.43 0.48 0.50 0.51 0.52 0.53 0.54 0.54 0.55 0.55 0.56 0.56 0.56 0.56 0.56 0.56 0.56 0.56 0.56 0.56 0.57 0.57 0.57 0.57 0.58 0.58 0.58 0.58 0.59
Median: 0.32 0.35 0.40 0.49 0.56 0.60 0.61 0.63 0.65 0.66 0.65 0.68 0.69 0.67 0.66 0.66 0.65 0.66 0.68 0.69 0.70 0.71 0.71 0.71 0.70 0.70 0.68 0.70 0.70 0.73 0.72 0.73
Pearson correlation
Predictions
Control
Genomic distance, kb

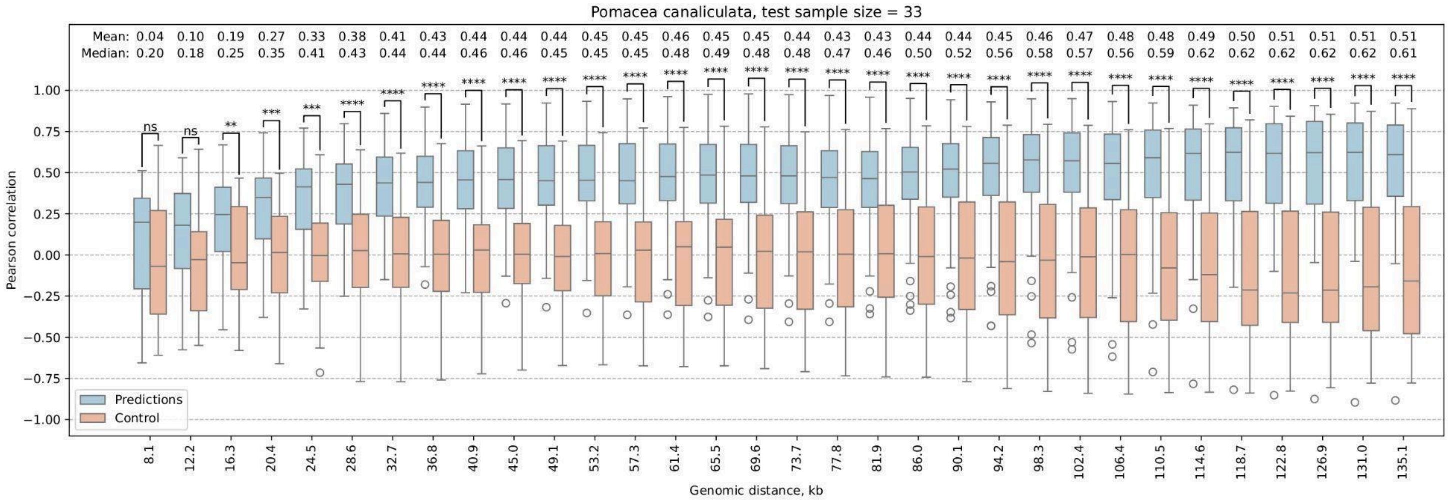
Pomacea canaliculata, test sample size = 33
Mean: 0.04 0.10 0.19 0.27 0.33 0.38 0.41 0.43 0.44 0.44 0.44 0.45 0.45 0.46 0.45 0.45 0.44 0.43 0.43 0.44 0.44 0.45 0.46 0.47 0.48 0.48 0.49 0.50 0.51 0.51 0.51 0.51
Median: 0.20 0.18 0.25 0.35 0.41 0.43 0.44 0.44 0.46 0.46 0.45 0.45 0.45 0.48 0.49 0.48 0.48 0.47 0.46 0.50 0.52 0.56 0.58 0.57 0.56 0.59 0.62 0.62 0.62 0.62 0.62 0.61
Pearson correlation
Predictions
Control
Genomic distance, kb

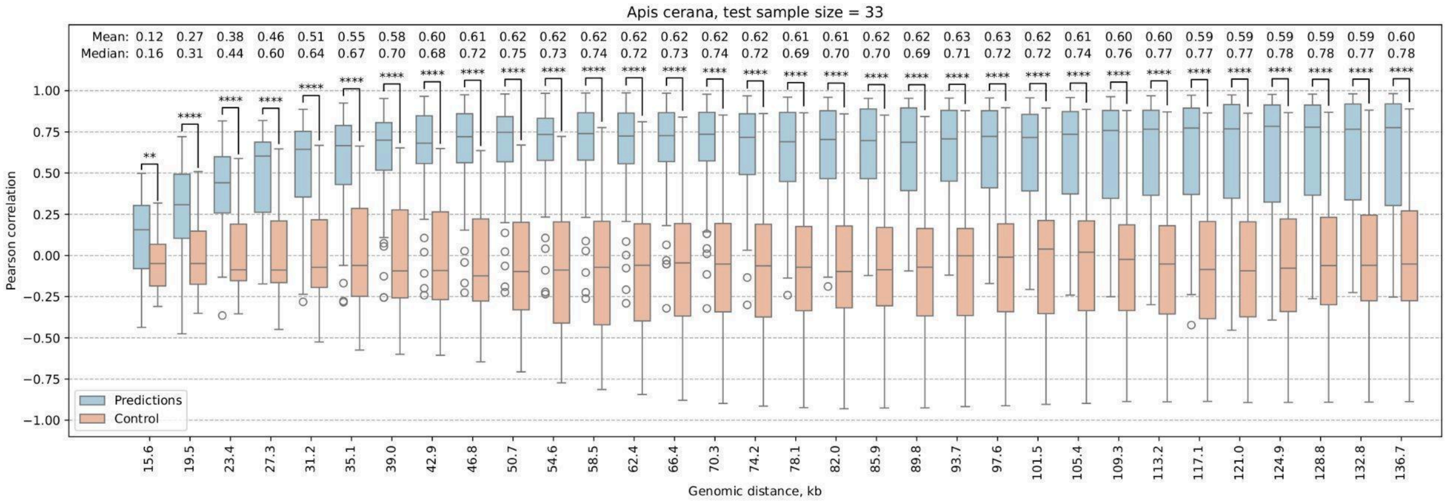
Apis cerana, test sample size = 33
Mean: 0.12 0.27 0.38 0.46 0.51 0.55 0.58 0.60 0.61 0.62 0.62 0.62 0.62 0.62 0.62 0.62 0.61 0.61 0.62 0.62 0.63 0.63 0.62 0.61 0.60 0.60 0.59 0.59 0.59 0.59 0.59 0.60
Median: 0.16 0.31 0.44 0.60 0.64 0.67 0.70 0.68 0.72 0.75 0.73 0.74 0.72 0.73 0.74 0.72 0.69 0.70 0.70 0.69 0.71 0.72 0.72 0.74 0.76 0.77 0.77 0.77 0.78 0.78 0.77 0.78
Pearson correlation
Predictions
Control
Genomic distance, kb

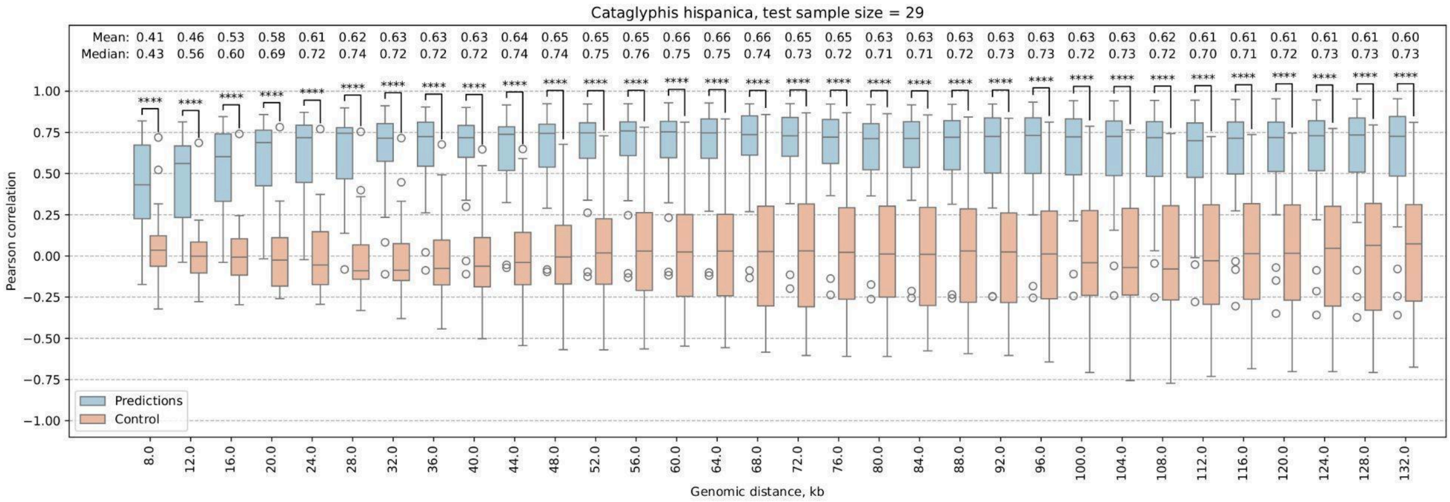
Cataglyphis hispanica, test sample size = 29
Mean: 0.41 0.46 0.53 0.58 0.61 0.62 0.63 0.63 0.63 0.64 0.65 0.65 0.65 0.66 0.66 0.66 0.65 0.65 0.63 0.63 0.63 0.63 0.63 0.63 0.63 0.62 0.61 0.61 0.61 0.61 0.61 0.60
Median: 0.43 0.56 0.60 0.69 0.72 0.74 0.72 0.72 0.72 0.74 0.74 0.75 0.76 0.75 0.75 0.74 0.73 0.72 0.71 0.71 0.72 0.73 0.73 0.72 0.73 0.72 0.70 0.71 0.72 0.73 0.73 0.73
Pearson correlation
Predictions
Control
Genomic distance, kb

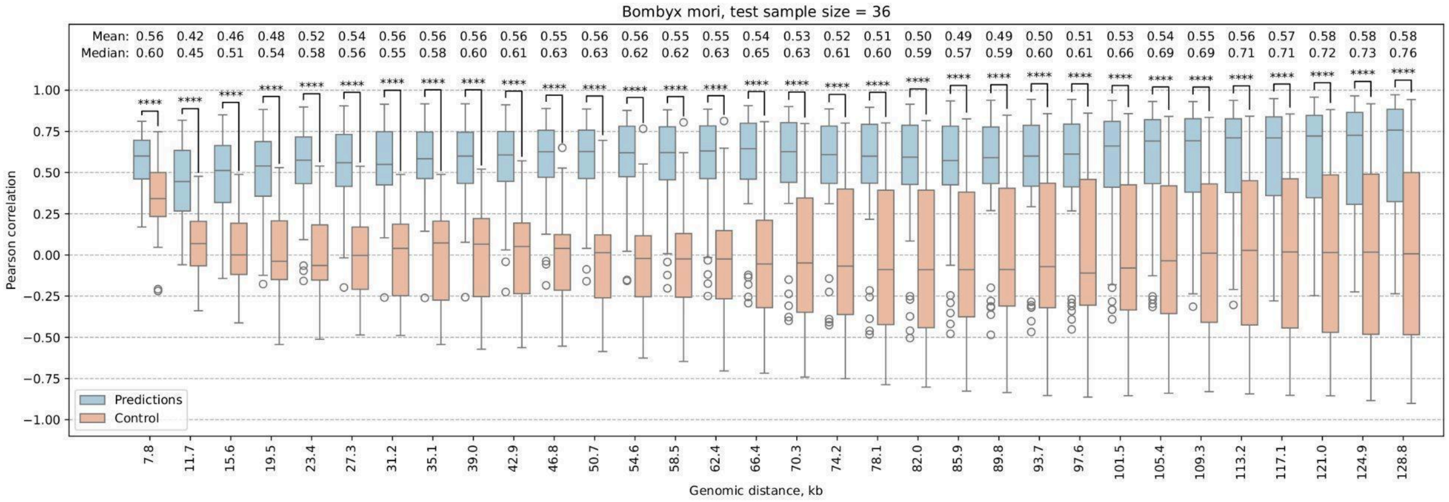
Bombyx mori, test sample size = 36
Mean: 0.56 0.42 0.46 0.48 0.52 0.54 0.56 0.56 0.56 0.56 0.55 0.56 0.56 0.55 0.55 0.54 0.53 0.52 0.51 0.50 0.49 0.49 0.50 0.51 0.53 0.54 0.55 0.56 0.57 0.58 0.58 0.58
Median: 0.60 0.45 0.51 0.54 0.58 0.56 0.55 0.58 0.60 0.61 0.63 0.63 0.62 0.62 0.63 0.65 0.63 0.61 0.60 0.59 0.57 0.59 0.60 0.61 0.66 0.69 0.69 0.71 0.71 0.72 0.73 0.76
Pearson correlation
Predictions
Control
Genomic distance, kb

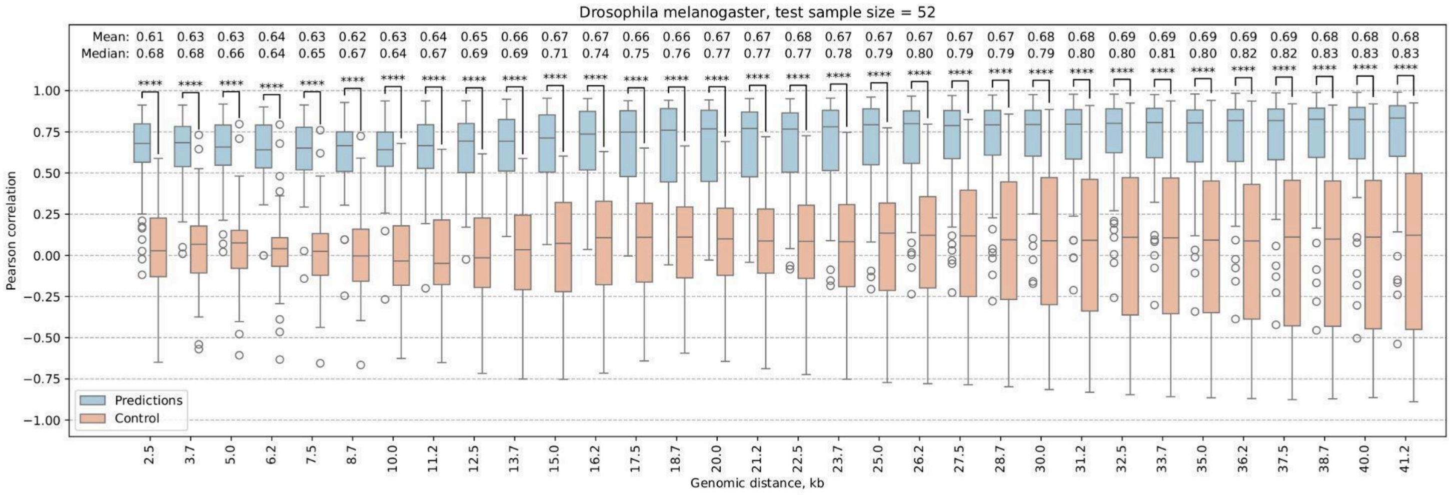
Drosophila melanogaster, test sample size = 52
Mean: 0.61 0.63 0.63 0.64 0.63 0.62 0.63 0.64 0.65 0.66 0.67 0.67 0.66 0.66 0.67 0.67 0.68 0.67 0.67 0.67 0.67 0.67 0.68 0.68 0.69 0.69 0.69 0.69 0.69 0.68 0.68 0.68
Median: 0.68 0.68 0.66 0.64 0.65 0.67 0.64 0.67 0.69 0.69 0.71 0.74 0.75 0.76 0.77 0.77 0.77 0.78 0.79 0.80 0.79 0.79 0.79 0.80 0.80 0.81 0.80 0.82 0.82 0.83 0.83 0.83
Pearson correlation
Genomic distance, kb
Predictions
Control

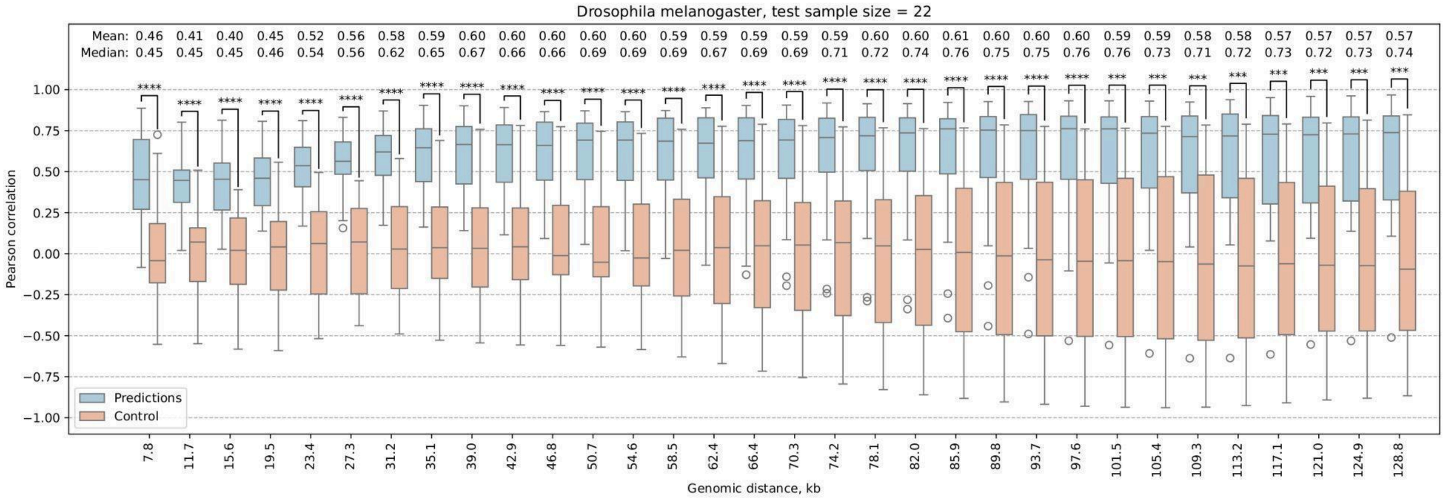
Drosophila melanogaster, test sample size = 22
Mean: 0.46 0.41 0.40 0.45 0.52 0.56 0.58 0.59 0.60 0.60 0.60 0.60 0.60 0.59 0.59 0.59 0.59 0.59 0.60 0.60 0.61 0.60 0.60 0.60 0.59 0.59 0.58 0.58 0.57 0.57 0.57 0.57
Median: 0.45 0.45 0.45 0.46 0.54 0.56 0.62 0.65 0.67 0.66 0.66 0.69 0.69 0.69 0.67 0.69 0.69 0.71 0.72 0.74 0.76 0.75 0.75 0.76 0.76 0.73 0.71 0.72 0.73 0.72 0.73 0.74
Pearson correlation
Genomic distance, kb
Predictions
Control

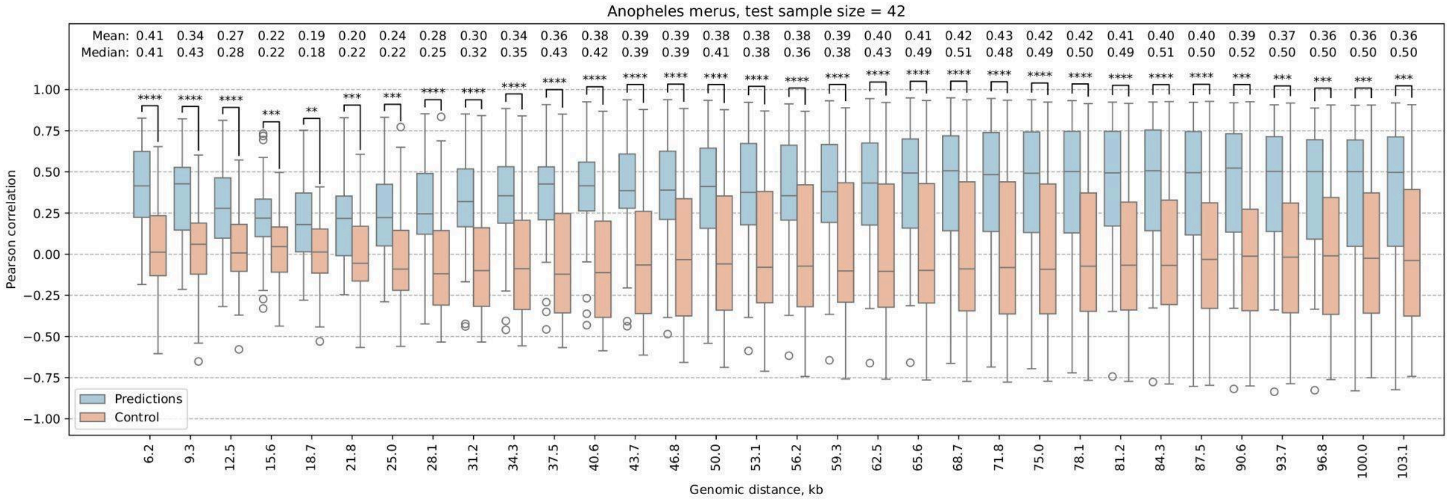
Anopheles merus, test sample size = 42
Mean: 0.41 0.34 0.27 0.22 0.19 0.20 0.24 0.28 0.30 0.34 0.36 0.38 0.39 0.39 0.38 0.38 0.38 0.39 0.40 0.41 0.42 0.43 0.42 0.42 0.41 0.40 0.40 0.39 0.37 0.36 0.36 0.36
Median: 0.41 0.43 0.28 0.22 0.18 0.22 0.22 0.25 0.32 0.35 0.43 0.42 0.39 0.39 0.41 0.38 0.36 0.38 0.43 0.49 0.51 0.48 0.49 0.50 0.49 0.51 0.50 0.52 0.50 0.50 0.50 0.50
Pearson correlation
Genomic distance, kb
Predictions
Control

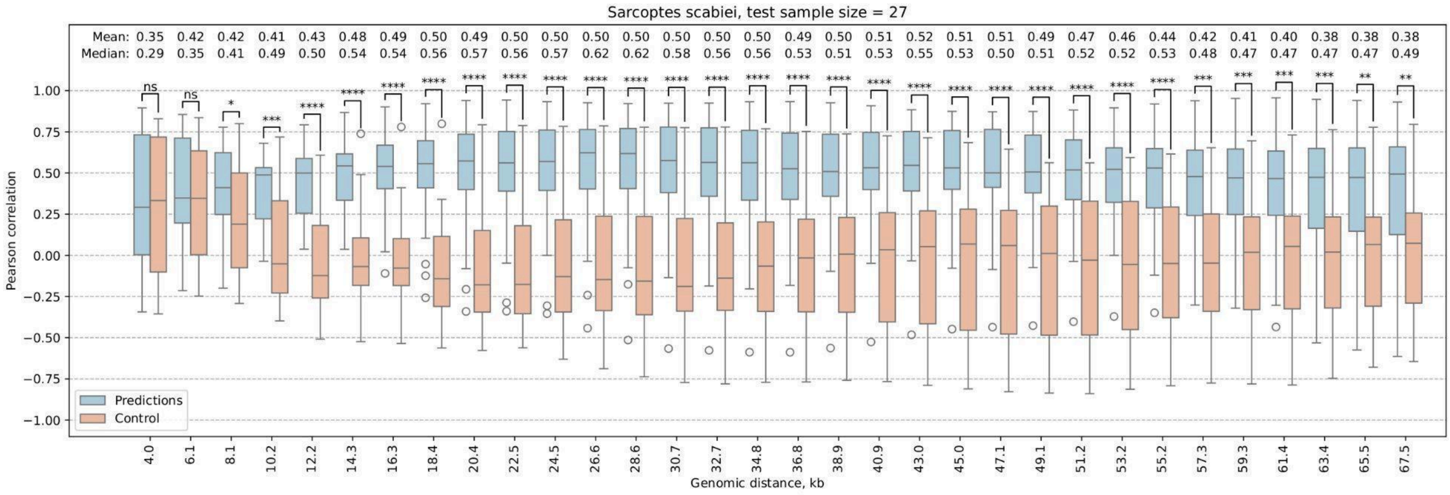
Sarcoptes scabiei, test sample size = 27
Mean: 0.35 0.42 0.42 0.41 0.43 0.48 0.49 0.50 0.49 0.50 0.50 0.50 0.50 0.50 0.50 0.50 0.49 0.50 0.51 0.52 0.51 0.51 0.49 0.47 0.46 0.44 0.42 0.41 0.40 0.38 0.38 0.38
Median: 0.29 0.35 0.41 0.49 0.50 0.54 0.54 0.56 0.57 0.56 0.57 0.62 0.62 0.58 0.56 0.56 0.53 0.51 0.53 0.55 0.53 0.50 0.51 0.52 0.52 0.53 0.48 0.47 0.47 0.47 0.47 0.49
Pearson correlation
Predictions
Control
Genomic distance, kb

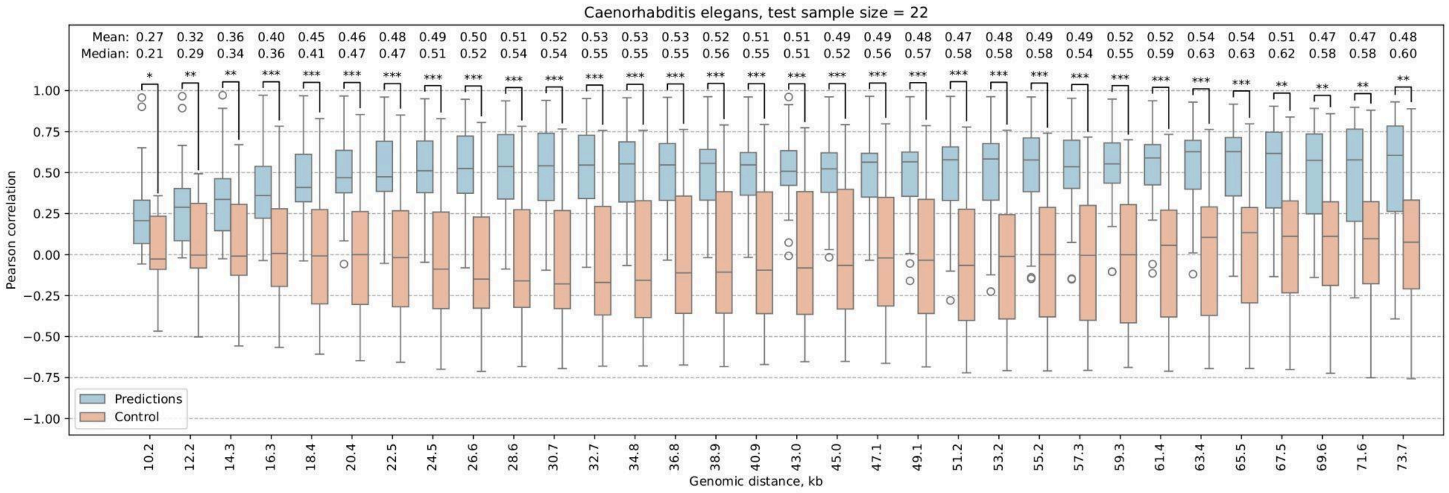
Caenorhabditis elegans, test sample size = 22
Mean: 0.27 0.32 0.36 0.40 0.45 0.46 0.48 0.49 0.50 0.51 0.52 0.53 0.53 0.53 0.52 0.51 0.51 0.49 0.49 0.48 0.47 0.48 0.49 0.49 0.52 0.52 0.54 0.54 0.51 0.47 0.47 0.48
Median: 0.21 0.29 0.34 0.36 0.41 0.47 0.47 0.51 0.52 0.54 0.54 0.55 0.55 0.55 0.56 0.55 0.51 0.52 0.56 0.57 0.58 0.58 0.58 0.54 0.55 0.59 0.63 0.63 0.62 0.58 0.58 0.60
Pearson correlation
Predictions
Control
Genomic distance, kb

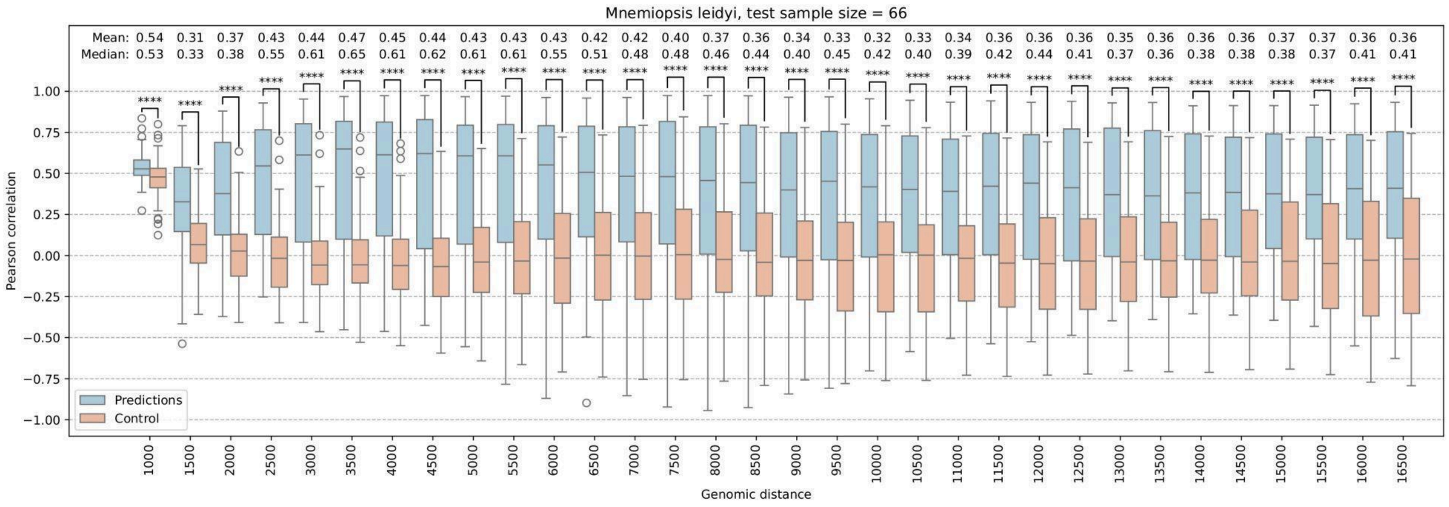
Mnemiopsis leidyi, test sample size = 66
Mean: 0.54 0.31 0.37 0.43 0.44 0.47 0.45 0.44 0.43 0.43 0.43 0.42 0.42 0.40 0.37 0.36 0.34 0.33 0.32 0.33 0.34 0.36 0.36 0.36 0.35 0.36 0.36 0.36 0.37 0.37 0.36 0.36
Median: 0.53 0.33 0.38 0.55 0.61 0.65 0.61 0.62 0.61 0.61 0.55 0.51 0.48 0.48 0.46 0.44 0.40 0.45 0.42 0.40 0.39 0.42 0.44 0.41 0.37 0.36 0.38 0.38 0.38 0.37 0.41 0.41
Pearson correlation
Predictions
Control
Genomic distance

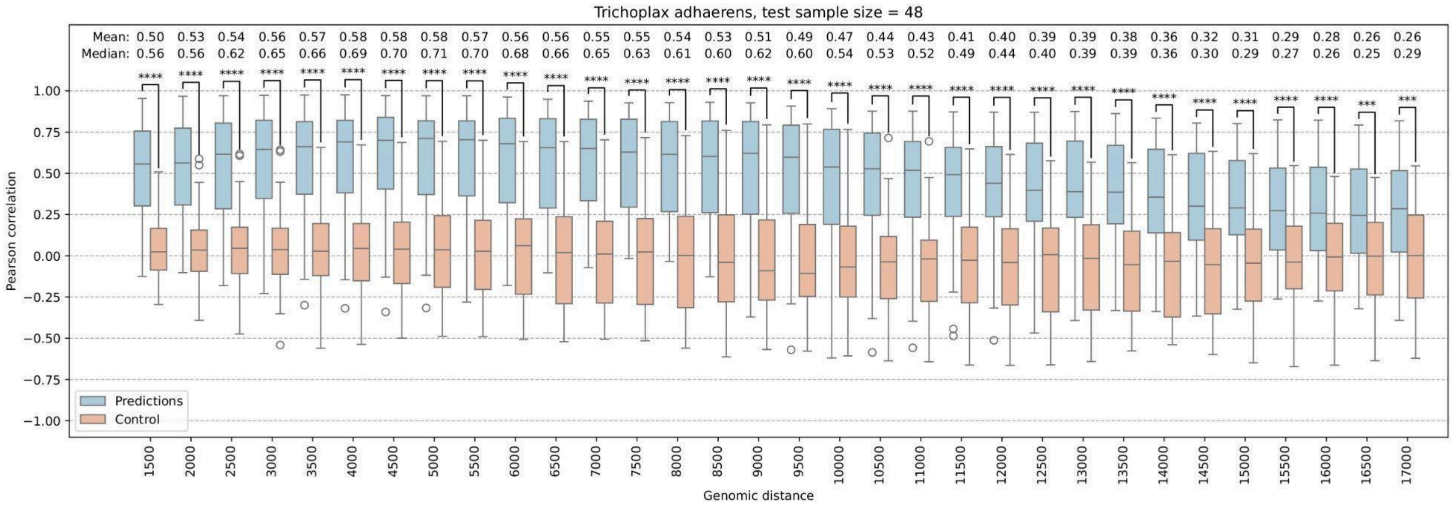
Trichoplax adhaerens, test sample size = 48
Mean: 0.50 0.53 0.54 0.56 0.57 0.58 0.58 0.58 0.57 0.56 0.56 0.55 0.55 0.54 0.53 0.51 0.49 0.47 0.44 0.43 0.41 0.40 0.39 0.39 0.38 0.36 0.32 0.31 0.29 0.28 0.26 0.26
Median: 0.56 0.56 0.62 0.65 0.66 0.69 0.70 0.71 0.70 0.68 0.66 0.65 0.63 0.61 0.60 0.62 0.60 0.54 0.53 0.52 0.49 0.44 0.40 0.39 0.39 0.36 0.30 0.29 0.27 0.26 0.25 0.29
Pearson correlation
Predictions
Control
Genomic distance
1500 2000 2500 3000 3500 4000 4500 5000 5500 6000 6500 7000 7500 8000 8500 9000 9500 10000 10500 11000 11500 12000 12500 13000 13500 14000 14500 15000 15500 16000 16500 17000

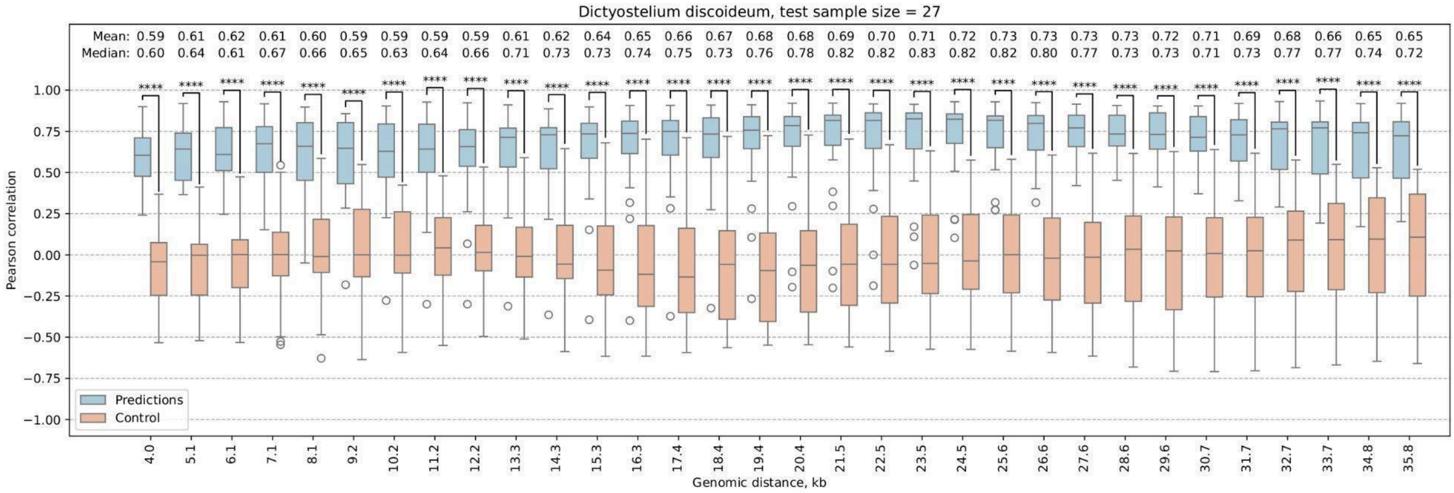
Dictyostelium discoideum, test sample size = 27
Mean: 0.59 0.61 0.62 0.61 0.60 0.59 0.59 0.59 0.59 0.61 0.62 0.64 0.65 0.66 0.67 0.68 0.68 0.69 0.70 0.71 0.72 0.73 0.73 0.73 0.73 0.72 0.71 0.69 0.68 0.66 0.65 0.65
Median: 0.60 0.64 0.61 0.67 0.66 0.65 0.63 0.64 0.66 0.71 0.73 0.73 0.74 0.75 0.73 0.76 0.78 0.82 0.82 0.83 0.82 0.82 0.80 0.77 0.73 0.73 0.71 0.73 0.77 0.77 0.74 0.72
Pearson correlation
Predictions
Control
Genomic distance, kb
4.0 5.1 6.1 7.1 8.1 9.2 10.2 11.2 12.2 13.3 14.3 15.3 16.3 17.4 18.4 19.4 20.4 21.5 22.5 23.5 24.5 25.6 26.6 27.6 28.6 29.6 30.7 31.7 32.7 33.7 34.8 35.8

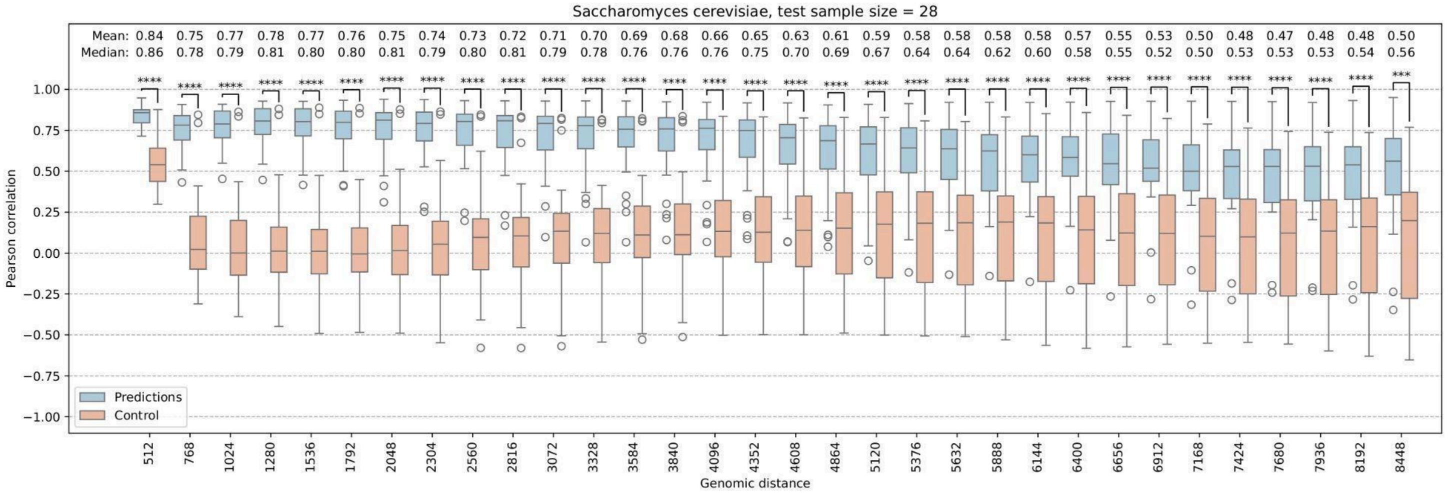
Saccharomyces cerevisiae, test sample size = 28
Mean: 0.84 0.75 0.77 0.78 0.77 0.76 0.75 0.74 0.73 0.72 0.71 0.70 0.69 0.68 0.66 0.65 0.63 0.61 0.59 0.58 0.58 0.58 0.58 0.57 0.55 0.53 0.50 0.48 0.47 0.48 0.48 0.50
Median: 0.86 0.78 0.79 0.81 0.80 0.80 0.81 0.79 0.80 0.81 0.79 0.78 0.76 0.76 0.76 0.75 0.70 0.69 0.67 0.64 0.64 0.62 0.60 0.58 0.55 0.52 0.50 0.53 0.53 0.53 0.54 0.56
Pearson correlation
Predictions
Control
Genomic distance
512 768 1024 1280 1536 1792 2048 2304 2560 2816 3072 3328 3584 3840 4096 4352 4608 4864 5120 5376 5632 5888 6144 6400 6656 6912 7168 7424 7680 7936 8192 8448

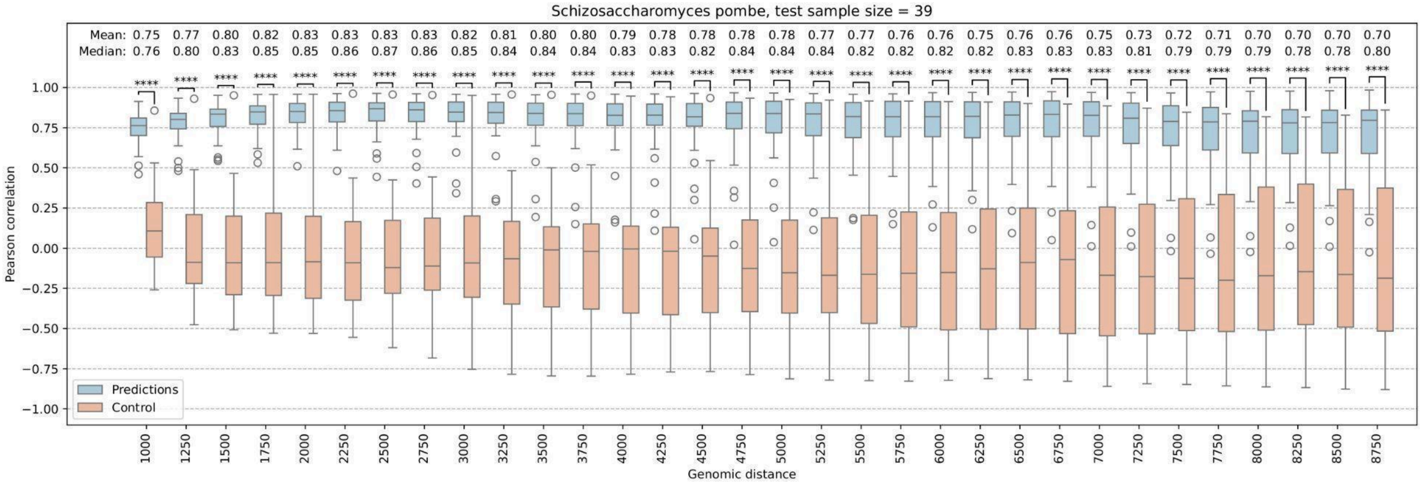


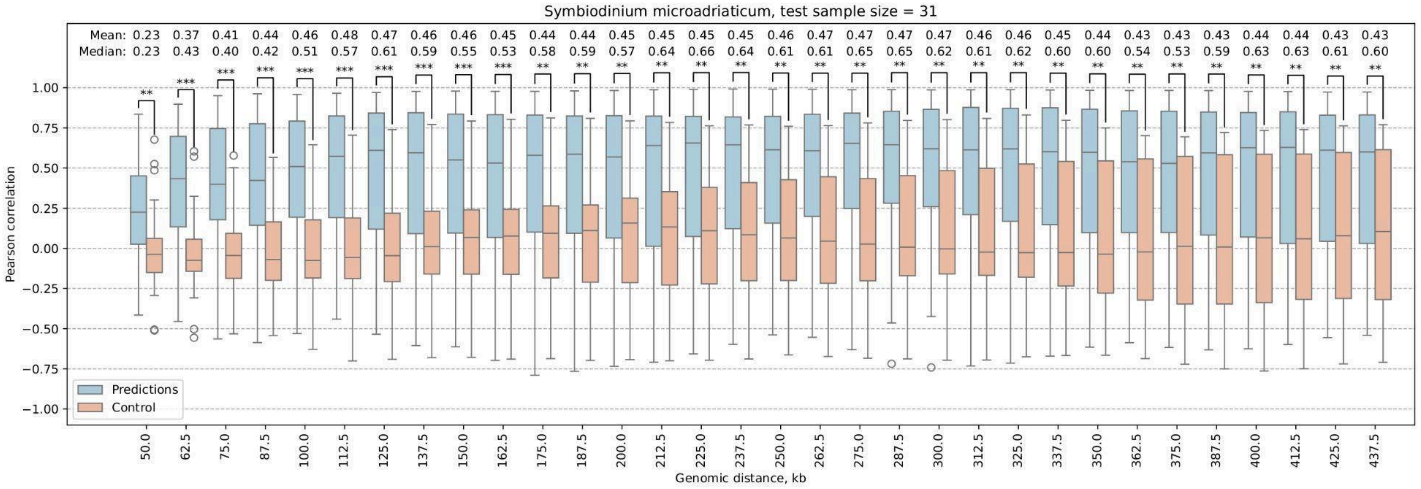


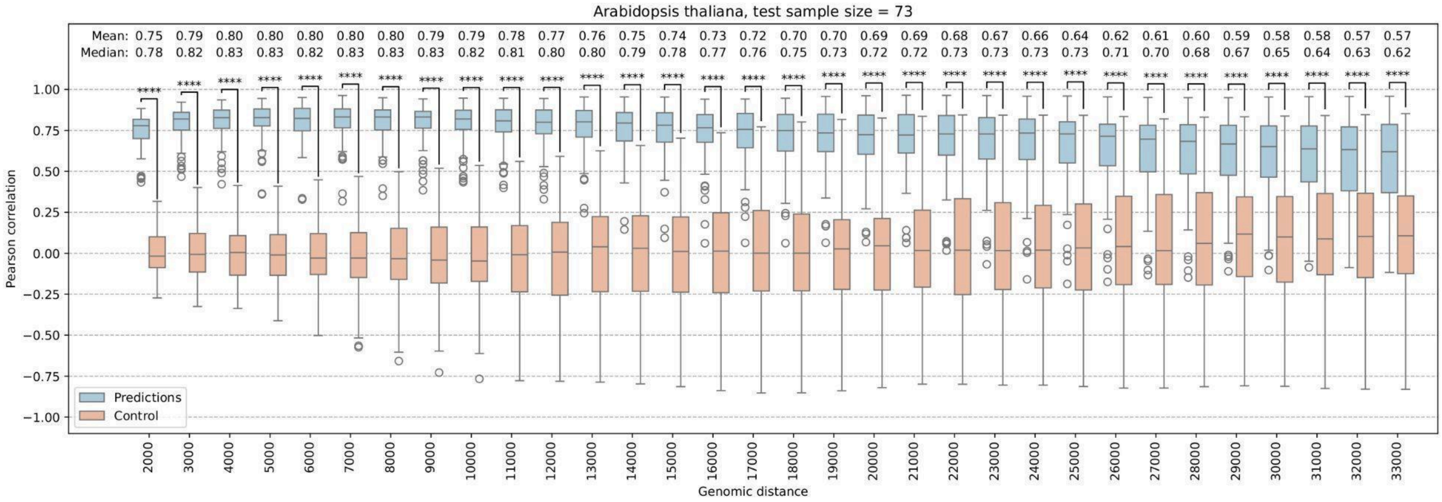


**Supplementary Figure 6. Detailed metrics for predictions on test samples.**

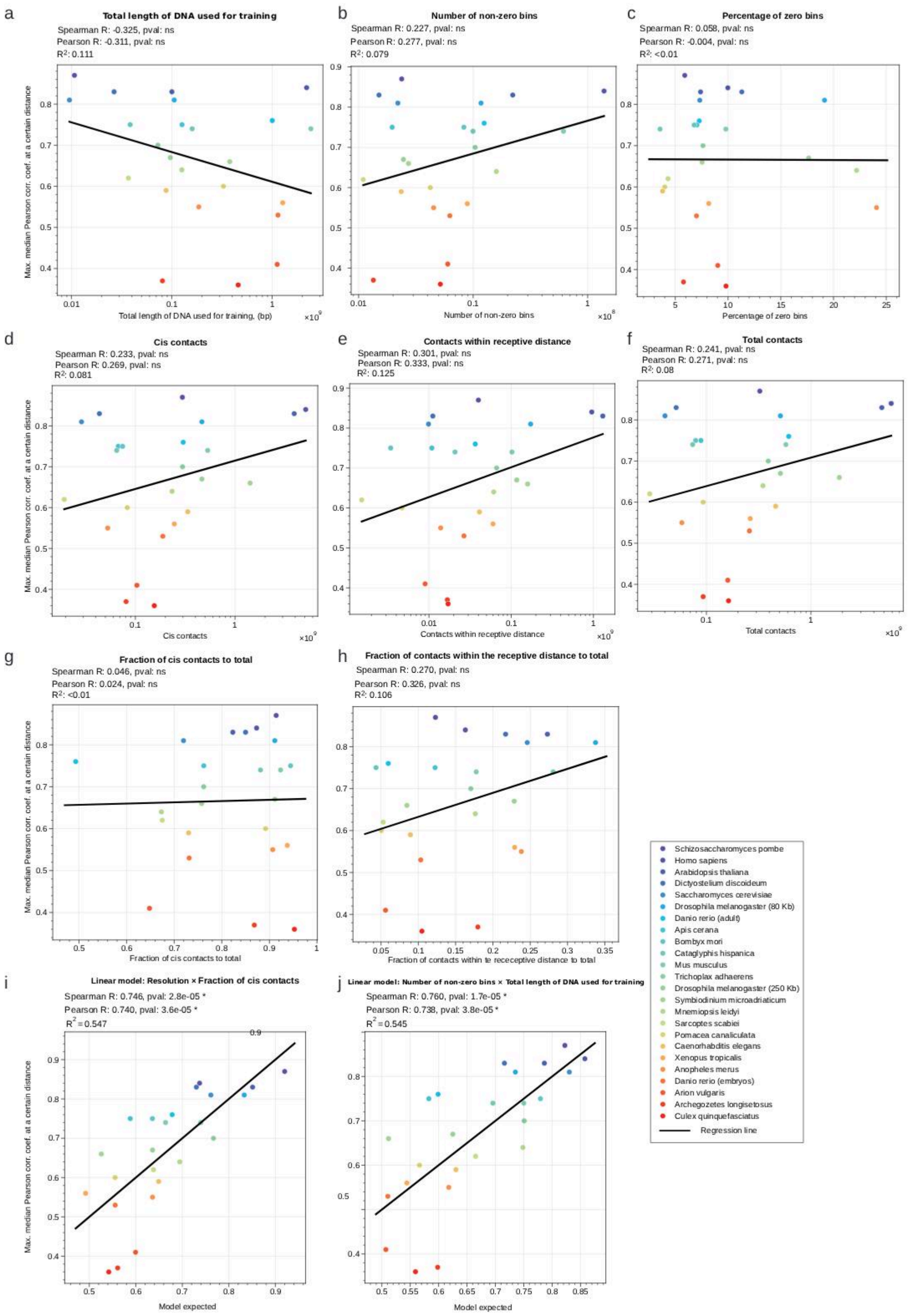


**Supplementary Figure 7.**

**The Chimaera performance depends on the quality of the input Hi-C/Micro-C data.** Each dot is a Chimaera model trained on a single species (except two window sizes studied for *Drosophila melanogaster*, and two Hi-C sources for *Danio rerio*, embryo and adult muscle tissues). The black line represents the regression line. All *p*-values were corrected for multiple testing by the Bonferroni correction (separately for the Spearman and Pearson correlations). ns, non-significant; *, significant ($p<0.01$).

**(A–H)** None of the input data characteristics explain Chimaera model performance.

**(I)** A linear combination of two characteristics—Hi-C/Micro-C resolution and cis-to-total contact ratio—explains 0.547 of the observed variance, which is the best among any single characteristic or pairwise combination.

**(J)** The second-best model (a combination of the number of non-zero bins and the genome size in the training set) explains 0.545 of the observed variance.

*Hi-C/Micro-C quality control.* The counts of Hi-C/Micro-C contacts (before iterative correction) and bins were calculated using the cooler library (37). For counts within the receptive-distance, we excluded contacts on the first two diagonals of the Hi-C matrix. Bins excluded by iterative correction (default parameters) were treated as zero bins; all remaining bins were considered non-zero.

*Relating Hi-C/Micro-C quality to model performance.* For each species, we took the maximum Pearson correlation observed across diagonals (the matrix diagonal with the highest correlation) and fitted linear ordinary least squares (OLS) models regressing this value on individual Hi-C/Micro-C quality metrics. Regressions were implemented in Python using statsmodels (38). Log-transformed predictors and two-predictor OLS models using all pairs of characteristics (original or log-transformed) were probed and selected based on the highest coefficient of determination ($R^2$). Each two-predictor model included an intercept, individual terms, and their combination. The models in (I) and (J) used log-transformed characteristics.

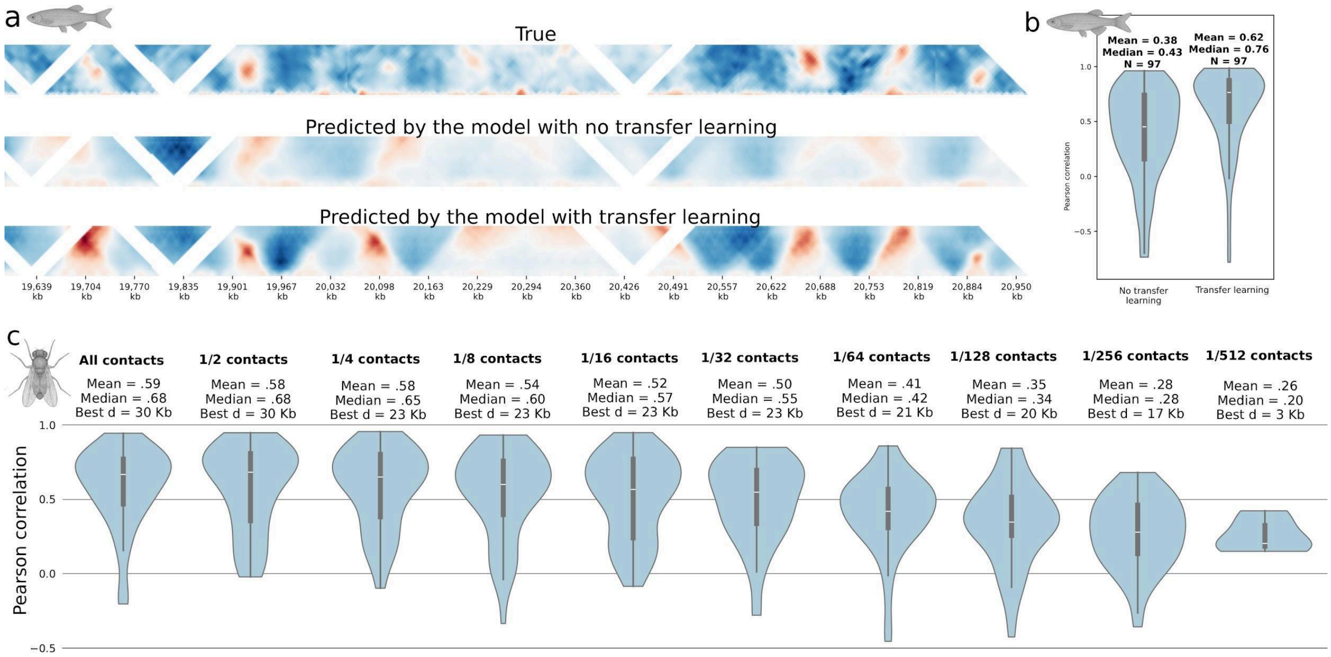


**Supplementary Figure 8.**

**Testing the model quality.** Transfer learning for model improvement for *Danio rerio*. The Hi-C data are for *D. rerio* adult muscle from (23). Pre-training was done on human HFFc6 Micro-C data (20).

**(A)** Hi-C maps for a test genomic region of *D. rerio*: top – true data, middle – predicted by a model without transfer learning, bottom – with transfer learning, where the model was first trained on the human data.

**(B)** The Pearson correlation between the original maps and the predicted by the models for *D. rerio* trained without (left) and with transfer learning (right). The test sample results are shown.

**(C)** Training the model on downsampled datasets. The quality of models trained on low-quality *Drosophila melanogaster* contact maps was assessed by the maximum Pearson correlation coefficient between predicted and true contacts. Low data quality was simulated by randomly removing contacts; the remaining fraction is specified above each plot. The value 'Best d' denotes the optimal distance at which the reported correlation was calculated. The trend is the decrease in 'Best d' corresponding to the reduction in training contacts. The window size is 100 Kb.

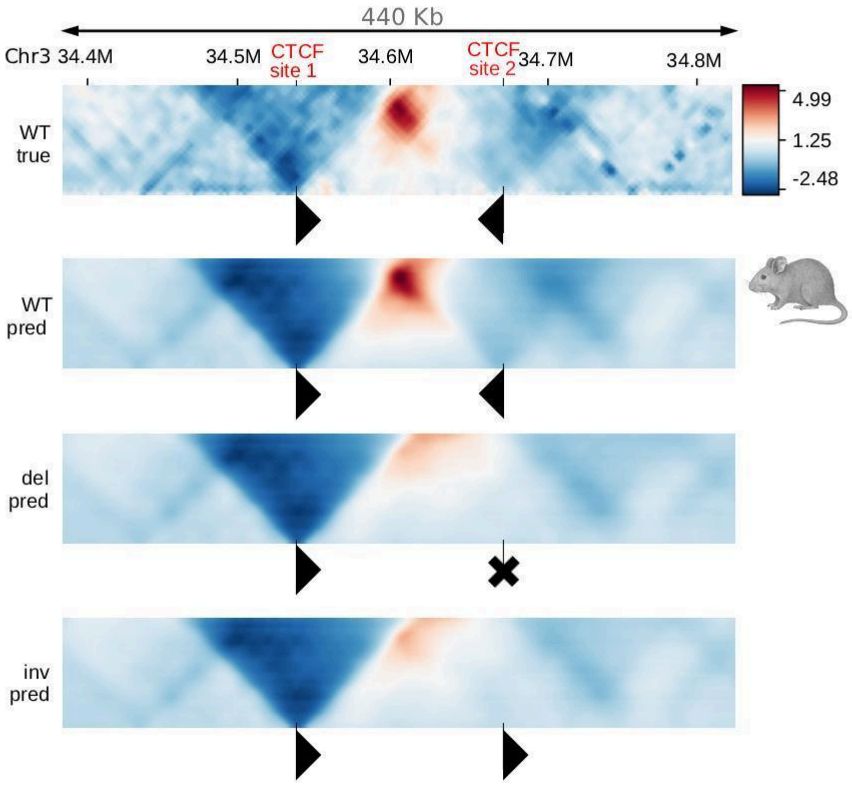


**Supplementary Figure 9.**

**Chimaera reproduces the results of *in vivo* CTCF site removal and inversion.** Region of chr3 of mouse with two strong convergent CTCF binding sites from de Wit et al. (39). In wild type cells, two convergent CTCFs form a strong loop both in real Hi-C maps (WT true) and in the Chimaera prediction (WT pred). After CTCF removal, the strong loop disappears in real Hi-C maps (39), which is well recapitulated by Chimaera (del pred). Inversion of the right inwards facing CTCF, results in the same effect both in real Hi-C (39) and in Chimaera prediction (inv pred).

Black triangles indicate CTCF site orientations (the vertical side is anchored to site position), cross indicates site removal.

WT true – actual Hi-C map, WT pred – a map predicted from the reference sequence, del pred – a map predicted from the sequence with removed right CTCF site, inv pred – predicted from the sequence with the right CTCF site inversion.

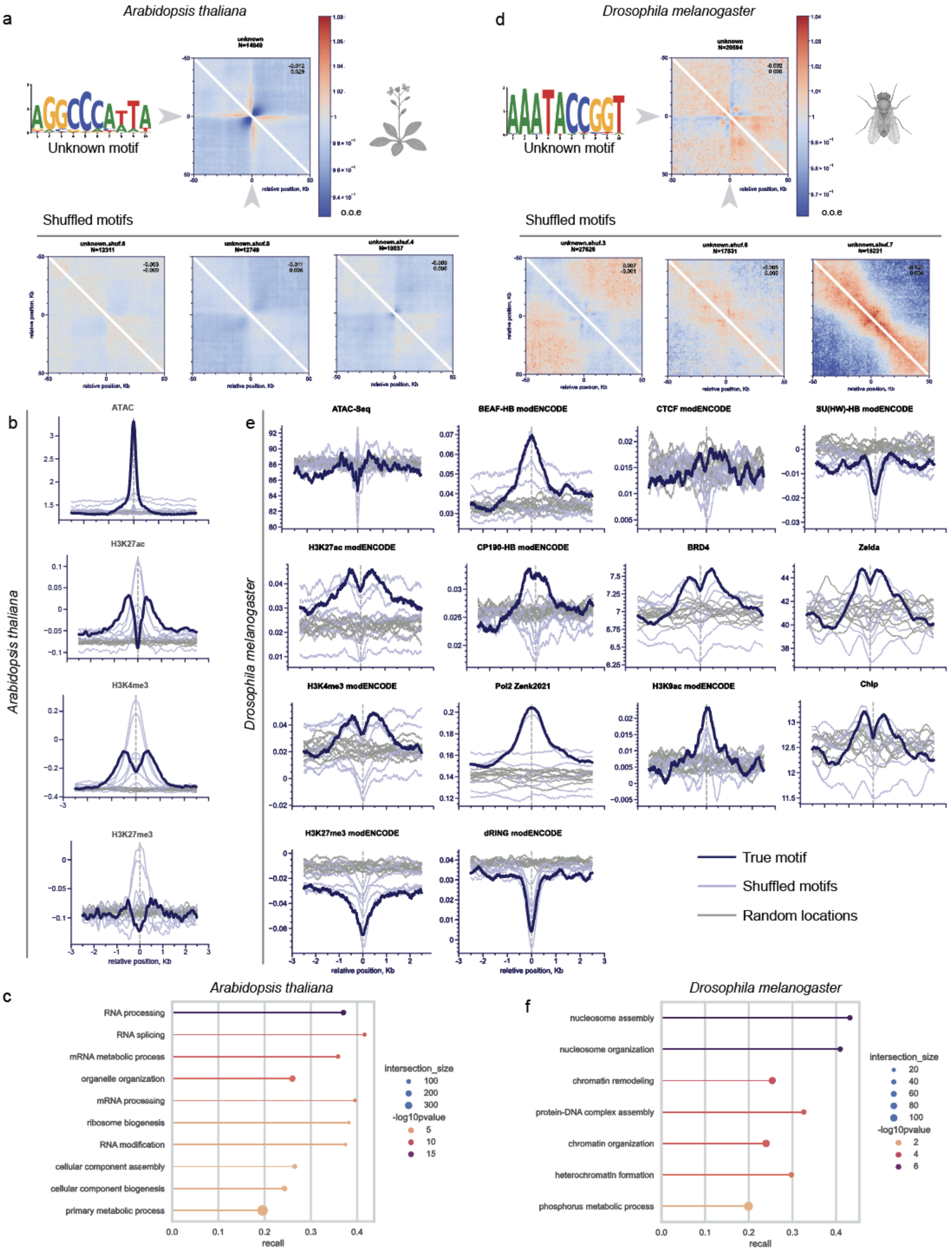


Supplementary Figure 10.

Properties of novel discovered motifs.

**(A)** Pileups of Micro-C data at the AGGCCCATTA motif sites in *Arabidopsis thaliana* (top) and controls with shuffled motifs (bottom).

**(B)** Pileups of ATAC-Seq and ChIP-Seqs at the AGGCCCATTA motif sites in *A. thaliana*. ATAC-Seq for shoots from (40) (GSE116287); histone marks from (41) (GSE120664).
**(C)** Enriched Gene Ontology (GO) terms of the genes that contain the AGGCCCATTA motif sites closer than 500 bp to their TSS.

**(D)** Pileups of Micro-C data at the AAATACCGGT motif sites in *Drosophila melanogaster* (top) and controls with shuffled motifs (bottom).

**(E)** Pileups of ATAC-Seq and ChIP-Seqs at the AAATACCGGT motif sites in *D. melanogaster*. ATAC-Seq from; histone marks, insulators and dRING from modENCODE (42); Pol2 ChIP-Seq from (43) (GSE140539); BRD4, Zelda and Chip are from (44) (PRJEB88447).

**(F)** Enriched Gene Ontology terms of the genes that contain the AAATACCGGT motif sites closer than 500 bp to their TSS.

Instances of newly discovered motifs in *D. melanogaster* and *A. thaliana* were called by whole-genome PWMScan-based (45) motif scanner via the JASPAR-UCSC-tracks pipeline (46); sites meeting the thresholds $p$ < 1×10ˆ-4 and $r$ > 0.8 thresholds were retained.

**(A, D)** Micro-C pileups were built with the cooltools *pileup* function (19).

**(B, E)** Epigenetics pileups were built with pybbi (47), a wrapper around the Big Binary Indexed file (BBI) library (48).

**(C, F)** For GO enrichment, we took all the genes of target species that have a called motif within 500 bps to its transcription start site. We then performed Gene Ontology enrichment with g:Profiler (49). We tested the GO Biological Process; *p*-values were adjusted by the Benjamini–Hochberg FDR.

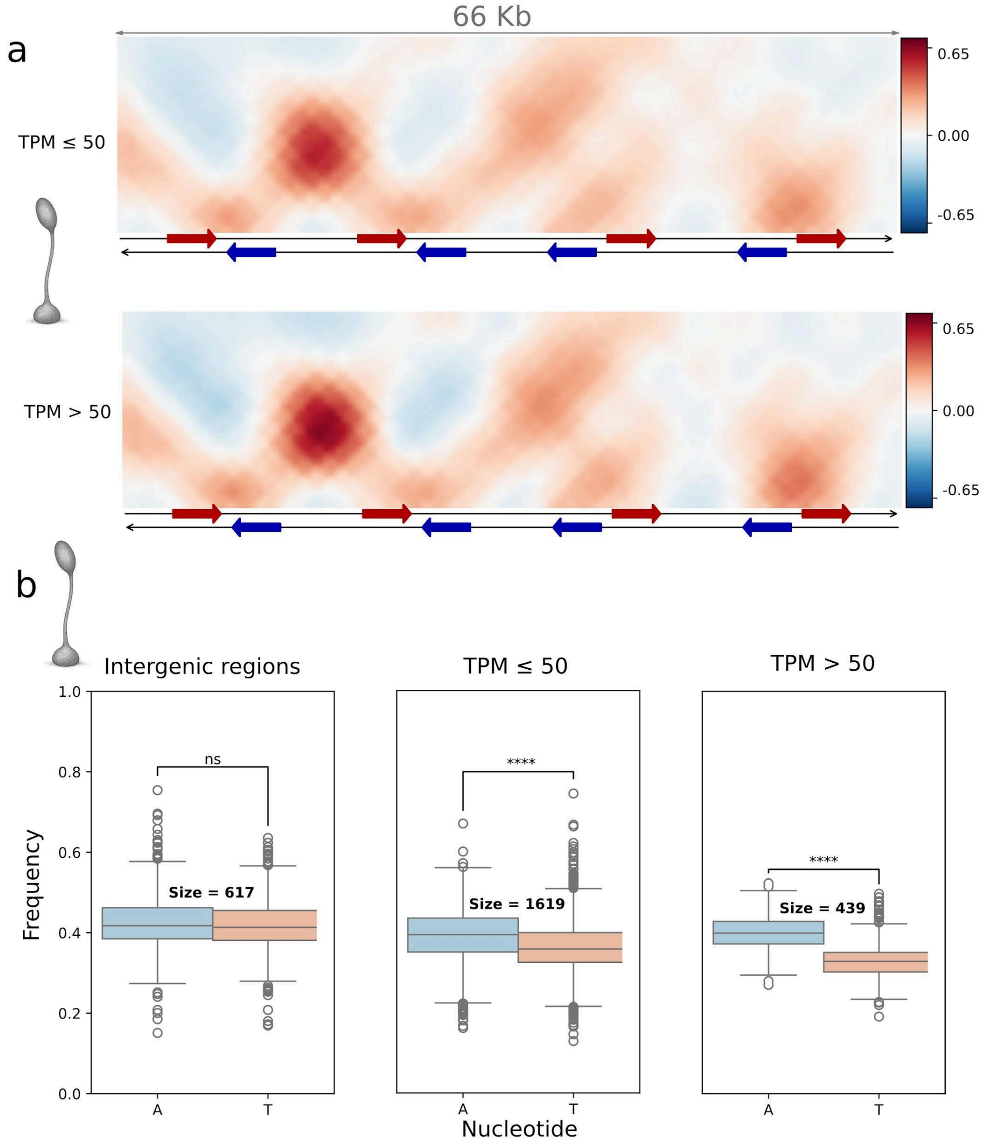


**Supplementary Figure 11.**

**Gene expression in *Dictyostelium discoideum* affects model predictions.** Properties of genes with different levels of expression at the vegetative state of *D. discoideum* development (32): Chimaera predictions (A) and nucleotide content (B). TPM – transcripts per million. **(A)** *In silico* generated sequence with genes with low (TPM<50, top) and high (TPM>50, bottom) expression level. The loops and other patterns are more pronounced for the highly expressed genes, although the model was not informed on the expression level. **(B)** Testing the hypothesis that nucleotide content is different in genomic regions with different expression properties: A and T nucleotide frequencies at intergenic regions and at the genes with low and high expression levels.

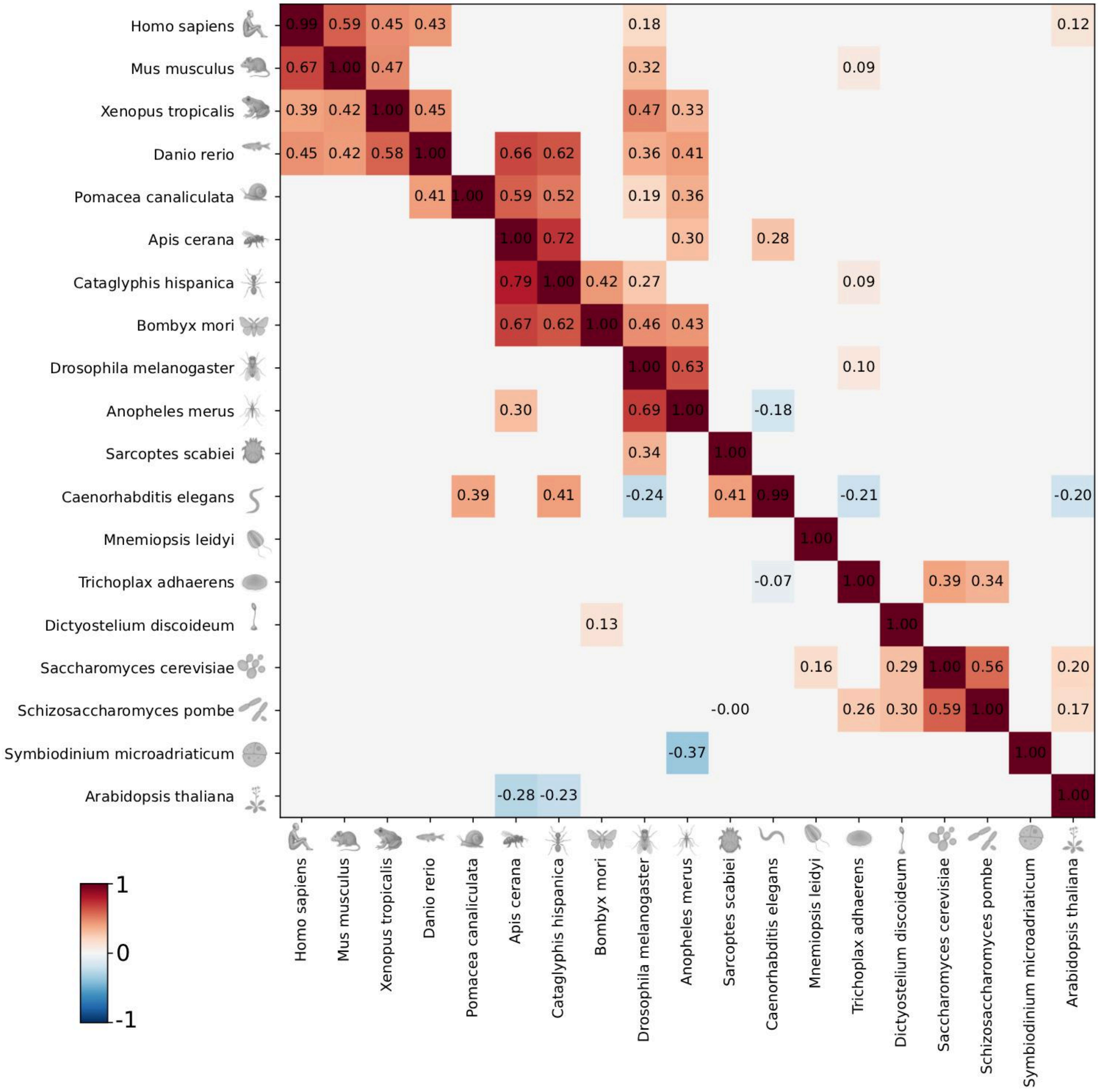


**Supplementary Figure 12.**

Supporting information for Figure 8. Correlations between Hi-C maps predicted by a model trained on a given species (columns) and Hi-C maps predicted by models trained for different species (rows). Empty squares – correlations that are not statistically significant.

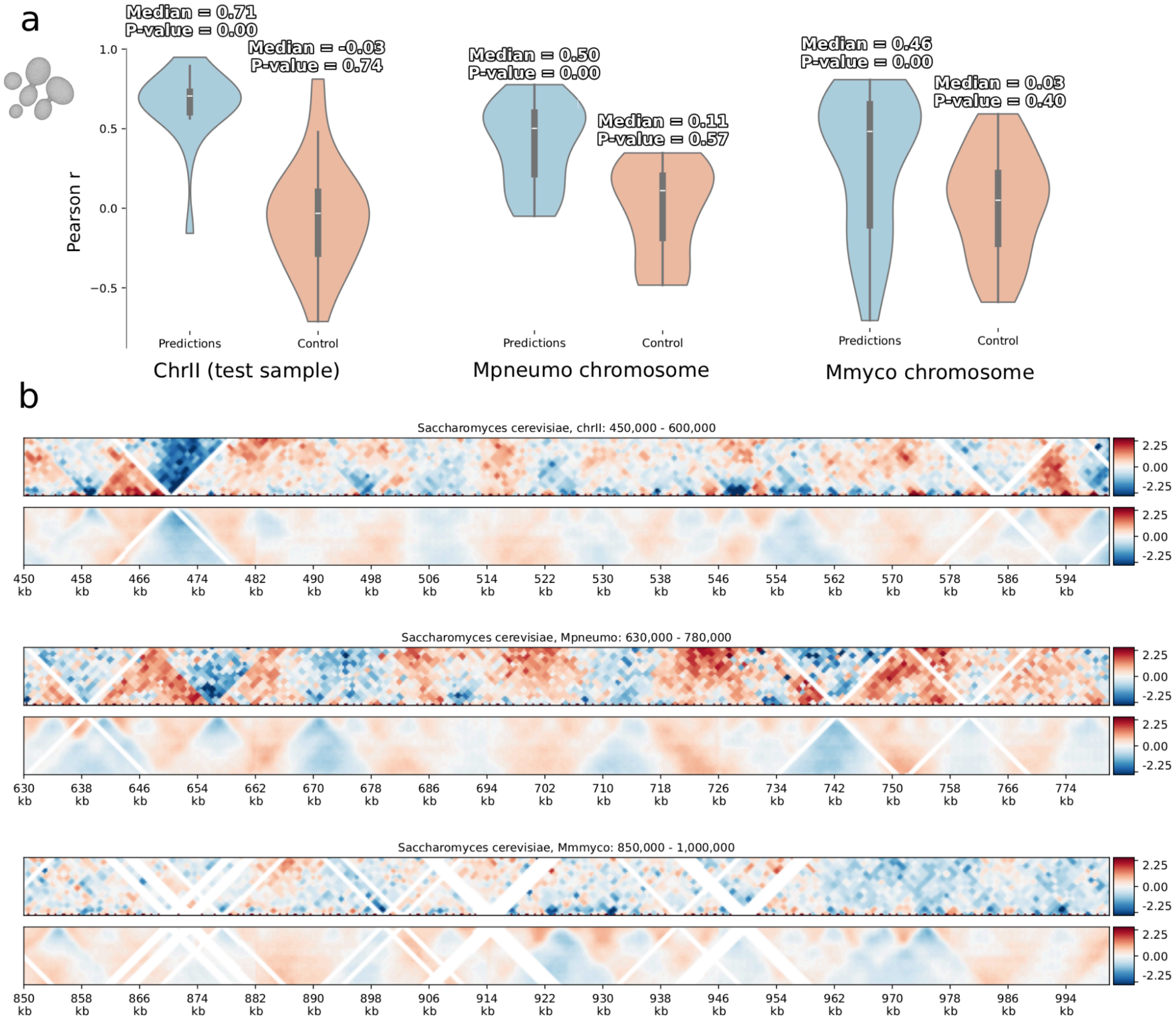


**Supplementary Figure 13.**

**Chimaera performance on exogenous DNA in *S. cerevisiae*.**

Hi-C data of *S. cerevisiae* genome, *Mycoplasma pneumoniae* (Mpneumo) and *Mycoplasma mycoides* (Mmyco) genomes after introduction into yeast nuclei from (50).

Meneu *et al. (50)* integrate megabase-scale foreign *Mycoplasma* chromosomes into budding yeast and demonstrate that the *M. pneumoniae* chromosome behaves similar to *S. cerevisiae*, while *M. mycoides* chromosome is less active. Here, we launch yeast-trained Chimaera on *Mycoplasma* DNA and demonstrate its ability to predict the genome organization of exogenous DNA in a given cellular context. The Chimaera model was trained on *S. cerevisiae*, window size 32 Kb. Neither Chr II nor *Mycoplasma* data were used for training.

**(A)** The Pearson correlation coefficient between the predicted and experimental contacts at the 9 Kb distance for yeast chromosome chrII and two exogenous chromosomes with bacterial DNA (Mpneumo and Mmyco). The controls are shuffled pairs of true and predicted map windows. The *p*-values are shown for the Mann-Whitney test with H0 that the median is 0.

**(B)** Examples of predictions for three chromosomes. The top half of each plot is the true map from the Meneu *et al.* data, the bottom half is the mirrored predicted map. Note that the Hi-C data is corrected by expected values from scaling, thus the global effect of inactivation and compactization in *M. mycoides (50)* is not prominent.

| 1 | 2 | 3 | 4 | 5 | 6 | 7 | 8 | 9 | 10 | 11 | 12 | 13 | 14 | 15 | 16 | 17 | 18 | 19 | 20 | 21 | 22 | 23 |
|---|---|---|---|---|---|---|---|---|---|---|---|---|---|---|---|---|---|---|---|---|---|---|
| Species | Assembly name | Genome size (approx.), Mbp | Hi-C snippet (window) size, Kbp | Original resolution, Kbp | Train sample size | Total length of DNA used for training, Mbp | Number of non-zero bins, millions | Fraction of zero bins | Contacts within the receptive distance of the model, millions | *Cis* contacts, millions | Total contacts, millions | Fraction of contacts within the receptive distance to total | Fraction of *cis* contacts | Organism for DNA encoder transfer | Organism for Hi-C autoencoder transfer | Median Pearson correlation coefficient of full true and predicted maps | Maximum median Pearson correlation coefficient of map contacts at a certain distance | Distance at which the correlation maximum is achieved (approx.), Kbp | Loop prediction quality | Insulation prediction quality | Fountain prediction quality | TAD prediction quality |
| *Homo sapiens* | hg38 | 3100 | 262,14 | 2 | 8424 | 2208,30 | 139,00 | 9,98 | 954,84 | 5112,22 | 5855,24 | 0,16 | 0,87 | | | 0,79 | 0,84 | 140 | 0,60 | 0,79 | 0,71 | 0,71 |
| *Mus musculus* | mm10 | 2700 | 262,14 | 4 | 9287 | 2434,53 | 61,47 | 9,79 | 102,23 | 531,44 | 575,48 | 0,18 | 0,92 | | | 0,68 | 0,74 | 135 | 0,58 | 0,70 | 0,64 | 0,77 |
| *Xenopus tropicalis* | xenTro10 | 1500 | 405,00 | 15 | 3151 | 1276,16 | 8,89 | 8,17 | 60,14 | 246,08 | 262,47 | 0,23 | 0,94 | | | 0,44 | 0,56 | 110 | 0,28 | 0,67 | 0,39 | 0,34 |
| *Danio rerio* (embryos) | danRer11 | 1400 | 400,00 | 20 | 2861 | 1144,40 | 6,26 | 7,01 | 26,64 | 188,67 | 257,95 | 0,10 | 0,73 | | *H. sapiens* | 0,49 | 0,53 | 120 | 0,36 | 0,46 | 0,40 | 0,44 |
| *Danio rerio* (adult) | danRer11 | 1400 | 262,14 | 10 | 3834 | 1005,06 | 12,47 | 7,27 | 36,56 | 302,04 | 612,45 | 0,06 | 0,49 | *H. sapiens* | | 0,69 | 0,76 | 135 | 0,55 | 0,72 | 0,64 | 0,56 |
| *Arion vulgaris* | ASM2079622v1 | 1500 | 500,00 | 25 | 2247 | 1123,50 | 6,00 | 9,03 | 8,97 | 103,77 | 160,10 | 0,06 | 0,65 | | *D. melanogaster* | 0,44 | 0,41 | 120 | 0,18 | 0,29 | ns | 0,42 |
| *Pomacea canaliculata* | ASM307304v1 | 440 | 262,14 | 10 | 1248 | 327,16 | 4,23 | 4,00 | 4,70 | 83,01 | 93,06 | 0,05 | 0,89 | | *H. sapiens* | 0,48 | 0,6 | 130 | 0,34 | 0,59 | 0,30 | 0,13 |
| *Apis cerana* | ASM1110058v1 | 215 | 262,14 | 10 | 483 | 126,62 | 1,96 | 7,05 | 10,84 | 67,47 | 88,53 | 0,12 | 0,76 | | *D. melanogaster* | 0,71 | 0,75 | 70 | 0,52 | 0,79 | 0,60 | 0,68 |
| *Cataglyphis hispanica* | ULB_Chis1_1.0 | 206 | 256,00 | 2 | 624 | 159,74 | 9,97 | 3,55 | 20,72 | 65,04 | 73,80 | 0,28 | 0,88 | | | 0,69 | 0,74 | 65 | 0,66 | 0,81 | 0,68 | 0,77 |
| *Bombyx mori* | Bmori_2016v1.0 | 463 | 25,00 | 5 | 1540 | 38,50 | 8,30 | 6,79 | 3,42 | 74,54 | 78,93 | 0,04 | 0,94 | | | 0,69 | 0,75 | 130 | 0,60 | 0,70 | 0,51 | 0,57 |
| *Drosophila melanogaster* (80 Kb) | dm6 | 175 | 80,00 | 1 | 1322 | 105,76 | 11,72 | 19,15 | 172,26 | 465,10 | 510,35 | 0,34 | 0,91 | | | 0,76 | 0,81 | 40 | 0,66 | 0,74 | 0,71 | 0,76 |
| *Drosophila melanogaster* (250 Kb) | dm6 | 175 | 250,00 | 5 | 387 | 96,75 | 2,46 | 17,64 | 116,64 | 465,10 | 510,35 | 0,23 | 0,91 | | | 0,65 | 0,67 | 40 | 0,54 | 0,76 | 0,56 | 0,41 |
| *Anopheles merus* | AmerM5.1 | 294 | 200,00 | 5 | 930 | 186,00 | 4,52 | 24,08 | 13,86 | 52,80 | 58,22 | 0,24 | 0,91 | | | 0,48 | 0,55 | 70 | 0,25 | 0,52 | 0,42 | 0,30 |
| *Culex quinquefasciatus* | VPISU_Cqui_1.0 | 573 | 262,14 | 10 | 1752 | 459,28 | 5,17 | 9,81 | 17,06 | 154,95 | 162,68 | 0,10 | 0,95 | | *D. melanogaster* | 0,5 | 0,36 | 75 | 0,38 | 0,50 | 0,48 | 0,52 |
| *Sarcoptes scabiei* | ASM2084414v1 | 57 | 131,07 | 5 | 282 | 36,96 | 1,10 | 4,31 | 1,52 | 19,36 | 28,68 | 0,05 | 0,68 | | | 0,6 | 0,62 | 25 | 0,33 | 0,45 | 0,57 | 0,47 |
| *Archegozetes longisetosus* | Caltech_Along_2.0 | 141 | 262,14 | 10 | 308 | 80,74 | 1,35 | 5,78 | 16,74 | 80,80 | 93,05 | 0,18 | 0,87 | | | 0,41 | 0,37 | 80 | ns | 0,36 | 0,21 | 0,51 |
| *Caenorhabditis elegans* | ce10 | 100 | 131,07 | 4 | 670 | 87,82 | 2,36 | 3,80 | 41,08 | 335,71 | 459,68 | 0,09 | 0,73 | | | 0,53 | 0,59 | 35 | 0,29 | 0,59 | 0,36 | 0,62 |
| *Mnemiopsis leidyi* | crg_Mlei_v2 | 208 | 32,00 | 1 | 3939 | 126,05 | 15,96 | 22,19 | 61,25 | 233,79 | 347,32 | 0,18 | 0,67 | | | 0,53 | 0,64 | 5 | 0,21 | 0,38 | 0,18 | 0,26 |
| *Trichoplax adhaerens* | ASM15027v1 | 105 | 32,00 | 1 | 2276 | 72,83 | 10,41 | 7,63 | 66,50 | 297,30 | 390,20 | 0,17 | 0,76 | | | 0,57 | 0,7 | 5 | 0,40 | 0,43 | 0,45 | 0,67 |
| *Saccharomyces cerevisiae* | sacCer3 | 12 | 16,38 | 0,5 | 583 | 9,55 | 2,20 | 7,32 | 9,87 | 28,87 | 40,12 | 0,25 | 0,72 | | | 0,72 | 0,81 | 2 | 0,58 | 0,58 | 0,60 | 0,70 |
| *Schizosaccharomyces pombe* | ASM294v2 | 12 | 16,00 | 0,5 | 670 | 10,72 | 2,38 | 5,90 | 39,87 | 296,41 | 324,24 | 0,12 | 0,91 | | | 0,86 | 0,87 | 2 | 0,67 | 0,81 | 0,85 | 0,83 |
| *Dictyostelium discoideum* | dicty_2.7 | 33 | 65,54 | 2 | 406 | 26,61 | 1,51 | 11,31 | 11,14 | 43,61 | 51,31 | 0,22 | 0,85 | | | 0,79 | 0,83 | 25 | 0,73 | 0,81 | 0,76 | 0,81 |
| *Symbiodinium microadriaticum* | Smic1.1 | 692 | 800,00 | 25 | 473 | 378,40 | 2,73 | 7,54 | 158,30 | 1415,14 | 1867,78 | 0,08 | 0,76 | | | 0,7 | 0,66 | 275 | 0,60 | 0,74 | 0,26 | 0,68 |
| *Arabidopsis thaliana* | TAIR10 | 119 | 64,00 | 0,5 | 1568 | 100,35 | 22,16 | 7,39 | 1300,62 | 3920,76 | 4763,17 | 0,27 | 0,82 | | | 0,79 | 0,83 | 10 | 0,70 | 0,73 | 0,78 | 0,77 |

**Supplementary Table 1.**
**Input data properties and the Chimaera models metrics on test samples.**

| Model variant | Model | *S. cerevisiae* | *H. sapiens* |
|---|---|---|---|
| 1 | Adapted Akita (modified for rotated maps) | 0.49 | **0.48** |
| 2 | Chimaera with Akita head | 0.50 | 0.44 |
| 3 | Chimaera with Hi-C-decoder trained on other organism data | 0.56 | **0.48** |
| 4 | Chimaera model omitting pre-training of Hi-C-decoder | 0.47 | 0.41 |
| 5 | Chimaera with pre-trained Hi-C-decoder trained to predict raw maps | 0.52 | 0.42 |
| 6 | Chimaera with pre-trained Hi-C-decoder trained to predict denoised maps | **0.59** | **0.49** |

**Supplementary Table 2.** The Pearson correlation between true non-denoised and predicted maps from the validation samples after a fixed number of training epochs for different versions of the models for the *S. cerevisiae* and *H. sapiens* data. Model variant #4 is a control that demonstrates the effectiveness of pre-training of the Hi-C decoder prior to the DNA encoder (see Discussion). Model variant #5 is a control that tests prediction of raw maps instead of denoised maps, which works slightly worse potentially due to contribution of unlearnable noise in the data (see Discussion). Model # 6 is the final variant used throughout the paper.

46. JASPAR-UCSC-tracks: Code and data used to create the JASPAR UCSC Genome Browser tracks data hub Github.

47. Abdennur,N. pybbi: Python bindings to UCSC BigWig and BigBed library Github.

48. Kent,W.J., Zweig,A.S., Barber,G., Hinrichs,A.S. and Karolchik,D. (2010) BigWig and BigBed: enabling browsing of large distributed datasets. *Bioinformatics*, **26**, 2204–2207.

49. Raudvere,U., Kolberg,L., Kuzmin,I., Arak,T., Adler,P., Peterson,H. and Vilo,J. (2019) g:Profiler: a web server for functional enrichment analysis and conversions of gene lists (2019 update). *Nucleic Acids Res.*, **47**, W191–W198.

50. Meneu,L., Chapard,C., Serizay,J., Westbrook,A., Routhier,E., Ruault,M., Perrot,M., Minakakis,A., Girard,F., Bignaud,A., *et al.* (2025) Sequence-dependent activity and compartmentalization of foreign DNA in a eukaryotic nucleus. *Science*, **387**, eadm9466.
